# An extensively optimized chromatin immunoprecipitation protocol for quantitatively comparable and robust results

**DOI:** 10.1101/835926

**Authors:** Wim J. de Jonge, Mariël Brok, Patrick Kemmeren, Frank C.P. Holstege

**Affiliations:** Princess Máxima Center for Pediatric Oncology, Heidelberglaan 25, 3584 CS Utrecht, the Netherlands

## Abstract

Chromatin immunoprecipitation (ChIP) is a commonly used technique to investigate which parts of a genome are bound by a particular protein. The result of ChIP is often interpreted in a binary manner: bound or not bound. Due to this focus, ChIP protocols frequently lack the ability to quantitatively compare samples with each other, for example in a time series or under different growth conditions. Here, using the yeast *S. cerevisiae* transcription factors Cbf1, Abf1, Reb1, Mcm1 and Sum1, we optimized the five major steps of a commonly used ChIP protocol: cross-linking, quenching, cell lysis, fragmentation and immunoprecipitation. Quenching with glycine is inefficient and can lead to large degrees of variability, an issue that is resolved by using tris(hydroxymethyl)aminomethane (Tris). Another source of variability is degradation of the protein of interest during the procedure. Enzymatic cell lysis with zymolyase can lead to extensive protein degradation, which is greatly reduced by mechanical lysis through bead beating. Degradation also occurs during sonication of chromatin, affecting large proteins in particular. An optimal mix of protease inhibitors and cross-linking with a higher percentage of formaldehyde reduces the extent of this degradation. Finally we also show that the immunoprecipitation step itself can be greatly improved with magnetic beads and optimized incubation/washing steps. The study results in a highly optimized protocol, which is shorter, easier to perform and has a stronger, more reproducible signal with less background. This protocol is presented in detail. In addition, the results highlight the greatest sources of variability in many other protocols, showing which steps are important to focus on for reproducible and quantitatively comparable ChIP experiments.

## Introduction

DNA is the carrier of genetic information and how it is decoded, replicated and packaged is largely determined by interactions between DNA and proteins. Studying protein-DNA interactions has therefore been a long-standing topic of interest in the field of molecular biology. There are several ways to assess which sites in the genome are bound by a specific protein *in vivo* (reviewed in Dey *et al*, 2012) and one of the oldest and most commonly used techniques is chromatin immunoprecipitation (ChIP). With this technique proteins and DNA are cross-linked and the protein of interest is separated from the chromatin extract using antibodies. The DNA that is bound to the protein of interest is subsequently identified and/or quantified (Figure 1).

**Figure 1.**
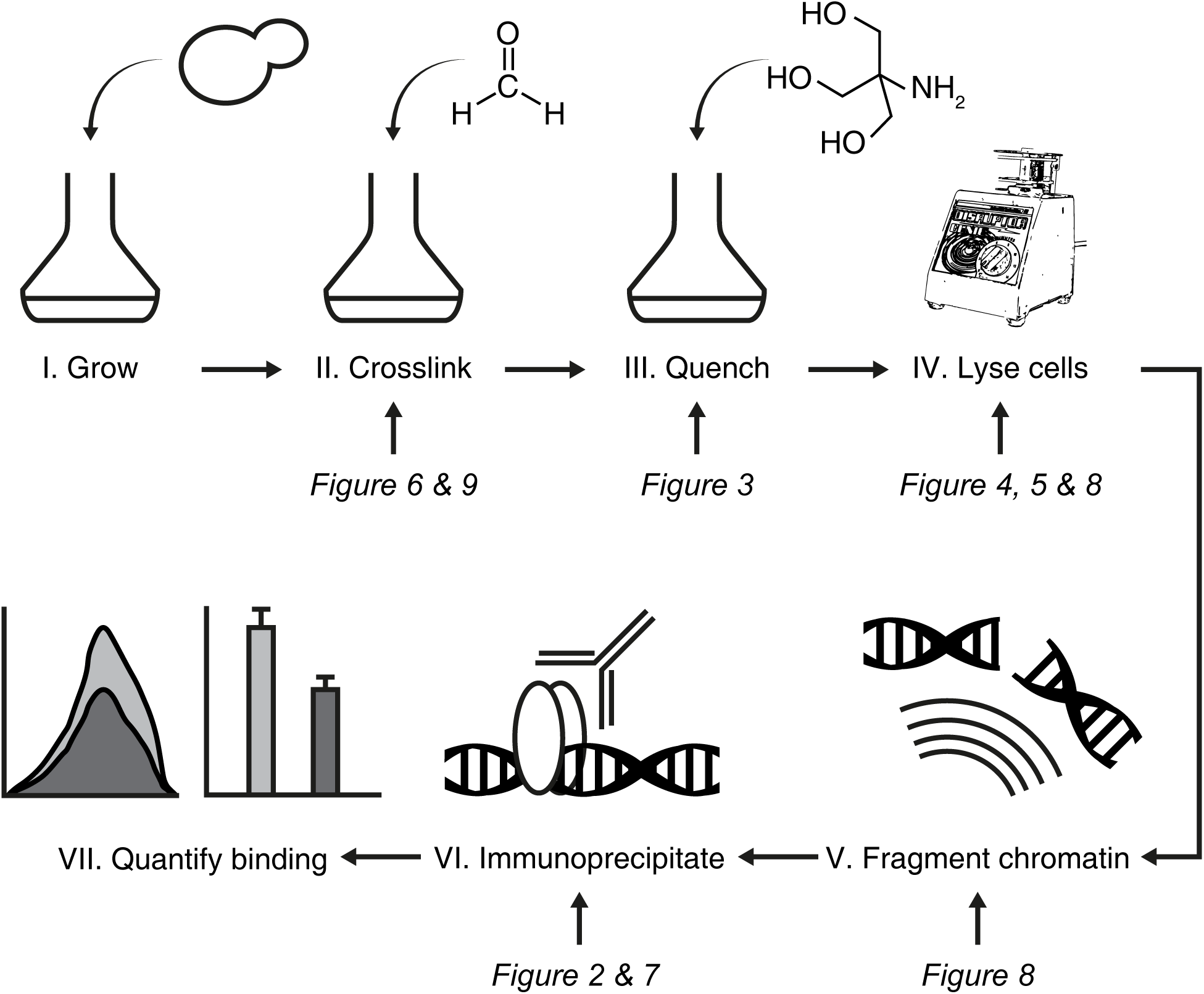
Overview of the ChIP protocol and steps optimized here. Schematic overview of the ChIP protocol. I. First, cells are grown and II. cross-linked using formaldehyde. III. Subsequently the formaldehyde is quenched with a quenching agent and IV. the cells are lysed either mechanically or enzymatically. V. After the cells are lysed, the chromatin is fragmented using sonication and VI. the protein of interest is immunoprecipitated. VII. The DNA that was bound by the protein of interest is isolated and quantified using for example quantitative PCR or sequencing. The arrows show which optimizations are described in which figure.

### The chromatin immunoprecipitation procedure

ChIP was developed in 1984 to investigate *in vivo* binding of RNA polymerase to two genes in bacteria (Gilmour & Lis, 1984). After its first application, the protocol was adapted for other species (reviewed Kuo & Allis, 1999) and modified extensively (O’Neill & Turner, 2003; Rhee & Pugh, 2011; Kasinathan *et al*, 2014; Skene & Henikoff, 2015; He *et al*, 2015; Gutin *et al*, 2018). Despite the many varieties of ChIP protocols, the key steps described in detail here, remain largely the same.

After obtaining the cells or tissue of interest (Figure 1, step I), the next step is often fixation of the protein-DNA interactions by cross-linking of the sample (Figure 1, step II), although in some protocols this step is omitted (Hebbes *et al*, 1988; O’Neill & Turner, 2003; Kasinathan *et al*, 2014). There are several ways to cross-link the samples, for example by irradiating with UV or by addition of cross-linking agents such as formaldehyde (Jackson, 1978; Gilmour & Lis, 1984; Solomon *et al*, 1988). Although in the first ChIP protocol UV cross-linking was used, the cross-linker of choice is usually formaldehyde. Formaldehyde is preferred due to ease of use and the fact that cross-links can easily be reversed by heating the sample. Additionally, formaldehyde cross-linking can be stopped by addition of a quenching agent (Figure 1, step III), although early adaptations of the protocol did not include a quenching step (Solomon *et al*, 1988; Dedon *et al*, 1991).

To gain access to the chromatin, the cross-linked cells have to be lysed (Figure 1, step IV). For most animal cells this is relatively straightforward, but it becomes challenging when working with organisms that have a cell wall such as bacteria, fungi or plants. The cell wall can be disrupted either by using enzymes such as lyticase or zymolyase, or mechanically by for example grinding the cells in liquid nitrogen or by vigorous shaking with glass beads.

With the cells lysed, the chromatin must be fragmented (Figure 1, step V). This step is needed to solubilize the chromatin and to make it accessible for the antibodies. Chromatin can be fragmented either mechanically or enzymatically. Mechanical fragmentation is often achieved using sonication. During sonication, high intensity sound waves exert a force that induces DNA breaks. Alternatively, enzymatic fragmentation with DNAses such as micrococcal nuclease (MNase) can be used to digest the DNA, by cleaving the DNA that is not protected by proteins.

Having access to the solubilized chromatin, the protein of interest can be immunoprecipitated (Figure 1, step VI). During the immunoprecipitation (IP) step the protein of interest is isolated from the rest of the chromatin using an antibody that specifically binds the protein of interest. The antibody can subsequently be conjugated to beads that can be harvested using either centrifugation or, in the case of magnetic beads, strong magnets. It is important that high-quality antibodies, specific to the protein of interest, are available. Otherwise, it is also possible to tag the protein of interest with a universal epitope-tag such as an HA-, FLAG- or V5-tag (Field *et al*, 1988; Hopp *et al*, 1988; Southern *et al*, 1991). High-quality antibodies are available for these tags, which makes the IP more efficient. Using the same tag for different proteins makes the IP more comparable between these proteins, because the same antibody can be used. However, this requires genetic modification of the cells before treatment to introduce the tag, which might not always be possible. Moreover, addition of the tag may alter the activity of the protein, potentially interfering with the result of ChIP.

The last step of any ChIP protocol is to quantify the binding of the protein of interest to DNA (Figure 1, step VII). This is performed by isolating the DNA that was bound to the protein of interest and quantifying this using (q)PCR or sequencing. When using qPCR to quantify the binding, the recovery of DNA bound by the protein of interest is often expressed as the percent of input. This is the signal of the immunoprecipitated sample divided by the signal found in the same sample that was not immunoprecipitated (input). The percentage of input recovered is determined by the efficiency of cross-linking, the efficiency of the IP step and the percentage of cells where the protein of interest was bound to the particular site. Theoretically, if a protein is bound to a certain locus in all cells and the efficiency of cross-linking and IP is 100%, the percent of input recovered is also 100%. High levels of recovery (e.g. 20-50%) are only achieved using highly abundant and very stably bound proteins such as histones. In general signals are typically much lower (e.g. 0.1-5%).

### Optimization of the chromatin immunoprecipitation procedure

Over the years many labs have optimized the ChIP protocol, leading to many versions of the method (O’Neill *et al*, 2006; Acevedo *et al*, 2007; Dahl & Collas, 2007; Goren *et al*, 2010; Adli *et al*, 2010; Goren *et al*, 2010; Brind’Amour *et al*, 2015). Most of these efforts focused on optimizing the protocol for a low number of mammalian cells using histone post-translational modifications or RNA polymerase II, which are easier to detect than transiently binding proteins. Often protocols allow the determination of where a protein of interest binds in the genome, but do not enable a quantitative comparison of binding to the same site in different samples. This limitation is not an issue if there is no interest in having accurate comparisons of binding levels, for example, when the main interest is to find where a specific protein is bound in the genome rather than at what level. On the other hand, if the aim is to determine how binding levels change under different conditions, accuracy of the measurements becomes crucial, as well as the ability to quantitatively compare between conditions. For these comparisons a quantitative and reproducible ChIP protocol is needed, but only a few efforts have focused on optimizing the protocol for this purpose.

Here, we describe how to make the ChIP protocol more quantitative and reproducible. This was achieved by optimizing the ChIP protocol using five different gene-specific transcription factors from the yeast *Saccharomyces cerevisiae*. Centromere binding factor 1 (Cbf1) is required for chromosome segregation (Cai & Davis, 1990) and can act both as a transcriptional activator and repressor (Kemmeren *et al*, 2014). ARS binding factor 1 (Abf1) is involved in transcription regulation (Buchman & Kornberg, 1990; Miyake *et al*, 2004), DNA repair (Reed *et al*, 1999) and replication (Rhode *et al*, 1992). RNA polymerase I enhancer binding protein (Reb1) is involved in transcription regulation of both RNA polymerase I and II transcripts (Chasman *et al*, 1990). Minichromosome maintenance 1 (Mcm1) is important for recombination and regulation of genes involved in arginine metabolism, cell cycle progression, cell wall maintenance and mating (Kuo & Grayhack, 1994; Messenguy & Dubois, 2003). Depending on the mating type Mcm1 can function either as an activator or as a repressor (Haber, 2012). Suppressor of mar1-1 (Sum1) is a repressor of middle-sporulation genes (Xie *et al*, 1999), but is also involved in activation of a subset of autonomous replicating sequences (ARS) (Irlbacher *et al*, 2005). Cbf1, Abf1, Reb1 and Mcm1 are known as general regulatory factors (GRFs). GRFs are abundant transcription factors (TF) (Ghaemmaghami *et al*, 2003) that can organize nucleosomes (Kent *et al*, 2004; Hartley & Madhani, 2009; Ganapathi *et al*, 2011) and all but Cbf1 are essential for viability (Passmore *et al*, 1988; Rhode *et al*, 1989; Cai & Davis, 1990; Ju *et al*, 1990). The yeast GRFs are therefore akin to chromatin pioneer TFs (Zaret & Mango, 2016).

All experiments described here were carried out using strains that harbor tagged TFs. The TFs were tagged with a V5-tag, which is part of a larger cassette containing GFP and an anchor away tag as well (Haruki et al, 2008). These V5 epitopes are recognized by the antibody used in all experiments described in this work.

Little effort has been done to make quantitative ChIP protocols. Therefore, here we optimized several key steps that are required to make the protocol more quantitative and reproducible. We highlight key points in the protocol that are a source of variation and point out artefacts that may arise during different steps of the procedure. Together, the findings presented here highlight common pitfalls during the ChIP procedure and result in a strongly revised protocol that is suitable for making quantitative and reproducible measurements of protein-DNA binding. This optimized protocol can be found in the supplemental materials.

## Results

### Use of magnetic beads improve ChIP enrichment

One of the key steps in any ChIP protocol is the IP step (Figure 1, step VI). During this step the protein of interest is bound by an antibody, which is then conjugated to beads prior to the precipitation step. The choice of antibody is crucial, as the specificity of the antibody determines the enrichment of genomic regions bound by the protein of interest over non-specific regions (background).

After the antibody binds to its epitope, the antibody is conjugated to beads. Typically these are either agarose or magnetic beads that can be separated using centrifugation or strong magnets, respectively. Agarose beads are often used, but magnetic beads offer the advantage of reduced washing steps and times. As a starting point for protocol optimization, we first tested whether there was a difference in ChIP signal (% of input) of a Cbf1 anchor away strain (Cbf1-aa) between these two types of beads. Figure 2A shows the difference in ChIP signal when using agarose or magnetic beads. Although the agarose beads have higher signal, the background signal is also increased, and therefore there is no enrichment of the Cbf1 targets (*YOS1* and *QCR10*) over the background (*ACT1* and *TUB1*, p value > 0.70). In contrast, when using magnetic beads there is a clear enrichment of *YOS1* over the background (p value = 0.0007), indicating that the ChIP enrichment can be improved by switching to magnetic beads.

**Figure 2.**
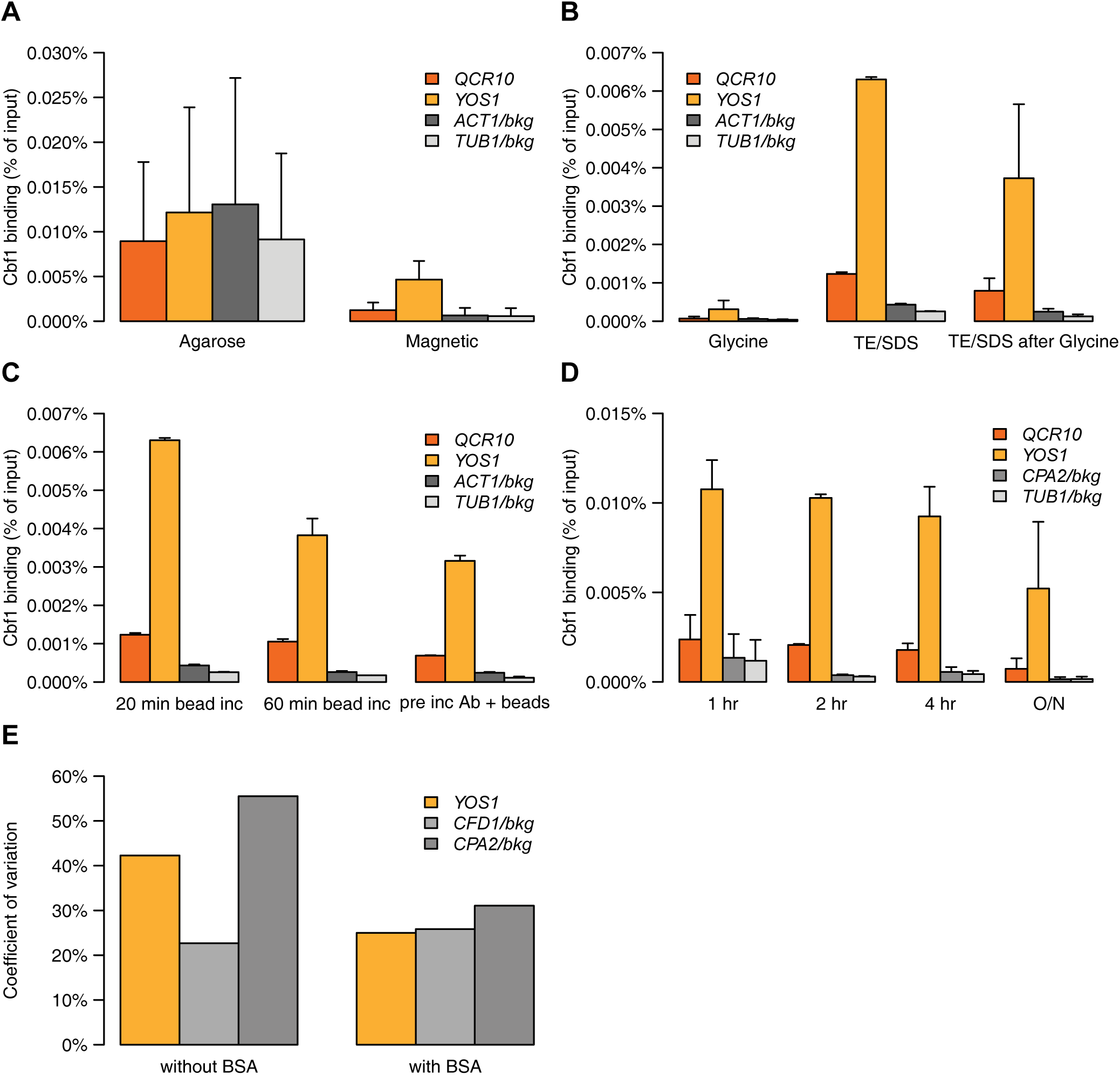
Improved speed and signal using magnetic beads. (A) Percent of input DNA recovered when using agarose versus magnetic beads during the IP. (B) Percent of input DNA recovered when eluting DNA from the beads using glycine, TE/SDS or first glycine and then TE/SDS. (C) Percent of input DNA recovered after incubating the chromatin and antibody with the magnetic beads for 20 minutes, 60 minutes or after first incubating the beads with the antibody and then incubating these with the chromatin. (D) Percent of input DNA recovered after incubating the chromatin with the antibody for 1 hour, 2 hours, 4 hours or overnight (O/N). (E) Variation in the percent of input as measured by the coefficient of variation after pre-incubating the beads without or with BSA. For all panels a Cbf1-aa strain was used. *ACT1, TUB1* and *CFD1* were used as background controls (gray) and *QCR10* and *YOS1* are targets of Cbf1 (McIsaac *et al*, 2012, orange). The number of replicates used varies per experiment. (A: Agarose and magnetic) have 4 and 8 replicates, respectively. (B and C) were performed using 2 replicates and (D and E) show the average of 3 replicates. Error bars represent the standard deviation for all experiments with 3 or more replicates (A, D and E) or the distance from the mean for experiments with two replicates (B and C). The same samples are shown in (B: TE/SDS) and (C: 20 min bead inc).

With the switch to magnetic beads, the best condition for eluting DNA from these beads was next investigated. The manufacturer of the magnetic beads suggests using a low pH glycine buffer to elute the protein of interest from the beads. Alternatively, a TE/SDS buffer at high temperature is often used to reverse the cross-links and elute the DNA from the beads. To determine which of the two is most efficient, these elution methods were compared. Eluting with glycine gives a much lower ChIP signal compared to eluting with a TE/SDS buffer (Figure 2B). This is highlighted by the observation that when glycine eluted beads are subjected to a second elution with TE/SDS, a large portion of the DNA can still be recovered. This shows that a low pH glycine buffer is not sufficient to elute the DNA from the beads and that it is best to use elution with a TE/SDS buffer.

When using magnetic beads, the recommended incubation time with the chromatin and antibody is 20 minutes. We hypothesized that longer incubation times might increase yield, but since the incubation is performed at room temperature this also increases the risk of proteolytic degradation of the sample. To determine whether a longer incubation time increases the ChIP signal, we tested incubation of the beads with the antibody and chromatin for 20 and 60 minutes. In addition, we also tested whether pre-incubating the antibody with the beads would increase the ChIP signal. The ChIP signal of YOS1 decreases by roughly half when the incubation time is increased from 20 to 60 minutes and pre-incubating the beads with the antibody before binding to the chromatin also decreases the ChIP signal (Figure 2C). This demonstrates that 20 minute incubation of the antibody bound to the chromatin with the beads yields the best signal.

One of the most time-consuming steps of the ChIP procedure is the incubation of antibody with chromatin. Often this step is carried out overnight, to ensure maximal binding of the protein of interest. If this step can be reduced to only a few hours, the protocol could be shortened by a day. For the anti-V5 antibody, reducing the incubation time increases the ChIP signal (Figure 2D), perhaps by reducing the amount of degradation that takes place during the incubation. When taking the signal and the variation into account, incubation of the chromatin with the V5 antibody for 2 hours is optimal, since this shows the strongest signal (% of input) with the lowest variation. This is an important improvement because compared to overnight incubation this not only increases the signal but also shortens the protocol by a day.

Besides improving the ChIP signal, another important goal is reducing the variation in both the specific and the background signal between replicate samples. The background signal can be caused by non-target proteins from the cells that stick to the beads aspecifically. This aspecific binding can be prevented by saturating the beads with proteins prior to antibody binding, which potentially reduces the variation in the background levels. Indeed, pre-incubating the beads with bovine serum albumin (BSA) shows a reduction in variation of the background (Figure 2E), indicating that addition of BSA can increase the reproducibility of the protocol by reducing the variation of the background signal.

Together these results show that the IP can be improved in several ways. The use of magnetic beads in combination with TE/SDS elution gives a better enrichment compared to agarose beads. The optimal incubation time of the anti-V5 antibody with chromatin is 2 hours and addition of BSA to the beads prior to antibody binding reduces variation. Besides the improved practicality and increased reproducibility, these optimizations also led to a greatly increased ChIP signal over background (compare figure 2A and 2D).

### Glycine is a poor quencher

An important first step of most ChIP procedures is cross-linking of proteins to DNA (Figure 1, step II). Cross-linking is stopped by quenching the cross-linking agent (Figure 1, step III). Proper quenching is an important step, especially in time course experiments where subsequent time points need to be quantitatively comparable. Most often the samples are cross-linked with formaldehyde and quenched by addition of glycine (Kuo & Allis, 1999; Acevedo *et al*, 2007; Rhee & Pugh, 2011; Poorey *et al*, 2013; Lara-Astiaso *et al*, 2014; He *et al*, 2015; Skene & Henikoff, 2015; Gutin *et al*, 2018). Glycine can quench formaldehyde because the amino group of glycine can react with formaldehyde, which prevents it from forming cross-links with other macromolecules. Remarkably, the concentration of glycine used is often sub-stoichiometric to formaldehyde (Kuo & Allis, 1999; Acevedo *et al*, 2007; Rhee & Pugh, 2011; Poorey *et al*, 2013; He *et al*, 2015; Skene & Henikoff, 2015; Gutin *et al*, 2018) and it has been shown that glycine is not necessarily an efficient quencher at these concentrations (Sutherland *et al*, 2008; Zaidi *et al*, 2017).

To investigate the effectiveness of quenching, a time course was performed. Yeast cultures were cross-linked with standard amounts of 1% formaldehyde (~333 mM) for 1 minute and subsequently quenched with 125 mM glycine for 1, 5 or 10 minutes. If glycine quenches formaldehyde efficiently there should be no difference in ChIP signal between the samples. However, there is a clear increase in ChIP signal with longer glycine incubation times (Figure 3A), which means that the cross-linking is not stopped by addition of glycine at a concentration of 125 mM. Potentially formaldehyde can be quenched more efficiently using a higher glycine concentration. When 250 mM glycine is added for 5 minutes the ChIP signal seems to decrease only slightly compared to quenching with 125 mM for 5 minutes (Figure 3B), and the ChIP signal still increases compared to 1 minute of quenching (compare Figures 3A and 3B). This confirms that formaldehyde can still form cross-links and that glycine, therefore, does not fully quench the formaldehyde when added at these levels.

**Figure 3.**
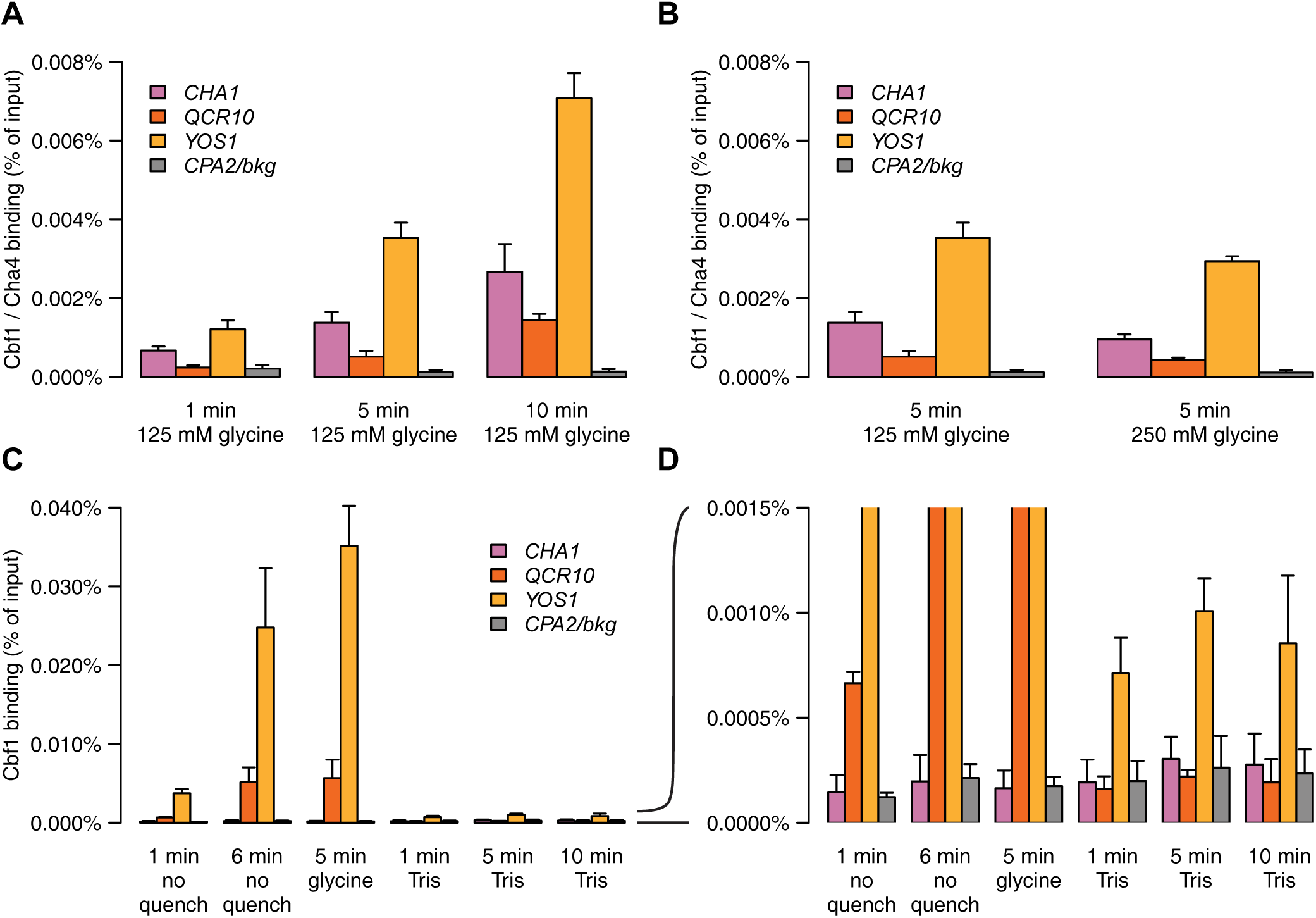
Tris is a superior quencher compared to glycine. (A) Percent of input DNA recovered after quenching for 1, 5 or 10 minutes with 125 mM glycine. (B) Percent of input DNA recovered when quenching with 125 mM or 250 mM glycine for 5 minutes. (C) Percent of input DNA recovered after cross-linking for 1 or 6 minutes without quenching, or 1 minute of cross-linking followed by quenching with glycine for 5 minutes or quenching with Tris for 1, 5 or 10 minutes. (D) Zoom in of (C) to 0.0015% recovery of input DNA. (A and B) were performed with a Cbf1-aa Cha4-V5 strain and (C and D) with a Cbf1-aa strain. *CPA2* was used as a background control (gray), *QCR10* and *YOS1* are Cbf1 targets (McIsaac *et al*, 2012, orange) and *CHA1* is a target of Cha4 (MacIsaac *et al*, 2006, pink). Error bars represent the standard deviation of 3 replicates. The same samples are shown for 5 min 125 mM in (A and B).

### Tris is a much more efficient quencher

Since glycine is not quenching properly when added at these levels, the quenching procedure has to be improved, since this will otherwise increase variation (see Discussion). Others have suggested two ways to quench more efficiently, either using very highly concentrated glycine solutions (Zaidi *et al*, 2017) or using excess Tris concentrations (Sutherland *et al*, 2008). Although highly concentrated glycine solutions (2.93M) do quench better than sub-stoichiometric glycine solutions (Zaidi *et al*, 2017), this requires concentrating the culture before cross-linking and addition of 440 ml glycine per 10 ml concentrated culture. This makes the procedure impractical and potentially stresses the cells during the steps needed to concentrate the cell culture. Quenching with Tris on the other hand, has been reported to be more efficient than with glycine (Sutherland *et al*, 2008) and thus can be achieved with lower concentrations. Tris can quench formaldehyde more effectively because a single Tris molecule is able to bind two formaldehyde molecules and it binds formaldehyde more stably than glycine (Hoffman *et al*, 2015). However, concerns have been raised that quenching with Tris will even de-cross-link because formaldehyde binding by Tris is so efficient (Hoffman *et al*, 2015; Zaidi *et al*, 2017). In contrast, others have demonstrated that prolonged incubation of cross-linked samples in Tris solution without heating does not de-cross-link (Sutherland *et al*, 2008; Kawashima *et al*, 2014).

To investigate if Tris quenches formaldehyde completely, and whether it de-cross-links the protein-DNA interactions, we performed the following experiment. Cbf1-aa cultures were cross-linked for 1 minute, without quenching, or with quenching either for 5 minutes using 125 mM glycine, or 1, 5 or 10 minutes using 750 mM Tris. Cultures that were cross-linked for 6 minutes without quenching were also taken along as a control for the samples that were quenched for 5 minutes. By comparing the samples that were cross-linked for 6 minutes without quenching with the samples that were quenched with glycine or Tris for 5 minutes, it is possible to determine to what extent glycine or Tris are able to quench formaldehyde. Surprisingly, there is no significant difference in the ChIP signal of *YOS1* (p value = 0.13) and *QCR10* (p value = 0.78) between samples that were cross-linked for 6 minutes and not quenched and samples that were cross-linked for 1 minute and quenched for 5 minutes with 125 mM glycine (Figure 3C). This indicates that glycine is not quenching at all at this concentration. In contrast, when using Tris the cross-linking is efficiently stopped, as is evident by the lower ChIP signal.

The ChIP signal of the Tris quenched samples are even lower than the samples that were cross-linked for 1 minute without quenching (Figure 3C). Although this could indicate that Tris de-cross-links the proteins from DNA, this is nevertheless unlikely. Firstly, for the non-quenched samples cross-linking is not stopped after 1 minute incubation with formaldehyde. After incubation, the samples are centrifuged for several minutes, and subsequently resuspended in Tris-buffered saline (TBS). The Tris present in this buffer will fully stop the cross-linking reaction. This means that the non-quenched samples were effectively cross-linked for several minutes. Cross-linking in the Tris quenched samples, on the other hand, was effectively stopped after 1 minute. The longer effective cross-linking time of the non-quenched sample likely leads to the observed higher ChIP signal. Secondly, prolonged incubation of the samples with Tris does not lower the ChIP signal (Figure 3D). If addition of Tris would de-cross-link proteins from DNA, the ChIP signal would have decreased with longer incubation times.

Together, these results show that glycine is a very poor quencher, that quenching with Tris is far more efficient and that short quenching times with Tris do not decross-link the samples. Thus, using Tris as a quenching agent improves the accuracy and reproducibility of the ChIP protocol, which is important for quantitative comparisons.

### Cbf1 is degraded during zymolyase treatment

Previous experiments focused on the immunoprecipitation and quenching steps. Another aspect that may be optimized is cell lysis (Figure 1, step IV). In contrast to mammalian cells, yeast cells have a strong cell wall that needs to be disrupted in order to gain access to the chromatin. In the experiments described so far, cells were lysed enzymatically by addition of zymolyase. Zymolyase digests the cell wall, creating spheroplasts that can subsequently be lysed by sonicating them in a lysis buffer. We reasoned that by extending the zymolyase treatment, a bigger proportion of cells would have their cell walls digested, potentially leading to a more efficient recovery of the protein of interest.

To test this, ChIP signals were compared for Cbf1-aa cells that were treated with zymolyase for either 10 or 25 minutes (Figure 4A). In contrast to our expectations, zymolyase treatment for 25 minutes decreased the ChIP signal compared to 10 minutes treatment. This suggests that there is a component present during zymolyase treatment that decreases the ChIP efficiency. One possibility is proteases, which are known to be present in zymolyase preparations. To test whether proteases were degrading the protein of interest during the zymolyase treatment, and thereby reducing the ChIP signal, we lysed cells using zymolyase with and without addition of protease inhibitors. Cells that were treated in the presence of protease inhibitors had a two-to threefold increase in ChIP signal (Figure 4B). This suggests that indeed Cbf1 is degraded by proteases during zymolyase treatment and that this decreases the ChIP signal.

**Figure 4.**
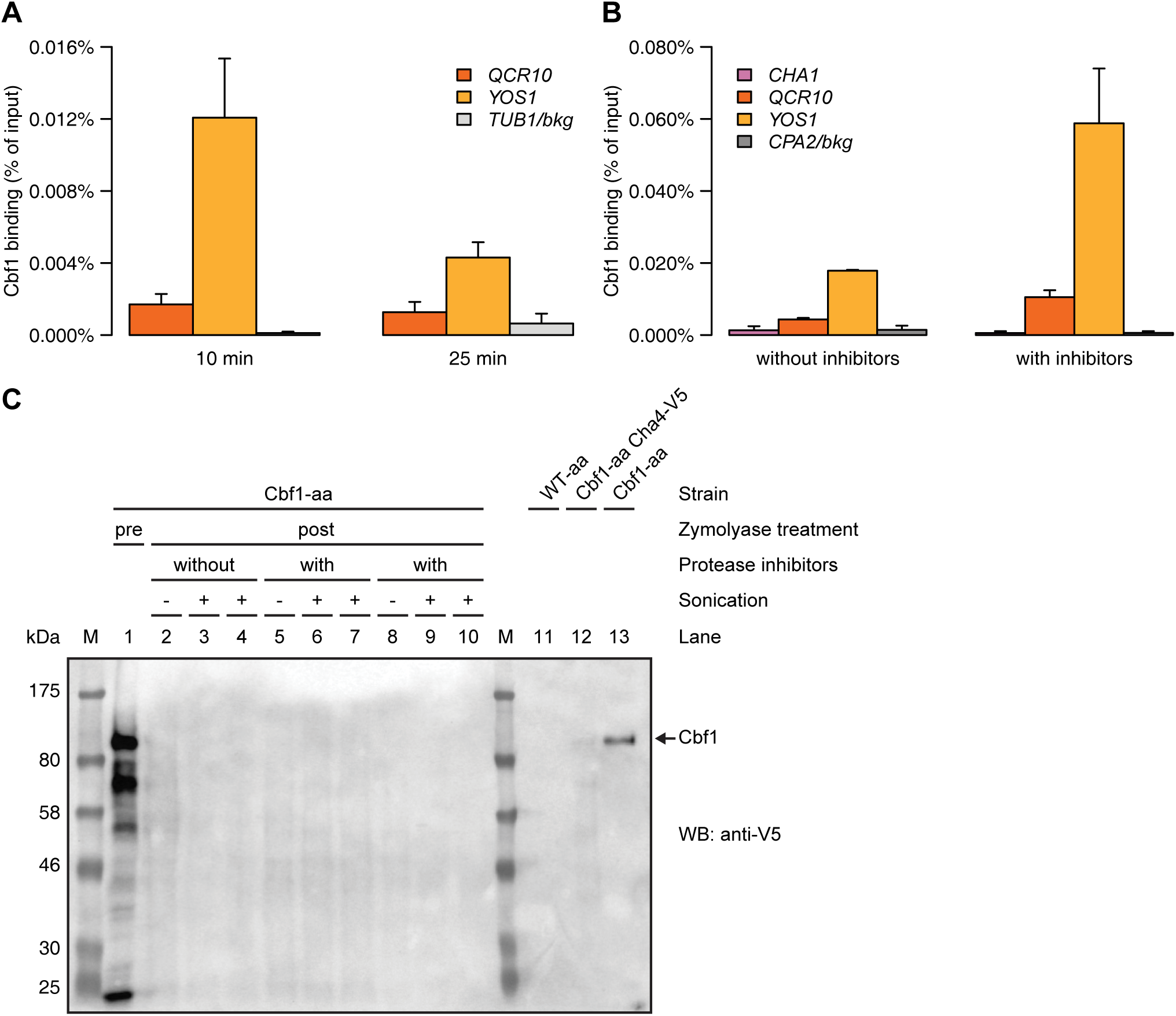
Zymolyase treatment causes degradation of proteins of interest. (A) Percent of input DNA recovered after treating the cells with zymolyase for 10 or 25 minutes. (B) Percent of input DNA recovered after treating the cells with zymolyase in the presence or absence of protease inhibitors. (C) Western blot showing the presence of Cbf1. Tagged Cbf1 has a mass of approximately 83 kDa. The location of tagged Cbf1 on the blot is indicated on the right. Different steps of the ChIP protocol were loaded on the gel: before lysis (pre-zymolyase treatment), after lysis (post-zymolyase, pre-sonication) and after the sonication (post-sonication). Cells were treated with zymolyase in the absence or presence of protease inhibitors. On the right side crude lysates were loaded as negative (WT-aa, the untagged parental strain) and positive (Cbf1-aa) controls. The blot was incubated with an antibody specific for V5. For (A and B) a Cbf1-aa strain was used without Cha4-V5 and for (C) both a Cbf1-aa and WT-aa, as indicated above the blot. In (A and B) *CPA2* and *TUB1* were used as background controls (gray), *QCR10* and *YOS1* are Cbf1 targets (McIsaac *et al*, 2012, orange) and *CHA1* is a target of Cha4 (MacIsaac *et al*, 2006, pink). The number of replicates used varies per experiment. (A) was performed with 3 replicates. (B: without inhibitors) had 2 replicates and (B: with inhibitors) shows the average of 4 replicates. Error bars represent the standard deviation for all experiments with 3 or more replicates (A and B: with inhibitors) or the distance from the mean for experiments with two replicates (B: without inhibitors).

To directly assess whether Cbf1 is degraded during zymolyase treatment, the amount of intact protein was measured at different steps during the protocol using Western blotting. The amount of intact Cbf1 was determined before zymolyase treatment, after zymolyase treatment with and without protease inhibitors as well as after sonication. Samples before and after sonication were taken along because the sonication process can potentially induce protein degradation (Pchelintsev *et al*, 2016). Before zymolyase treatment there is a strong Cbf1 band present (Figure 4C, lane 1), but, to our surprise, after zymolyase treatment there is no longer any detectable protein, even in the presence of protease inhibitors (Figure 4C, lanes 2-10). This indicates that most of the Cbf1 is degraded during zymolyase treatment, likely by the proteases present in the zymolyase preparation. Addition of protease inhibitors is likely not sufficient to prevent proteolytic degradation of the proteins of interest. This suggests that lysing the cells with a different mechanism that does not degrade the proteins being studied could strongly improve the ChIP signal.

### Mechanical cell disruption improves ChIP signal

An alternative mechanism to lyse cells is mechanical disruption. A common way to disrupt cells mechanically is by adding small glass beads to the cells in a tube containing lysis buffer and agitating the tubes at high speed, which is called bead beating. Since bead beating does not involve incubating the cells with proteases, we reasoned that this might improve the ChIP signal. Indeed, when comparing the zymolyase protocol with the beat beating protocol using a Cbf1-aa strain that also had a V5 tagged Cha4, there is a more than 10-fold increase in the ChIP signal of both TFs tested (Figure 5A). This shows that the ChIP signal can be improved by replacing zymolyase with mechanical disruption such as bead beating. However, the bead beat protocol is also more cumbersome and time-consuming.

**Figure 5.**
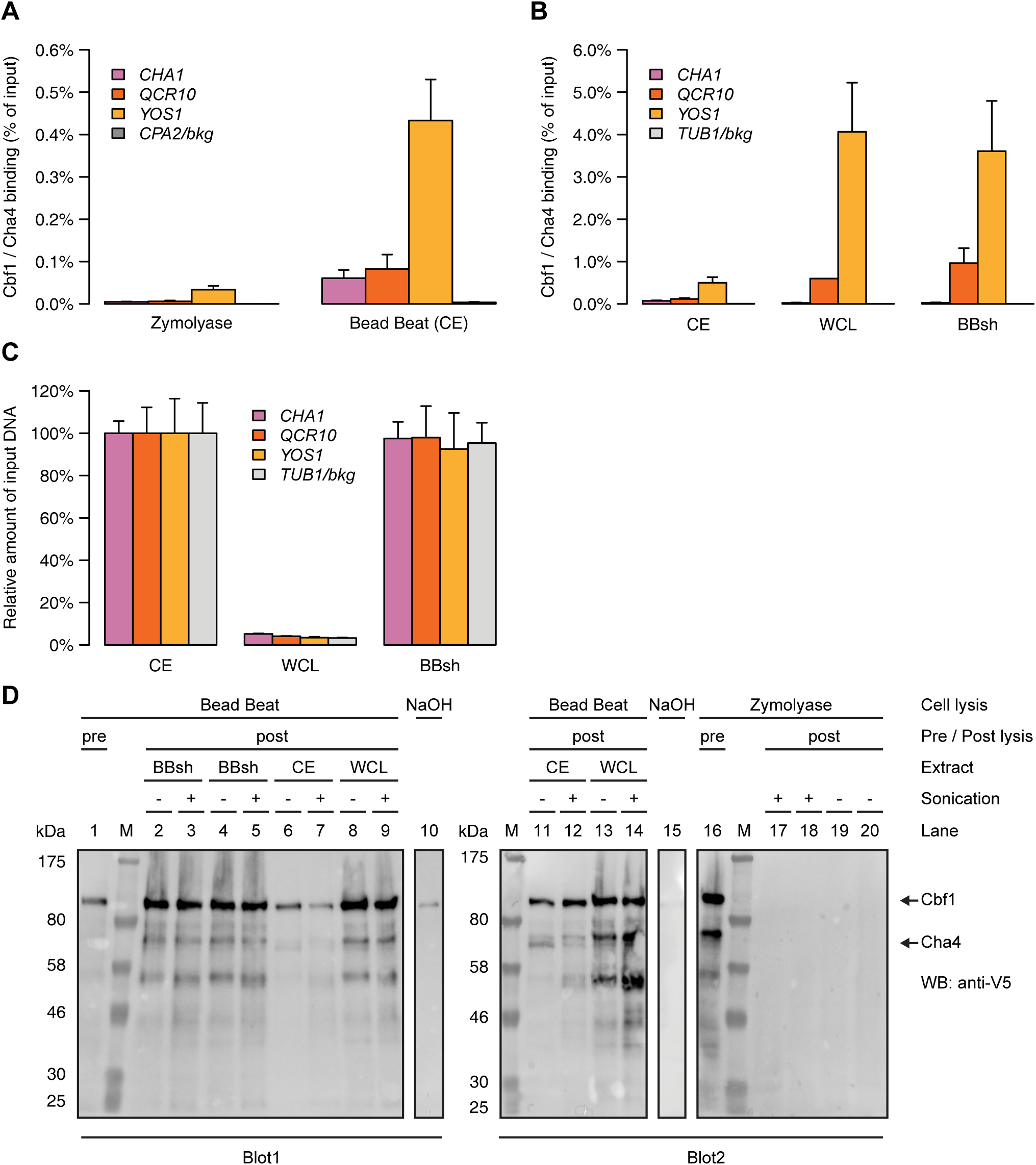
Cell lysis by bead beating greatly improves ChIP signal. (A) Percent of input DNA recovered after lysing the cells using either zymolyase or bead beating. (B) Percent of input DNA recovered after lysing the cells using the bead beat protocol with separation of chromatin extract (CE) and whole cell lysate (WCL), or without (BBsh). (C) Relative recovery of the input DNAs. The amount of input DNA recovered in the CE sample was set to 100% and the amount of input DNA recovered in the WCL and BBsh was scaled accordingly. (D) Western blot showing the presence of Cbf1 and Cha4. Tagged Cbf1 has a mass of approximately 83 kDa and Cha4-V5 has a mass of approximately 85 kDa. The location of these proteins on the blot is indicated on the right. Cha4 is less abundant than Cbf1, which makes it hard to detect on the blots. On both blots a crude lysate (NaOH) of a Cbf1-aa strain, without Cha4-V5 was used as a positive control. The different bead beat extracts and the zymolyase treated lysate before and after sonication are shown in duplicate. The blots were incubated with an antibody specific for V5. For (A, B and C) a Cbf1-aa Cha4-V5 strain was used, for (D) all but the NaOH samples were from a Cbf1-aa Cha4-V5 strain and for the NaOH samples a Cbf1-aa strain was used. In (A, B and C) *CPA2* and *TUB1* were used as background controls (gray), *QCR10* and *YOS1* are Cbf1 targets (McIsaac *et al*, 2012, orange) and *CHA1* is a target of Cha4 (MacIsaac *et al*, 2006, pink). The number of replicates used varies per experiment. (B and C: WCL) were performed using 2 replicates. (A) has 3 replicates and (B and C: CE and BBsh) show the average of 4 replicates. Error bars represent the standard deviation for all experiments with 3 or more replicates (A, B and C: CE and BBsh) or the distance from the mean for experiments with two replicates (B and C: WCL).

Therefore, our next aim was to determine if the number of steps in the protocol could be reduced, speeding up the procedure and possibly also increasing the reproducibility. A step that potentially can be omitted during cell lysis is the last step, when chromatin is separated from the rest of the cell lysate. During this step the lysate is centrifuged at high speed for 20 minutes, which concentrates the cross-linked chromatin. The supernatant, called the whole cell lysate (WCL), is discarded while the pellet, called the chromatin extract (CE), is resuspended and used for the IP. However, resuspending this pellet is often quite difficult, which could be a potential source of variability. If the IP is similarly efficient without separating the CE from the rest of the WCL, this could allow for a faster and potentially more robust protocol. To assess if the protocol is similarly efficient without the last centrifugation step, the ChIP signal of the CE and the WCL from the full bead beat protocol was compared to the signal obtained with a shorter protocol without the last centrifugation step (BBsh). This comparison was made using a Cbf1-aa strain that also had a V5 tagged Cha4. Interestingly, the WCL and BBsh samples showed an even stronger ChIP enrichment of the Cbf1 targets *YOS1* and *QCR10* compared to the CE (Figure 5B). The Cha4 target *CHA1*, on the other hand, does not show increased signal using the shorter protocol or the WCL. This indicates that, at least for Cbf1, a higher ChIP signal can be obtained by using the short protocol or the WCL. It is important to note, however, that while the WCL has a high percent of input recovered, it has much lower absolute levels of DNA (Figure 5C), because most of the DNA is contained in the CE. Since more than 95% of the DNA is lost, the WCL is not suited for doing IPs.

The higher ChIP signal obtained with the bead beat protocols compared to the zymolyase protocol could be explained by an increased stability of the proteins. To verify if this was the case, the protein stability was compared between the different bead beat protocols and versus the zymolyase protocol. There is an improvement in protein stability with the bead beat protocols compared to the zymolyase protocol, since the protein levels are now readily detectable by Western blot (Figure 5D). Nevertheless, there is still some degradation taking place during the bead beat protocols (Figure 5D, lanes 2-9 and 11-14), but only to a limited extent compared to the zymolyase treatment (Figure 5D, lanes 17-20). This increase in stability explains the higher ChIP signals observed with the bead beat protocols and shows that the protocol can be improved by lysing the cells mechanically. Using a shorter version of the protocol allows for a faster protocol that has even stronger ChIP signal for Cbf1, but not for Cha4.

### Cbf1 may re-bind DNA during the IP step

The strong increase in Cbf1 ChIP signal using the short bead beat protocol suggests that this protocol is the best for Cbf1 ChIP. However, in contrast to Cbf1, the ChIP signal of Cha4 does not increase with the short protocol (Figure 5B). This suggests that perhaps the increase in Cbf1 ChIP signal is artefactual, and may not reflect actual *in vivo* binding. To test this, Cbf1-aa cells were grown with and without cross-linking, and subjected to the short bead beat protocol. Indeed, even without cross-linking Cbf1 binding can easily be detected (Figure 6A). In contrast, two other TFs, Abf1 (Figure 6C) and Reb1 (Figure 6D), do not show any binding without cross-linking. The binding of Cbf1 without cross-linking could be explained either by strong DNA binding, which does not need cross-linking to be retained, or by binding of unbound Cbf1 to these sites during the IP procedure. To test the latter, Cbf1 was depleted from the nucleus for 60 minutes using the anchor away technique (Haruki *et al*, 2008). The depleted cells were cross-linked for 5 minutes and the ChIP signal was compared to Cbf1 binding without depletion. Cbf1 is depleted to background levels from the nucleus after approximately 15 minutes (Figure 6B). Thus, depletion of Cbf1 from the nucleus for 60 minutes should abolish Cbf1 binding to DNA. Therefore, if there is still ChIP signal detected after nuclear depletion, this would indicate that Cbf1 rebinds DNA during the IP step. Indeed, with Cbf1 depleted from the nucleus, the binding levels are only reduced by one third compared to no depletion (Figure 6A), while Abf1 (Figure 6E) and Reb1 (Figure 6F) show at least a fourfold reduction in binding after nuclear depletion.

**Figure 6.**
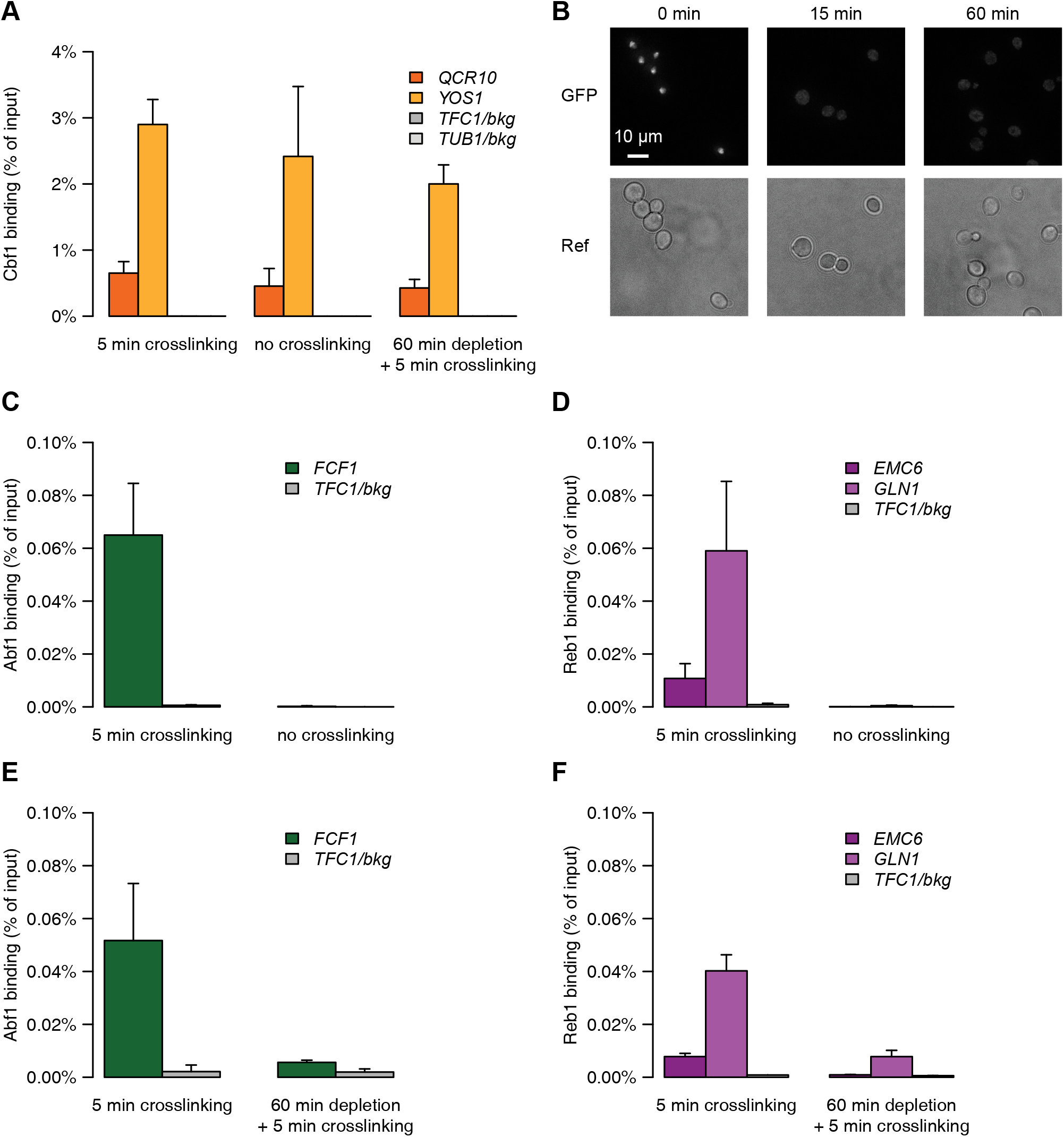
Cbf1 can re-bind DNA during immunoprecipitation. Percent of input DNA recovered after cross-linking the cells for 5 minutes, in the absence of cross-linking or when Cbf1 is depleted from the nucleus for 60 minutes prior to cross-linking for 5 minutes. (B) Fluorescence microscopy images of Cbf1-FRB-GFP-3V5 before, 15 minutes and 60 minutes after induction of nuclear depletion. Scale bar: 10 µm. (C and D) Percent of input DNA recovered with 5 minutes or without cross-linking for (C) Abf1 and (D) Reb1. (E and F) Percent of input DNA recovered before or after 60 minutes of depletion of (E) Abf1 and (F) Reb1 from the nucleus prior to cross-linking for 5 minutes. For (A and B) a Cbf1-aa strain, for (C and E) an Abf1-aa strain and for (D and F) a Reb1-aa strain was used. In all experiments the cells were lysed using the short bead beat protocol. *TFC1* and *TUB1* were used as background controls (gray), *QCR10* and *YOS1* are Cbf1 targets (McIsaac *et al*, 2012, orange), *FCF1* is an Abf1 target (Kasinathan *et al*, 2014, green) and *EMC6* and *GLN1* are Reb1 targets (Kasinathan *et al*, 2014, purple). (A: 5 min cross-linking and 60 min depletion, C, D, E and F) were performed using 3 replicates and (A: no cross-linking) had 6 replicates. Error bars represent the standard deviation.

These observations suggest that free and depleted, cytosolic Cbf1 can re-bind DNA during the IP step because in the short bead beat protocol the cytosolic and nuclear fraction are mixed during the IP step. Unfortunately, this makes the short bead beat protocol in this form unsuitable for determining *in vivo* Cbf1 binding levels, since binding may arise *in vitro* during the IP procedure. However, it can still be used for other proteins, such as Abf1, Cha4 and Reb1, which do not bind DNA during the IP step on the promoters tested.

### Re-optimization of the immunoprecipitation step

So far, several steps of the ChIP protocol have been optimized and changed (Figures 1-6). Because many such changes were made after the initial optimizations of the immunoprecipitation step, we re-evaluated the way the immunoprecipitation is performed (Figure 1, step VI). Since the short bead beat protocol is not well suited for determining binding levels of Cbf1, the optimizations were carried out using a Sum1-aa strain. Sum1 has a lower abundance than Cbf1 (Ghaemmaghami *et al*, 2003), and may therefore have a lower ChIP signal. Thus, using Sum1 would allow for optimization of the IP procedure also for less abundant proteins.

As a starting point, the elution was re-investigated. As shown in Figure 2B, elution of the immunoprecipitated DNA with TE/SDS is more efficient than eluting with low pH glycine. Still, this does not necessarily show that all DNA is efficiently eluted. To confirm that all DNA is eluted, an IP was performed with overnight elution using TE/SDS and after the first elution a second elution was performed on the same beads, again with TE/SDS. If overnight elution is not sufficient to elute all DNA, a significant amount of DNA should be recovered in the secondary elution. Figure 7A shows that when a second elution is performed, only about 2% of the initial elution is recovered. This demonstrates that overnight elution is sufficient, as the vast majority of DNA is eluted in this step.

**Figure 7.**
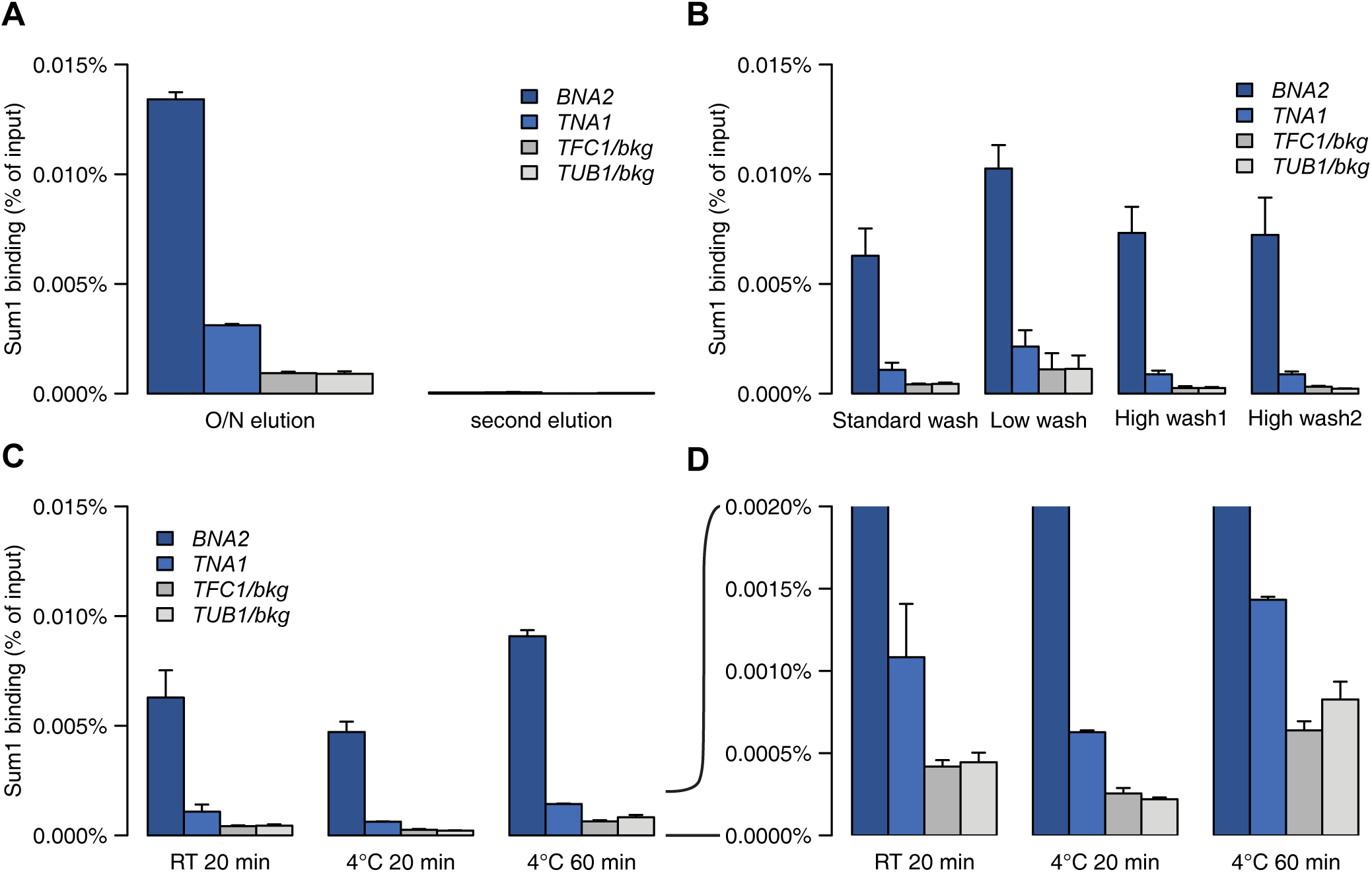
Re-optimization of the immunoprecipitation for the bead beat protocol. (A) Percent of input DNA recovered after eluting the DNA from the beads overnight or after a second elution was performed on the same beads. (B) Percent of input DNA recovered after using different washes during the IP, see Materials and Methods for details. (C) Percent of input DNA recovered after incubating the chromatin and the antibody with the beads at room temperature (RT) for 20 minutes or at 4°C for 20 or 60 minutes. (D) Zoom in of (C) to 0.002% recovery of input DNA. For all panels a Sum1-aa strain was used. *TFC1* and *TUB1* were used as background controls (gray) and *BNA2* and *TNA1* are Sum1 targets (MacIsaac *et al*, 2006, blue). The number of replicates used varies per experiment. (A, B: high wash1 and C and D: 4°C 20 min and 4°C 60 min) were performed using 2 replicates. (B: standard wash and C and D: RT 20 min) both have 3 replicates and (B: low wash and high wash1) show the average of 4 replicates. Error bars represent the standard deviation for all experiments with 3 or more replicates (B: standard wash, low wash and high wash2 and C and D: RT 20 min) or the distance from the mean for experiments with two replicates (A, B: high wash1 and C and D: 4°C 20 min and 4°C 60 min).

After confirming the efficiency of the elution, we checked whether washing of the beads could be further optimized. The manufacturer of the magnetic beads recommends two washes with PBS followed by two washes with PBS-Tween, which has been used as the standard wash in the experiments described so far. To explore which washing conditions are best for the IP of Sum1, this standard wash was compared to a less stringent wash and two kinds of more stringent washes: high wash1 and high wash2 (see Materials and Methods). Doing a less stringent wash leads to a higher ChIP signal, but also to an increase in background signal (Figure 7B). On the other hand, both stringent washes had similar Sum1 ChIP signals compared to the standard wash, but a reduction in the background signal of about 40%. Therefore, at least for Sum1, washing more stringently can improve the signal to noise ratio.

Lastly, incubation of the chromatin and antibody with the beads was optimized. This incubation is normally carried out at room temperature (RT) for 20 minutes. Proteases are more active at RT, thus reducing the temperature during this step could lead to less degradation and higher ChIP signal. On the other hand, lowering the temperature could also reduce the efficiency of antibody binding to the beads. To test the effect of temperature on this step, the ChIP signal was compared between samples that were incubated at RT for 20 minutes and at 4°C for 20 or 60 minutes. When the chromatin and antibody are incubated with the beads at 4°C for 20 minutes there is a small decrease in ChIP signal compared to 20 minute incubation at RT (Figure 7C). Incubating at 4°C for 60 minutes increases the ChIP signal, but also increases the background (Figure 7D). This indicates that there is no added benefit of doing the incubation at 4°C.

In summary, no changes were made to the short bead beat protocol, because overnight elution with TE/ SDS and incubation of chromatin and antibody with the beads for 20 min at RT are sufficient. The signal to noise ratio of the IP could be further improved by washing more stringently.

### Protein degradation during sonication

We next examined how well the short bead beat protocol was performing using other TFs. When the stability of Abf1 was analyzed using the short bead beat protocol, a substantial amount of degradation was observed during cell lysis (Figure 8A, lanes 1-6). This degradation was even more pronounced after the sonication step (Figure 8A, lanes 2 and 5), with only 1% of the signal in the lane coming from intact Abf1. This means that the vast majority of Abf1 that was present during the IP was degraded, which can have detrimental effects on the ChIP efficiency. During the bead beat protocol, proteases are inhibited by addition of a tablet containing a cocktail of protease inhibitors. The substantial amount of Abf1 degradation can potentially be explained by insufficient inactivation of proteases by these protease inhibitors. To assess if proteases could be inhibited more efficiently, addition of four separate protease inhibitors during multiple steps in the protocol was tested. These protease inhibitors were aprotinin, pepstatin, leupeptin and PMSF (collectively abbreviated as APLP), which together inhibit a broad spectrum of proteases. With the addition of APLP, there is almost no additional degradation during cell lysis, and the degradation during sonication is also reduced (Figure 8A, lanes 7-12). As expected, besides reduced degradation of Abf1, there was also an increase in ChIP signal of Abf1 targets *FCF1* and *NHX1* when APLP is added (Figure 8B). This indicates that addition of APLP at multiple steps in the protocol rather than a protease inhibitor tablet at the beginning, offers better protection against proteases, and thus a stronger ChIP signal. Therefore, a revised version of the short bead beat protocol (version 2) is performed with the addition of APLP at multiple steps rather than a protease inhibitor tablet at the start.

**Figure 8.**
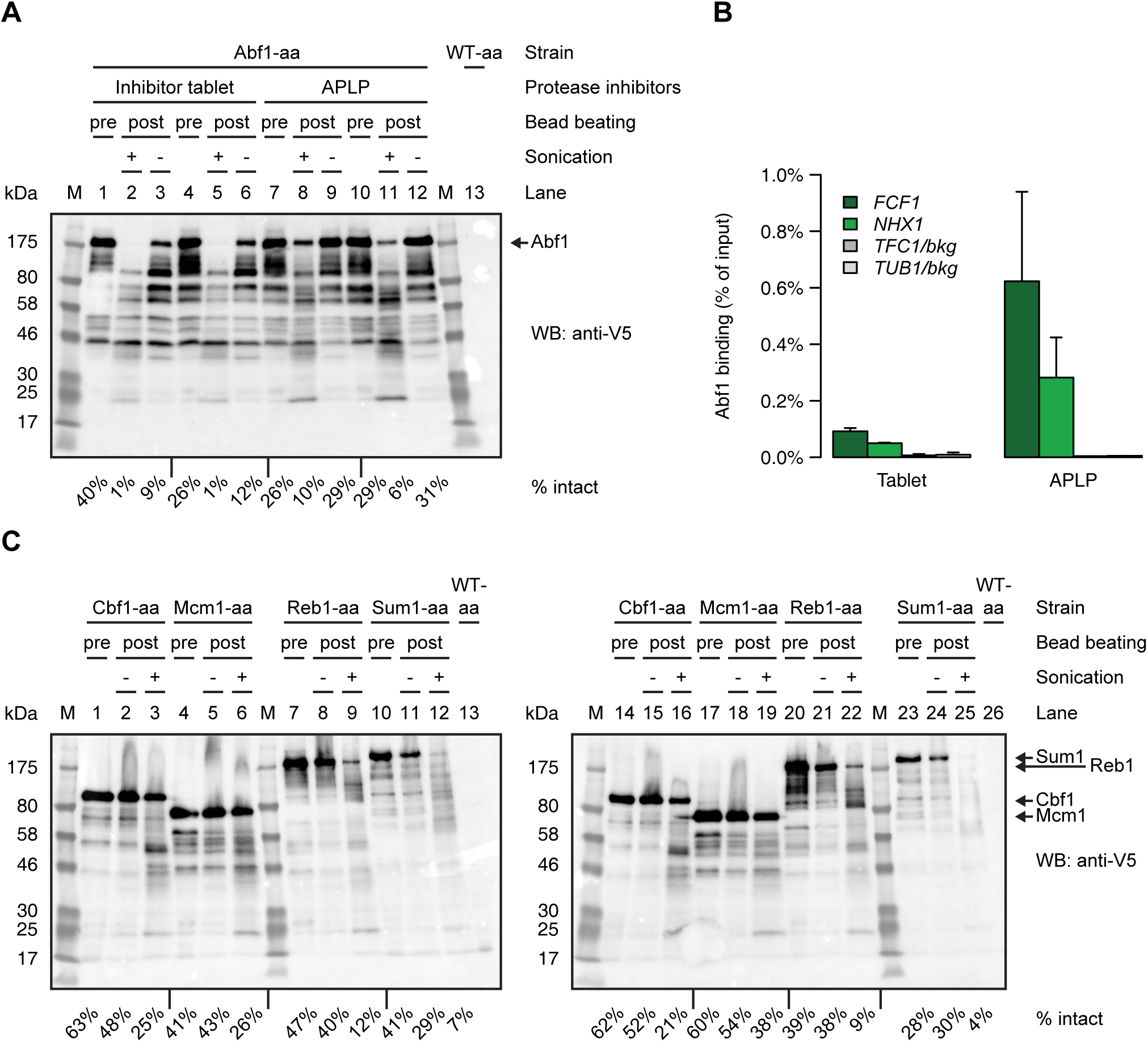
Addition of separate protease inhibitors partly prevents degradation during sonication. (A) Western blot showing the presence of Abf1, which was obtained with either addition of a protease inhibitor tablet or addition of separate protease inhibitors (aprotinin, pepstatin, leupeptin and PMSF). Samples before bead beating, after sonication and before sonication are shown. (B) Percent of input DNA recovered of the samples shown in (A). Two replicates were used and the error bars show the distance from the mean. *TFC1* and *TUB1* were used as background controls (gray), *FCF1* and *NHX1* are Abf1 targets (Kasinathan *et al*, 2014, green) (C) Western blots showing presence of Cbf1, Mcm1, Reb1 and Sum1. Samples before bead beating, before sonication and after sonication are shown. All the blots shown were incubated with an antibody specific for V5. Tagged Abf1, Cbf1, Mcm1, Reb1 and Sum1 have a mass of approximately 125, kDa, 83 kDa, 76 kDa, 135 kDa and 162 kDa respectively. The locations of these proteins on the blot are indicated on the right side of the corresponding blots. For all blots a crude lysate from a WT-aa strain was used as a negative control. Below all blots is indicated what percentage of the protein was still intact. This was calculated by dividing the signal from the intact protein (upper band) by the total signal in the lane (upper band + all lower bands).

After establishing the superior protection of Abf1 from proteases by APLP, we asked whether the stability of other proteins was also affected during the ChIP protocol. Therefore, the amount of degradation was determined for other TFs as well (Cbf1, Mcm1, Reb1 and Sum1), using the short bead beat (version 2) protocol. All four TFs were degraded during sonication to some extent, but especially the larger proteins, Reb1 (135 kDa) and Sum1 (162 kDa) were more severely affected compared to the smaller proteins Cbf1 (83 kDa) and Mcm1 (76 kDa, Figure 8C, compare lanes 7-12 and 20-25 with lanes 1-6 and 14-19). This indicates that large proteins are more sensitive to degradation during sonication, as expected and shown previously (Pchelintsev *et al*, 2016). The degradation could be caused by residual protease activity, the mechanical force during the sonication or a combination thereof.

### Increased formaldehyde concentration increases ChIP signal

The last part of the protocol that was re-optimized was the cross-linking step (Figure 1, step II). It has been shown that for some TFs cross-linking efficiency can be improved by using a higher concentration of formaldehyde (Zaidi *et al*, 2017). An improved cross-linking efficiency means that with a similar time of cross-linking, more cross-links are formed between the protein of interest and DNA, which should result in an increased ChIP signal. To test this, Abf1-aa and Reb1-aa strains were cross-linked with 1%, 2% or 3% formaldehyde and processed with the short bead beat (version 2) protocol. Comparing the integrity of Abf1 treated with increasing amounts of formaldehyde indicates that a higher concentration of formaldehyde leads to less degradation during sonication (Figure 9A, compare lanes 1-3 with lanes 4-12). Whereas for Abf1 a similar increase in stability is seen between 2% and 3% formaldehyde, for Reb1 this increase in stability was only observed with 2% formaldehyde (Figure 9B, compare lanes 17-19 with lanes 14-16 and 20-25). As expected, with an increased formaldehyde concentration, the ChIP signal of Abf1 also increased (Figure 9C). The ChIP signal of Reb1 increased with 2% and 3% formaldehyde compared to the 1% formaldehyde ChIP signal, but was highest with 2% formaldehyde (Figure 9D). This suggests that for both proteins the ChIP signal can be improved by using higher concentrations of formaldehyde, although it has to be pointed out that these experiments were carried out using only single replicates. Note that due to a handling error, a similar amount of IP material was lost for the 1% and 2% Abf1 samples, which means that these signals would otherwise have been higher. In conclusion, a higher percentage of formaldehyde can increase the ChIP signal for both Abf1 and Reb1, not only by increasing cross-linking efficiency but likely also by preventing a part of the proteolytic degradation during sonication.

**Figure 9.**
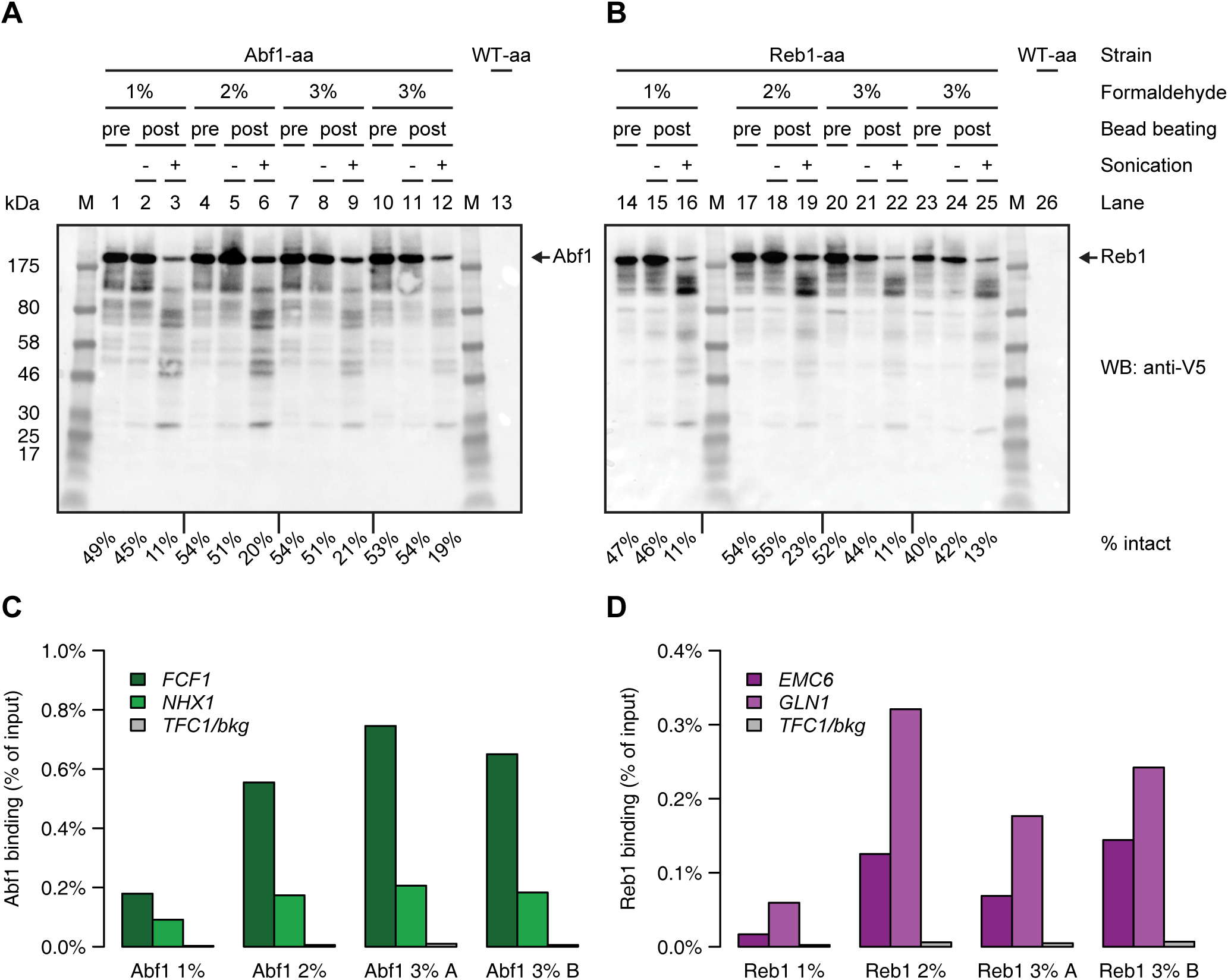
A higher percentage of formaldehyde leads to less degradation and higher ChIP signal. (A and B) Western blots showing the presence of (A) Abf1 and (B) Reb1 with increasing concentrations of formaldehyde. Samples before bead beating, before sonication and after sonication are shown. For both blots a crude lysate from a WT-aa strain was used as a negative control. Tagged Abf1 and Reb1 have a mass of approximately 125 kDa and 135 kDa, respectively. The location of these proteins on the blot is indicated on the right side of the corresponding blots. Both blots were incubated with an antibody specific for V5. Below the blots is indicated what percentage of the protein was still intact. This was calculated by dividing the signal from the intact protein (upper band) by the total signal in the lane (upper band + all lower bands). (C and D) Percent of input DNA recovered for Abf1 (C) and Reb1 (D), the samples are the same as shown in (A and B). Single replicates were used. *TFC1* was used as a background control (gray), *FCF1* and *NHX1* are Abf1 targets (Kasinathan *et al*, 2014, green) and *EMC6* and *GLN1* are Reb1 targets (Kasinathan *et al*, 2014, purple). The 3% formaldehyde samples differed only in the final concentration of Tris used for quenching. The 3% A samples (lanes 7-9 and 20-22) were quenched with a final concentration 2.1M Tris and the 3% B samples (lanes 10-12 and 23-25) were quenched with a final concentration of 1.5M Tris.

### An optimized ChIP protocol

This study results in several improvements to a standard ChIP protocol (see detailed protocol in the supplemental materials). The use of magnetic beads rather than agarose beads improves the speed and practicality of the IP. Using magnetic beads also leads to reduced background signal and a higher ChIP enrichment. Quenching with glycine is very inefficient at levels commonly used in ChIP protocols. Concentrated Tris is a much more efficient quencher. In addition, the method for cell lysis was optimized, since zymolyase treatment can degrade the protein of interest, likely by the presence of proteases. This step can be improved by mechanically disrupting the cells using bead beating, which causes less degradation and a strongly increased ChIP signal compared to zymolyase treated samples. The bead beating protocol can be further improved by removing the last centrifugation step, leading to an easier and quicker protocol. However, care has to be taken that the protein of interest does not re-bind DNA during the subsequent incubation steps. There is a substantial amount of degradation taking place during cell lysis and sonication as well. Addition of a more optimal mix of protease inhibitors at several steps during the protocol, rather than a tablet containing a mix of inhibitors added only once, can alleviate part of this degradation. Nonetheless, even with these inhibitors there is still a substantial amount of degradation observed during sonication, especially for very large proteins. Cross-linking with a higher concentration of formaldehyde may reduce the extent of this degradation for some proteins.

Based on the results shown here we recommend the following protocol: 1) cross-linking the cells with 2% formaldehyde and 2) quenching them with an appropriate concentration of Tris (1.5M). 3) Not lysing the cells using zymolyase but lysing them mechanically, using for example a bead beater. 4) Using a shorter version of the bead beat protocol, but taking care that the protein of interest does not bind DNA *in vitro* during the protocol. 5) Adding the protease inhibitors aprotinin, pepstatin, leupeptin and PMSF during cell lysis and subsequent steps. 6) Using magnetic beads for the IP steps, as this allows for a faster and more reproducible IP, with stronger ChIP signal and less background. The complete optimized protocol can be found in the supplemental materials.

## Discussion

### Extent of cross-linking affects downstream steps

We optimized multiple steps of the protocol, and one of the steps that varies strongly between ChIP protocols is the time of cross-linking. In general, longer cross-linking times lead to stronger ChIP signals. Increased crosslinking can result in stronger signal in two ways. Firstly, this increase in signal can be explained by proteins that bind DNA during cross-linking (Poorey *et al*, 2013). These additional protein-DNA interactions get fixed by the formaldehyde, which will be picked up as increased signal. Alternatively, cross-linking of the protein of interest to DNA may not be very efficient and with longer incubation times there is an increased chance of the protein being cross-linked (Zaidi *et al*, 2017). Either one or both may be happening, depending on how efficiently the protein studied can be cross-linked to DNA.

Considering that longer cross-linking times lead to stronger ChIP signals, it might be tempting to increase the cross-linking time as long as possible. However, in addition to crosslinking protein-DNA interactions, formaldehyde can also crosslink protein-protein interactions (Sutherland *et al*, 2008). Prolonged crosslinking can cause extensive crosslinking of proteins to other proteins, and large protein aggregates could form in the cells. Therefore, the extent of cross-linking can have a significant effect on downstream processing steps. For example, during cell lysis there is a centrifugation step that clears cell debris from the chromatin. With increased cross-linking times a large part of the chromatin may co-precipitate with the cell debris, depending on the centrifugation speed. This could lead to reduced yields if the majority of the chromatin is subsequently discarded with the cell debris. The efficiency of sonication also changes with cross-linking time. Increased cross-linking leads to stiffer and more rigid DNA, which may be more resistant to shearing by sonication. More sonication cycles may be needed to sufficiently fragment the DNA and this could have detrimental effects on the stability of the protein of interest.

Apart from technical issues, long cross-linking times also increase well-known ChIP artefacts. Highly expressed genes tend to show as peaks in ChIP data (Park *et al*, 2013; Teytelman *et al*, 2013) and this is more prevalent with increased cross-linking times (Baranello *et al*, 2015). This artefactual signal might stem from the open chromatin that is present at these loci, or from crosslinking of the protein of interest to other DNA-bound factors. Additionally, with longer cross-linking times we also noticed an increase in background signal and of the non-tagged controls. Thus, longer cross-linking times may lead to increased signal, but not necessarily to an increased enrichment over background or biologically meaningful signal.

An alternative to longer cross-linking times is increasing the concentration of formaldehyde. A higher concentration of formaldehyde will increase the efficiency of cross-linking, which could increase the ChIP signal if this efficiency is rate-limiting. However, it may also increase ChIP artefacts, depending on their cause. In our optimized protocol we use 5 minutes cross-linking with 2% formaldehyde as a good balance between a relatively short cross-linking time, but still long enough to get clear signal. Note, however, that optimal cross-linking conditions may differ depending on the proteins studied.

### Proper quenching is crucial for quantitative protocols

The results presented here show that glycine is an inefficient quencher (Figure 3) (Sutherland *et al*, 2008; Zaidi *et al*, 2017). Interestingly, most published ChIP protocols that cross-link with formaldehyde, quench with glycine at sub-stoichiometric concentrations (Kuo & Allis, 1999; Acevedo *et al*, 2007; Rhee & Pugh, 2011; Poorey *et al*, 2013; He *et al*, 2015; Skene & Henikoff, 2015; Gutin *et al*, 2018). How is it possible that such an inefficient quenching agent became the quencher of choice for virtually all ChIP protocols? A likely explanation lies in its inefficiency as a quencher, which is linked to the stronger ChIP signals that can be observed with longer cross-linking times. Inefficient quenching effectively means that the cross-linking time is increased and thus that there is increased ChIP signal. Considering this, it is easy to imagine that glycine was preferred as a quencher, since it would result in a higher signal. An alternative explanation could be that during the development of the early ChIP protocols inactivation of formaldehyde was only desirable to discard the waste in a safe way. If the only concern is to inactivate formaldehyde for waste routing purposes, it is apparently less critical to efficiently and rapidly quench formaldehyde, and thus to efficiently stop the cross-linking reaction.

Considering that glycine does not quench and that the first ChIP protocols did not include a quenching step (Solomon *et al*, 1988; Dedon *et al*, 1991), the question arises as to whether it is really necessary to quench at all. In fact it may be better not to quench and to be aware that the cross-linking reaction is still proceeding, rather than quenching inefficiently and assuming that the reaction has stopped. If it is assumed that the reaction is efficiently stopped, quenching with glycine gives a false sense of security and this can have detrimental effects on the reproducibility. For example, this can happen when some samples are left on ice after quenching while others are being processed, because it is assumed that the cross-linking has stopped. In fact, cross-linking is still proceeding, thus the effective cross-linking time will be longer than intended. Even worse, the extent of cross-linking in this case will vary between samples or replicates. This can be a major source of variation and make the protocol irreproducible.

The experiments here show that in contrast to glycine, Tris is an efficient quencher (Figure 3), as has indeed been proposed previously (Sutherland *et al*, 2008). Some have raised concerns that quenching with Tris may revert the protein-DNA cross-links (Hoffman *et al*, 2015; Zaidi *et al*, 2017). The results here show no evidence of de-cross-linking when using short Tris incubation times, as has been shown by others for long incubation times without heating (Sutherland *et al*, 2008; Kawashima *et al*, 2014). An alternative to quenching with Tris is using highly concentrated (2.93 M) glycine solutions (Zaidi *et al*, 2017). While it is a more efficient way to quench compared to sub stoichiometric glycine solutions, it also involves concentrating the samples using centrifugation. This concentrating of the samples potentially stresses the cells. This, in combination with the impracticality of quenching with 2.93M glycine, which involves addition of 440 ml 3M glycine to 10 ml of cells, makes quenching with Tris preferable.

In conclusion, quenching with sub stoichiometric levels of glycine is very inefficient and a source of variation in ChIP protocols. Quenching with Tris is much more efficient, quicker, does not require an extra centrifugation step and does not de-cross-link the sample. Together this makes Tris a better quencher than glycine. The quenching step that we now incorporated in the optimized protocol facilitates higher reproducibility of the procedure.

### Strategies that reduce protein degradation increase ChIP signal

Many steps of the ChIP protocol can have an effect on the stability of the protein studied, including the extent of cross-linking, cell lysis, shearing of the DNA and the IP step. The steps that have the biggest influence on the stability of the proteins studied are cell lysis and sonication. In particular, cell lysis with zymolyase caused a significant decrease in detectable levels of the protein of interest in the samples (Figure 4). This is likely caused by proteases that are present in zymolyase preparations. A significant part of this degradation can be prevented by lysing cells mechanically using bead beating with appropriate protease inhibitors (Figure 5). Although it was not tested here, using additional protease inhibitors and more extensive washes may also prevent part of the degradation caused by zymolyase treatment.

Even when cells are used that do not have a cell wall, cell lysis is an important step. Differences in lysis efficiency between different mammalian cell types are a major source of variation in ChIP protocols (Arrigoni *et al*, 2016). A lysis strategy that is efficient across different cell types can greatly reduce variation between protocols (Arrigoni *et al*, 2016), which highlights the importance of the reproducibility of this step.

The second step that causes significant degradation is sonication. Using more efficient protease inhibitors reduces the amount of degradation during this step, but cannot prevent all degradation (Figure 8). This indicates that the mechanical force of the sonication also degrades the protein of interest. Large proteins are more sensitive to this degradation compared to smaller proteins (Pchelintsev *et al*, 2016). Thus, sonication may be further optimized to reduce the amount of degradation, but care has to be taken that the fragmentation of the DNA is still suited for downstream applications in terms of reproducibility and fragment size.

Even after the optimizations described here, there is still a substantial amount of degradation of most proteins that were studied. Future efforts to further reduce the extent of this degradation likely will increase the ChIP signal even more. For example, as an alternative to sonication, the DNA can also be fragmented enzymatically using DNases such as MNase (Skene & Henikoff, 2015; Arrigoni *et al*, 2018; Gutin *et al*, 2018). This approach could prevent the degradation that was observed during sonication and may subsequently increase the ChIP signal. Especially for large proteins this approach may be beneficial.

### Limitations and outlook

Although the optimizations carried out here greatly increase the specific ChIP signal and reduce variation, there are still some aspects that have to be taken into account before extrapolation to other experiments. This protocol was optimized using yeast. Most likely the optimizations are applicable for ChIP protocols using other organisms as well, but this has not been tested. In addition, all optimizations were performed using V5-tagged strains and an antibody that specifically recognizes this tag. Some conditions that were found to be optimal here might be specific for this antibody and should be verified when performing ChIP using other antibodies.

The optimizations described here were not all carried out with identical numbers of replicates and with the same proteins. In addition, even though this protocol has been extensively optimized, there is still some variation between replicates. This is clearly evident in Figures 3 and 7, where there is a difference in ChIP signal between ChIPs performed using the same conditions on different days. This highlights the need for the additional optimization steps carried out afterwards. Still, the protocol may yet be further improved. For example, optimizations that additionally reduce the number of steps or pool the samples early in the protocol could be beneficial to further reduce the variation between replicates. An example of such a strategy is barcoding of the DNA while it is bound to the beads (Lara-Astiaso *et al*, 2014; Schmidl *et al*, 2015; Wallerman *et al*, 2015; Gutin *et al*, 2018) or fragmentation and barcoding of DNA while still in the nucleus (Arrigoni *et al*, 2018). Alternatively, many orthogonal method exist that quantify protein-DNA binding, which may be used if no clear ChIP signal can be obtained (Galas & Schmitz, 1978; Garner & Revzin, 1981; Siebenlist & Gilbert, 1980; Siebenlist *et al*, 1980; Smith, 1985; Nick & Gilbert, 1985; Ellington & Szostak, 1990; Tuerk & Gold, 1990; Wang & Reed, 1993; Yin *et al*, 1995; van Steensel & Henikoff, 2000; Bulyk *et al*, 2001; Schmid *et al*, 2004; Cremazy *et al*, 2005; Lushnikov *et al*, 2006; Maerkl & Quake, 2007; Zentner *et al*, 2015; Skene & Henikoff, 2017). Using the short bead beat protocol, one protein (Cbf1) was able to re-bind DNA during the IP step of the protocol (Figure 6). Depending on the affinity of the protein for DNA, more stringent washes may remove such binding.

Although many labs have undertaken efforts to optimize the ChIP protocol, it remains a challenging technique. Many different protocols exist and most will not be ideal for the majority of proteins or conditions studied. Since many proteins of interest will have a different set of optimal ChIP parameters, ideally every ChIP protocol should be optimized for each protein separately. Nevertheless, this work highlights the steps that are crucial for accurate ChIP quantifications. The final protocol, described in full detail in the materials and methods section, is superior to previous standard protocols and will form an excellent starting point for many studies employing ChIP.

## Materials and Methods

Here, the general methods used are described. In the supplemental methods the differences from the general methods are highlighted for each figure separately as well as 3 supplemental tables that describe the strains, primers and reagents that were used in this study. The full, optimized ChIP protocol, including additional comments and notes, is also included in the supplemental materials.

### Strains

All the *Saccharomyces cerevisiae* strains that were used are anchor away strains (Haruki *et al*, 2008) that were re-made in the BY4742 background, as is described in (de Jonge *et al*, 2017). The anchor away tag consists of the FRB domain from the mammalian target of rapamycin (mTOR), a yeast enhanced green fluorescent protein (yEGFP) and three times a V5 tag (3V5). In addition to the Cbf1-aa strain, an additional strain was made where Cha4 was tagged with a 3V5 tag as an internal control. The Cbf1-aa strain with a 3V5 tagged Cha4 was used in Figures 3A, 3B and 5. The genotypes of all strains are shown in Table 1.

### Growth conditions

The growth conditions were similar for all the experiments performed. Strains were streaked directly from −80°C stocks on appropriate selective plates (YPD + Nourseothricin for the WT-aa strain and YPD + Hygromycin + Nourseothricin for anchor away strains) and grown for 3 days at 30°C. Liquid pre-cultures were inoculated in the morning, diluted at the end of the day and grown overnight at 30°C with shaking (230 rpm) in synthetic complete (SC) medium: 2 g/l dropout mix complete and 6.71 g/l yeast nitrogen base without amino acids, carbohydrate & w/AS (YNB) from US Biologicals (Swampscott, USA) with 2% D-glucose. The next morning, cultures were diluted in 100 ml to an OD of 0.2 (WPA Biowave CO8000 Cell Density Meter) and grown at 30°C, 230 rpm for 2 doublings until an OD of 0.8 was reached, which corresponds to about 2*10^7^ cells per ml.

### Cross-linking and quenching

When the cultures reached an OD of 0.8, formaldehyde was added to a final concentration of 1%, by adding 2.7 or 2.8 ml of 37% formaldehyde (Sigma-Aldrich #252549) to a 100 ml culture (Figure 2E, 3, 6, 8 and 9). When 250 ml cultures were used (Figure 2A-D, 4, 5 and 7), 6.8 ml 37% formaldehyde was added. A final concentration of 1% was used for all experiments except those shown in Figure 9, where some cultures were cross-linked with a final concentration of 2% (5.7 ml of 37% formaldehyde) or 3% (8.8 ml of 37% formaldehyde). To accurately control the time of cross-linking, the addition of formaldehyde and quenching agent was performed outside the shaker, on a heated stir plate (IKA C-MAG HS 7) set to 30°C and ~250 rpm, with a magnetic stir bar. An exception was made for the 250 ml cultures that were grown as a big batch. For these cultures, the formaldehyde and quencher were added and mixed in the incubator (Figure 2A-D, 4, 5 and 7).

Quenching was performed using either glycine or Tris (tris(hydroxymethyl)aminomethane). Glycine was added to a final concentration of 125 mM by adding 5.1 ml of 2.5M glycine to a 100 ml culture, except when otherwise indicated (Figure 3B, 250 mM: addition of 10.3 ml 2.5M glycine). When 250 ml cultures were used, 12.8 ml of 2.5M glycine was added. Glycine quenching was used for all experiments shown in Figure 2, 3 and 4. When the samples were quenched using Tris, this was done with a final concentration of 500 mM (Figure 5 and 7), 750 mM (Figure 3C and 3D, 6, 8 and 9: 1% samples), 1.5M (Figure 9: 2% and 3% B samples) or 2.0M (Figure 9, 3% A samples) Tris pH 8.0. Cells that were grown in 250 ml cultures were quenched using 500 mM Tris by adding 32.1 ml 4.5M Tris pH 8.0 (Figure 5 and 7) or using 750 mM Tris by adding 51.4 ml 4.5M Tris pH 8.0 (Figure 8A and 8B). When 100 ml cultures were quenched with 750 mM Tris, 20.6 ml 4.5M Tris pH 8.0 was added (Figure 6, 8C and 9). The cultures that were quenched with 1.5M Tris after cross-linking with 2% or 3% formaldehyde were quenched by adding 52.9 ml or 54.4 ml of 4.5M Tris pH 8.0, respectively (Figure 9). For the 3% formaldehyde cross-linked samples that were quenched with 2.0M Tris, 87.1 ml 4.5M Tris pH 8.0 was added (Figure 9). Quenching with glycine was done for 5 minutes and with Tris for 1 minute, unless otherwise stated (Figure 3).

### Harvesting cross-linked cells

100 ml cultures were harvested after quenching by splitting the culture over either 2 (glycine quenched), 3 (750 mM and 1.5M Tris quenched) or 4 (2.0M Tris quenched) 50 ml tubes. The cells were pelleted by centrifugation at 3220g (4000 rpm) in an Eppendorf 5810 R centrifuge at 4°C for 3 minutes. The supernatant was discarded, the pellet from the first tube was resuspended in 10 ml TBS (150 mM NaCl, 10 mM Tris pH 7.5) and subsequently combined with the other pellets that came from the same culture. The cells were pelleted again at 3220g for 3 minutes at 4°C, resuspended in 1 ml MQ and transferred to a 2 ml safe-lock tube. The cells were centrifuged at 3381g (6000 rpm) in an Eppendorf 5424 centrifuge at room temperature (RT) for 20 seconds. The supernatant was discarded and the samples were subsequently snap frozen and stored at −80°C.

250 ml cultures were harvested slightly differently. After quenching, the cells were centrifuged in 500 ml centrifuge bottles at 2831g (4000 rpm) in a Beckman Coulter Avanti J-E 369003 centrifuge for 3 minutes at 4°C. The supernatant was discarded and the pellets were pooled per 2 pellets by resuspending the first pellet in 40 ml ice-cold TBS (150 mM NaCl, 10 mM Tris pH 7.5), combining this with another pellet and transferring them to a 50 ml tube. By combining 2 pellets, the tube contains the equivalent of a 500 ml culture. The cells were next pelleted by centrifugation at 3220g (4000 rpm) in an Eppendorf 5810 R centrifuge at 4°C for 3 minutes. The supernatant was discarded, the cells were resuspended in 4.5 ml ice-cold MQ and divided in 1 ml aliquots over 5x 2 ml safe-lock tubes. After centrifuging at 3381g in an Eppendorf 5424 centrifuge for 20 seconds at RT, the supernatant was discarded and the cells snap frozen in liquid nitrogen. Each pellet now contains the equivalent of a 100 ml culture.

### Nuclear depletion

Nuclear depletion was achieved by using the anchor away system (Haruki *et al*, 2008). The nuclear depletion was induced by addition of 2 mM DMSO dissolved rapamycin to the cultures to a final concentration of 7.5 µM. The rapamycin was added in such a way that the samples were ready for harvesting at OD = 0.8. For 60 and 15 minutes depletion of Cbf1 (Figure 6A and 6B), 367.4 µl of 2 mM rapamycin was added to a 100 ml culture at OD = 0.52 and 0.72 respectively. In addition, 367.4 µl of DMSO was added to the non-depleted control at OD = 0.52. For nuclear depletion of Abf1 and Reb1 (Figure 6E and 6F) 758 µl and 750 µl 2 mM rapamycin was added to 215 ml and 200 ml cultures at OD = 0.56, respectively. For both Abf1 and Reb1 the same amount of DMSO (758 µl and 750 µl respectively) was added to the non-depleted controls at OD = 0.56.

### Fluorescence microscopy

Fluorescence microscopy was used to assess the degree of Cbf1 depletion from the nucleus. Cbf1, like all other TFs used in this study, was tagged with yEGFP, which allows monitoring of nuclear localization. To assess the degree of nuclear depletion of Cbf1-aa, cells were grown and Cbf1 was depleted from the nucleus as detailed in the previous sections. After depletion, the cells were fixed using methanol as described previously (Haruki *et al*, 2008). At OD = 0.8, after either 60 minutes of DMSO, or 60 minutes or 15 minutes of rapamycin treatment, 1 ml aliquots of the cultures were transferred to 1.5 ml Eppendorf tubes. The cells were collected by centrifugation at 3381g (6000 rpm) in an Eppendorf 5424 centrifuge for 1 minute at RT. The supernatant was discarded and the cells were fixed in 1 ml −20°C 100% methanol for 6 minutes. The cells were again pelleted at 3381g, and rehydrated in PBS containing 0.2% Tween-20 for 5 minutes. After centrifuging the cells at 3381g, the cells were resuspended in 100 µl PBS. 1.5 µl was used for imaging, by combining this with 1.5 µl of 1% agarose on a microscopy slide (50 × 24 mm) and quickly covering the cells with a coverslip (22 × 22 mm).

The cells were imaged using a DeltaVision Elite high resolution microscope, with an Olympus 100X/1.40 oil objective and a CoolSNAP HQ2-ICX285 camera operating at −25°C. The resolution was set to 512×512 pixels and binning to 2×2. To visualize GFP, EX and EM filters were set to FITC, the ND filter was set to 100% and an exposure time of 0.1 seconds was used. For each image, a z-stack of 11 images was taken with optical sectioning set to 1 µm and the images were deconvolved using the Resolve3D softWoRx software. A projection was made from the deconvolved images using the maximum intensity with the softWorx software. These projections are shown in Figure 6B. All GFP images shown here were set to have the same brightness and contrast levels.

Along with each z-stack of GFP images, a single reference image was taken. This was achieved by setting the EX filter to POL, the EM filter to BLANK, the ND filter to 100% and by using an exposure time of 0.01 seconds. The reference images were processed as follows. Two blank images were taken on the same microscope using the same settings without any cells in the view. The average signal of these two blank images was calculated and from this blank image a value of 100 was subtracted using the subtract function in the ImageJ software (Schneider *et al*, 2012). Subsequently, this reduced blank image was subtracted from each reference image using the image calculator function in ImageJ. The brightness and contrast values were then set to the same levels for all reference images.

### Chromatin isolation using zymolyase

Chromatin was isolated using zymolyase for the experiments shown in Figure 2, 3, 4, 5A and 5C. To isolate the chromatin, each cell pellet, which is equivalent to a 100 ml culture of cells OD = 0.8, was resuspended in 1 ml buffer Z (1M sorbitol, 50 mM Tris pH 7.5) to change buffer. The cells were pelleted by centrifugation at 15871g (13000 rpm) for 30 seconds in an Eppendorf 5424 centrifuge at RT. The cells were then resuspended in 1 ml of buffer Z containing 10 mM β-mercaptoethanol and 10 mg/ml zymolyase (zymolyase 20T MP biomedical #08320921) and incubated in a rotating wheel for 10 minutes (unless stated otherwise) to create spheroplasts. For some samples shown in 4B and 4C protease inhibitors were added at this step (one tablet of Roche EDTA-free cOmplete protease inhibitor cocktail (#11873580001) per 10 ml buffer Z containing β-mercaptoethanol and zymolyase). The cells were pelleted by centrifugation at 15871g for 10 seconds at room temperature and each pellet was washed with ice-cold buffer Z by inverting the tubes. The centrifugation and wash were repeated once. For some samples shown in 4B and 4C, protease inhibitors were added at this wash step as well (one tablet Roche cOmplete protease inhibitor cocktail (#11836145001) per 10 ml buffer Z). The spheroplasts were carefully resuspended in 550 µl FA lysis buffer (50 mM HEPES-KOH pH 7.5, 150 mM NaCl, 1 mM EDTA pH 8.0, 1% Triton X-100, 0.1% Na-deoxycholate, 0.1% SDS) with protease inhibitors (1 tablet Roche cOmplete protease inhibitor cocktail per 25 ml of FA lysis buffer) and split over 2x 1.5 ml bioruptor pico microtubes (Diagenode). The cells were lysed and the chromatin was sheared in a bioruptor pico sonicator (Diagenode) by sonicating the samples for 3 or 4 cycles, 30 seconds on / 30 seconds off. The cell debris and unfragmented chromatin was subsequently pelleted by centrifugation at 21130g (15000 rpm) in an Eppendorf 5424 R centrifuge for 20 minutes at 4°C. The 2 supernatants, containing the fragmented chromatin, of the samples that were split before sonication, were then combined again in a 1.5 ml Eppendorf tube. The chromatin was snap frozen in liquid nitrogen and stored at −80°C, except for a 5 µl aliquot that was taken to assess the extent of fragmentation of the DNA.

### Chromatin isolation using the full bead beating protocol

Chromatin was isolated using the full bead beating protocol for a few samples shown in Figure 5. To isolate the chromatin, each cell pellet, which is equivalent to a 100 ml culture of cells OD = 0.8, was resuspended in 700 µl of FA lysis buffer (50 mM HEPES-KOH pH 7.5, 150 mM NaCl, 1 mM EDTA pH 8.0, 1% Triton X-100, 0.1% Na-deoxycholate, 0.1% SDS containing protease inhibitors (1 tablet Roche cOmplete protease inhibitor cocktail (#11836145001) per 25 ml of FA lysis buffer). The resuspended cells were subsequently added to a 2-ml screw-cap tube containing 500 µl of zirconium/silica beads 0.5 mm (BioSpec Products, #11079105z). If needed, the tubes were closed and inverted once to remove air trapped in the beads. The tube was then filled completely with FA lysis buffer, to keep as little air as possible. The tubes were then bead beated 7 times for 3 minutes in a genie disruptor. The samples were put one ice for 1 minute in between each run. During the bead beating, 15 ml tubes were prepared containing 1 ml pipette tips with the end cut off. After bead beating, the lysate was recovered by burning a hole in the top and bottom of the screw-cap tubes using a hot 23G needle and placing them on top of the pipette tip in the 15 ml tube. This combination was then centrifuged at 201g (1000 rpm in an Eppendorf 5810 R) at 4°C for 1 minute and the lysate was transferred to a 2-ml screw-cap tube. To remove the majority of unbroken cells and cell debris, the lysate was centrifuged at 1503g (4000 rpm) for 2 minutes in an Eppendorf 5424 R centrifuge at 4°C. The supernatant was transferred to a new 2-ml Eppendorf tube and centrifuged at 18407g (14000 rpm) for 15 minutes in an Eppendorf 5424 R centrifuge at 4°C. The pellet (chromatin extract, CE) and the supernatant (whole cell lysate, WCL) were then treated differently.

### Chromatin extract (CE)

The CE was washed using 1.5 ml FA lysis buffer by resuspending the pellet using a 23G needle and incubating in a rotating wheel at 4°C for 30 minutes. Chromatin was pelleted again by centrifugation at 18407g (14000 rpm) for 15 minutes in an Eppendorf 5424 R centrifuge at 4°C. For Figure 5A, this pellet was washed again once with 700 µl FA lysis buffer without resuspending the pellet, but this step was skipped for the other experiment (Figure 5B-5D). The pellet was then resuspended in 600 µl FA lysis buffer using a syringe and a 23G needle and split over 2x 1.5 ml bioruptor pico microtubes (Diagenode). The chromatin was sheared in a bioruptor pico sonicator (Diagenode) by sonicating the samples for 3 cycles, 30 seconds on / 30 seconds off. The chromatin was snap frozen in liquid nitrogen and stored at −80°C, except for a 5 µl aliquot that was taken to assess the extent of fragmentation of the DNA.

### Whole cell lysate (WCL)

For Figure 5B-D, the WCL was fragmented as well by sonicating for 3 cycles 30 seconds on / 30 seconds off in 15 ml bioruptor pico tubes containing 300 µl of sonication beads (Diagenode). After sonication, the sample was transferred a 2 ml Eppendorf tube and the unfragmented chromatin was subsequently pelleted by centrifugation at 18407g (14000 rpm) in an Eppendorf 5424 R centrifuge for 20 minutes at 4°C. The fragmented chromatin was transferred to a 2 ml Eppendorf tube, snap frozen in liquid nitrogen and stored at −80°C, except for a 20 µl aliquot that was taken to assess the extent of fragmentation of the DNA.

### Chromatin isolation using the short bead beating protocol

Chromatin was isolated using the short bead beating protocol for the experiments shown in Figure 5B-5D, 6, 7, 8 and 9. To isolate the chromatin, each cell pellet, which is equivalent to a 100 ml culture of cells OD = 0.8, was resuspended in ~700-900 µl of FA lysis buffer (50 mM HEPES-KOH pH 7.5, 150 mM NaCl, 1 mM EDTA pH 8.0, 1% Triton X-100, 0.1% Na-deoxycholate, 0.1% SDS) containing protease inhibitors: either 1 tablet Roche cOmplete protease inhibitor cocktail (#11836145001) per 25 ml of FA lysis buffer or 30 µl aprotinin (Sigma-Aldrich: #A6279), 1 µl leupeptin (Sigma-Aldrich #L2884, 1 mg/ml in MQ), 1 µl pepstatin A (Sigma-Aldrich #P4265: 1 mg/ ml in 100% Methanol) and 15 µl PMSF (Sigma-Aldrich: #P7626, 200 mM in isopropanol) per milliliter of FA lysis buffer. The resuspended cells were subsequently added to a 2 ml screw-cap tube containing 500 µl of zirconium/ silica beads 0.5 mm (BioSpec Products, #11079105z). If needed, the tubes were closed and inverted once to remove air trapped in the beads. The tubes were then filled completely with FA lysis buffer to remove as much air as possible. The tubes were bead beated 7 times in a genie disruptor for 3 minutes. The samples were put one ice for either 1 or 3 minutes in between each run, depending on the number of samples that were processed at the same time. During the bead beating, 15 ml tubes were prepared containing 1 ml pipette tips with the end cut off. After bead beating, the lysate was recovered by burning a hole in the top and bottom of the screw-cap tubes using a 23G needle and placing them on top of the pipette tip in the 15 ml tube. This combination was then centrifuged at 201g (1000 rpm in an Eppendorf 5810 R) at 4°C and the lysate was transferred to a 2 ml Eppendorf tube. To remove the majority of unbroken cells and cell debris, the lysate was centrifuged at 1503g (4000 rpm) for 2 minutes in an Eppendorf 5424 R centrifuge at 4°C. The supernatant was transferred to a fresh 2 ml Eppendorf tube. Either the whole extract was sonicated in a 15 ml bioruptor pico tube with 300 µl of sonication beads (Diagenode) added (Figure 5B-5D) or 2x 300 µl was sonicated in 2x 1.5 ml bioruptor pico microtubes (Diagenode, Figure 6-9). When using the APLP protease inhibitors, before sonication 4.5 µl aprotinin, 0.15 µl leupeptin, 0.15 µl pepstatin and 3 µl PMSF was added to each tube containing 300 µl chromatin. The samples were sonicated for 3 cycles (Figure 5 and 6A) or 4 cycles (Figure 6C-6F, 7 and 8) 30 seconds on / 30 seconds off or 10 cycles 15 seconds on / 30 seconds off (Figure 9). The chromatin was snap frozen in liquid nitrogen and stored at −80°C, except for a 20 µl aliquot that was taken to assess the extent of fragmentation of the DNA.

### Quality control after sonication

The aliquot that was taken after sonication was diluted to 95 µl by addition of 90 µl (zymolyase protocol and CE, full protocol) or 75 µl (short bead beat protocol and WCL, full protocol) TE/SDS (10 mM Tris-HCl pH 8.0, 1 mM EDTA pH 8.0, 1% SDS) and decross-linked overnight at 65°C. The next morning the decross-linked chromatin was treated with 5 µl of RNAse A/T1 (Thermo Scientific #EN0551) for 30 min at 37°C. Subsequently, proteins were digested by addition of 40 µl proteinase K (Roche #03115852001, dissolved to 10 µg/µl) and incubation for 2 hours at 37°C. The DNA was isolated using a Qiagen PCR purification cleanup kit (Qiagen #28106). The standard cleanup protocol was used, except for the wash with PE buffer, which was done three times using 500 µl PE buffer. The DNA was eluted in 40 µl buffer EB. The degree of DNA fragmentation was assessed by running 1 µl of each sample on a High Sensitivity DNA bioanalyzer chip. The fragmentation was considered good if the peak of the distribution of DNA fragment sizes was between 200-300 bp. If the peak of a specific sample was bigger than 300bp, the chromatin was fragmented for a number of additional cycles to make the distribution similar to that of the other samples.

### Immunoprecipitation using agarose beads

The immunoprecipitation with agarose beads was performed using anti-V5-agarose beads (Sigma-Aldrich #A7345), which are agarose beads that are conjugated to a mono-clonal anti-V5 antibody. The calculated amount of beads for all IPs that were done at the same time (20 µl per IP) was pre-washed together three times in FA lysis buffer (50 mM HEPES-KOH pH 7.5, 150 mM NaCl, 1 mM EDTA pH 8.0, 1% Triton X-100, 0.1% Na-deoxycholate, 0.1% SDS) by resuspending the beads in 1 ml of FA lysis buffer and centrifuging for 1 minute at 845g (3000 rpm) in an Eppendorf 5424 at RT. During the last wash, the beads were divided over 1.5 ml Eppendorf tubes such that there was an equivalent of 20 µl beads per tube. Per IP, the beads were mixed with 250 µl chromatin and incubated for 2 hours at RT. The beads were then pelleted by centrifugation at maximum speed (21130g, 15000 rpm) for a few seconds in an Eppendorf 5424 R centrifuge and the supernatant was removed using a 23G needle and a syringe. The beads were subseqently washed twice in FA lysis buffer, twice with wash buffer 1 (FA lysis buffer containing 0.5 M NaCl), and twice with wash buffer 2 (10 mM Tris pH 8.0, 0.25 mM LiCl, 1 mM EDTA pH 8.0, 0.5% Nonidet P-40 and 0.5% Na-deoxycholate) by centrifugation to maximum speed (21130g, 15000 rpm) for a few seconds. After the last wash, all liquid was removed and the beads were resuspended in 100 µl TE/ SDS (10 mM Tris-HCl pH 8.0, 1 mM EDTA pH 8.0, 1% SDS). 2 µl of RNAse A/T1 (Thermo Scientific #EN0551) was added and the beads were incubated overnight at 65°C in a thermoshaker. The next morning, proteins were digested by addition of 40 µl proteinase K (Roche, dissolved to 10 µg/µl) and incubation for 2 hours at 37°C. After protein digestion, the DNA was recovered using a Qiagen PCR purification cleanup kit (Qiagen #28106). The standard cleanup protocol was used, except that the wash with PE buffer was done three times using 500 µl PE buffer. The DNA was eluted in 40 µl buffer EB.

### Immunoprecipitation using magnetic beads

The anti-V5 antibody (Life Technologies #R96025) was bound to the chromatin by incubating 150 µl (Figure 4B), 200 µl (Figure 2D, 2E, 3 and 5), 250 µl (Figure 2A-2C and 4A) or 450 µl (Figure 5B, 6-9) chromatin with 1 µl (Figure 2D, 2E, 3-9) or 2 µl (Figure 2A-2C) of the anti-V5 antibody, either overnight (Figure 2A-2D and 4A), for 1 hour (Figure 2D), 2 hours (Figure 2D, 2E, 3-9) or 4 hours (Figure 2D) in a rotating wheel at 4°C (Figure 7C and 7C) or RT (Figure 2-9). For the samples with separate protease inhibitors added (Figure 8 and 9), in addition, also 15 µl aprotinin (Sigma-Aldrich #A6279), 0.5 µl leupeptin (Sigma-Aldrich #L2884, 1 mg/ml in MQ), 0.5 µl pepstatin (Sigma-Aldrich #P4265: 1 mg/ml in 100% Methanol) and 5 µl PMSF (Sigma-Aldrich: #P7626, 200 mM in isopropanol) were added right before the incubation.

During the incubation of chromatin with the antibody, the magnetic beads were prepared. For each IP, either 50 µl (Figure 2A-2C, 7 and 8) or 25 µl (Figure 2D, 2E, 3-7 and 9) magnetic beads (Dynabeads protein G, Life Technologies #10004D) was used. The preparation was done in either one of two ways. For the experiments shown in Figure 2A-2D and 4A the amount of beads was taken for the number of IPs + 0.5 and transferred to a 1.5 ml Eppendorf tube. The beads were separated on a magnetic stand (DynaMag-2, Life Technologies #12321D) and the supernatant was removed. The beads were washed once in PBS-T (PBS pH 7.4, 0.02% Tween-20) by pipetting up and down. The beads were then divided over 1.5 ml Eppendorf tubes, separated using a magnetic stand and the supernatant was removed. When the beads were pre-incubated with BSA (Figure 2E, 3, 4B, 5-9) the beads were prepared as follows. 25 µl (Figure 2E, 3, 4B, 5-7 and 9) or 50 µl (Figure 8) of magnetic beads per IP were transferred to a 1.5 ml Eppendorf tube and separated on a magnetic stand. The supernatant was removed and the beads were resuspended in 500 µl of PBS-T by gentle vortexing. After separating on the magnetic stand and removal of the supernatant, the beads were resuspended in 200 µl PBS + 12.5 µl of BSA (10 mg/ml in TBS-T: 150 mM NaCl, 10 mM Tris pH 7.5, 0.05% Tween-20, Figure 2E, 3, 4B, 5 and 6A), or 400 µl PBS + 25 µl of BSA (Figure 6C-6F, 7-9) and incubated at 4°C until the chromatin + anti-V5 antibody incubation was finished. 10 minutes before the end of the incubation of the chromatin with the antibody, the beads were centrifuged for 5 seconds to collect them in the bottom of the tube and subsequently separated on a magnetic stand. The supernatant was removed and the beads were washed once with 500 µl PBS-T. After removing the supernatant, the chromatin + antibody was added to the beads. For the experiments shown in Figure 8 and 9, in addition, 5 µl of PMSF was added. The beads, chromatin and antibody combination was resuspended and incubated at RT (Figure 2-9) or 4°C (Figure 7C and 7D) for 20 minutes (Figure 2-9) or 60 minutes (Figure 7C and 7D) in a rotating wheel. For the “pre incubation Ab + beads” samples shown in Figure 2C, the beads were incubated overnight with 2 µl anti-V5 antibody in 200 µl PBS-T at 4°C after washing. The next day, the supernatant was removed and the antibody conjugated beads were incubated with the chromatin in a rotating wheel for 20 minutes at RT.

After the incubation of the beads with the chromatin, the beads were washed by separating them on a magnetic stand and resuspended using either gentle pipetting (Figure 2A-2D, 4A) or gentle vortexing (Figure 2E, 3, 4B, 5-9). The beads were washed with any of the washes: twice with PBS and once with PBS-T (Figure 2-9, standard wash), once with PBS (Figure 7B: low wash), twice with FA-lysis buffer, twice with wash buffer 1 (FA lysis buffer containing 0.5 M NaCl) and twice with wash buffer 2 (10 mM Tris pH 8.0, 0.25 mM LiCl, 1 mM EDTA pH 8.0, 0.5% Nonidet P-40 and 0.5% Na-deoxycholate) (Figure 7B: high wash 1) or twice with PBS, twice with wash buffer 1 and twice with PBS-T (Figure 7B: high wash 2). The beads were then resuspended in 100 µl PBS-T and transferred to a fresh 1.5 ml DNA LoBind Tube (Eppendorf #0030108051). After removal of the PBS-T, the cells were resuspended in 95 µl (Figure 2A-2C) or 98 µl (Figure 2D, 2E, 3-9) TE/SDS (10 mM Tris–HCl pH 8.0, 1 mM EDTA pH 8.0, 1% SDS) and decross-linked at 65°C overnight in a thermoshaker. As an input control, 5 µl (Figure 2-5), 10 µl (Figure 7), 11.25 µl (Figure 5B) or 20 µl (Figure 6-9) of the chromatin that was not incubated with the antibody was also decross-linked overnight after dilution to 95 µl with TE/SDS at 65°C in a thermoshaker. For the glycine eluted samples (Figure 2B), the beads were first resuspended in 20 µl 50 mM glycine pH 2.8 and incubated for 2 minutes. The beads were separated and the supernatant was mixed with 75 µl TE/SDS, and subsequently the beads were resuspended in 95 µl of TE/ SDS. Both were decross-linked overnight at 65°C.

The next morning, RNA was first degraded by addition of 2 µl (IP: Figure 2D, 2E, 3-9) or 5 µl (IP: Figure 2A-2C and input: Figure 2-9) RNAse A/T1 (Thermo Scientific #EN0551) and incubation the beads at 37°C for 30 minutes. Subsequently, proteins were digested by addition of 40 µl proteinase K (Roche #03115852001, dissolved to 10 µg/µl) and incubation for 2 hours at 37°C. After protein digestion, the DNA was recovered using a Qiagen PCR purification cleanup kit (Qiagen #28106). The standard cleanup protocol was used, except that the wash with PE buffer was done three times using 500 µl PE buffer. The DNA was eluted in 30 µl (Figure 2, 4A) or 40 µl (Figure 3, 4B, 5-9) buffer EB.

### Western blot

The integrity of the proteins of interest was assessed using Western blots at different steps during the protocol. 5 µl (Figure 9), 20 µl (Figure 4, 5) or 40 µl (Figure 8) aliquots were taken at the indicated steps, snap frozen and stored at −80°C. The pre-lysis samples and the control samples were lysed by resuspending the cells in 200 µl of 0.1 M NaOH and incubating them for 5 minutes at RT. After centrifugation at 3381g (6000 rpm) in an 5424 Eppendorf centrifuge at RT, the supernatant was discarded and the cell pellet was resuspended in 2x sample buffer (2% SDS, 80 mM Tris pH 6.8, 10 % glycerol, 572 mM β-mercaptoethanol, 0.016% bromophenol blue). These samples were heated for 5 minutes (non-cross-linked controls) or 30 minutes (cross-linked, pre-lysis samples) at 95°C before loading 10 µl on a protein gel.

The post-lysis and post-sonication samples were diluted with 5x sample buffer (5% SDS, 200 mM Tris pH 6.8, 25% glycerol, 1.43 M β-mercaptoethanol, 0.032% bromophenol blue). 10 µl of sample was mixed with 2.5 µl of 5x sample buffer (Figure 9), 20 µl was mixed with 20 µl of 5x sample buffer (Figure 4 and 5) or 40 µl was mixed with 10 µl 5x sample buffer (Figure 8). The cross-linked samples were decross-linked for 30 minutes at 95°C before loading 10 µl on the gel. Either regular (Figure 4 and 5) or stain-free (Figure 8 and 9, Bio-Rad #1610182) 10 % acrylamide gels were used. 5 µl of a broad range protein marker (New England Biolabs, #P7708) was run alongside the samples on each gel to estimate the molecular weight. After the samples migrated through the gel, the proteins were transferred to a nitrocellulose membrane (Amersham Protran nitrocellulose membrane #10600001) overnight in transfer buffer containing 10% methanol. After transfer, the membrane was first washed with TBS-T (150 mM NaCl, 10 mM Tris pH 7.5, 0.05% Tween-20) and then blocked in TBS-T containing 2% protifar (Nutricia #56317) either for 1 hour at RT or overnight at 4°C. After two washes with TBS-T, the membrane was incubated with an anti-V5 antibody (Life Technologies #R96025) and incubated for 1.5 hours at RT. The blot was washed 4 times with TBS-T and subsequently incubated with a goat anti-mouse-HRP antibody (Bio-Rad #1706516) for 30 minutes. The blot was again washed 3 times with TBS-T and the last wash was performed using TBS. The blot was then imaged after incubation with ECL (PerkinElmer, Western Lightning Plus #NEL105001EA) on a chemidoc touch imaging system (Bio-Rad).

The quantification of the percentage of intact protein was performed as follows. An image of each blot was taken without any satured pixels. The image was exported for analysis as a TIFF image using the Image Lab software (Bio-Rad) and analyzed in the ImageJ software (Schneider *et al*, 2012). Using the rectangle tool, a rectangle was drawn around each intact protein band and the sum of the pixels was recorded. The same sized rectangle was used for each intact protein band on the same gel. The amount of degraded signal was determined by drawing a rectangle that encompassed all the visible lower bands and recording the sum of the pixels in this rectangle. Since each protein had a different number of degradation bands, a differently sized rectangle was used for the degraded bands of each protein. The signal of the intact and degraded bands was background corrected by subtracting the sum of the pixels of an identically shaped rectangle that was drawn somewhere on the blot where there were no protein bands present. The percent intact protein was then calculated by dividing the signal from the intact protein band by the sum of the signals from the intact and the degraded protein bands. This is represented by the following formula:

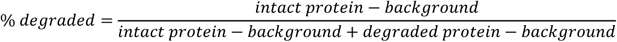

### qPCR

DNA binding levels were quantified using qPCR. qPCRs were performed in 384-well plates (Bio-Rad white hard-shell plate #hsp3805) in a final volume of 10 µl. For each reaction, 5 µl IQ SYBR Green super mix (Bio-Rad #1708886), 2.8 µl MQ, 0.2 µl primer mix (containing 10 µM of both the forward and the reverse primer), 0.014 µl precision blue (Bio-Rad #1725555) and 2 µl template were mixed. First, a master mix was made of MQ and IQ SYBR Green for the total number of reactions + ~20%. Subsequently, a separate master mix was made for each primer pair for the number of reactions for each primer pair + ~10%. After addition of the primers, precision blue was added to this master mix using a 1:700 dilution, to aid pipetting in the 384-well plate. An electronic pipette was used to accurately pipette 8 µl of this mix to the wells of the 384-well plate. Next, 2 µl template or MQ was added to each well using an 8-channel multichannel pipette.

On each plate, a 5 times, 10-fold serially diluted standard curve was taken along for each primer pair, together with at least 2 no template controls (MQ). All IP measurements were performed either undiluted or 2 times diluted and were performed in technical quadruplicate. The inputs were diluted 10, 25, 50 or 100 times, depending on the experiment, and were either done in technical triplicate or quadruplicate. qPCRs were performed using a CFX384 Touch Real-Time PCR Detection system (Bio-Rad) with a 2-step PCR protocol. First, the DNA was denatured for 30 seconds at 95°C, followed by 40 cycles of 15 seconds denaturing at 95°C and 30 seconds of amplification at 60°C. The amplification was followed by a melting curve that started with 1 minute denaturing at 95°C and then ramped up from 70°C with half a degree every 5 seconds to 95°C. All primers were designed to have a melting temperature above 60°C in the IQ SYBR green super mix.

In all figures the binding is shown as a percentage of input. This was calculated by first calculating the starting quantities (SQ) for each IP and input for each primer pair using the CFX Meastro (Bio-Rad) software. These values were then exported and the average value of the replicates was calculated. If there was a single measurement that was too high or low due to a pipetting error, this measurement was removed before calculating the average. The starting quantities were first corrected for the dilution and subsequently for volume, since for the inputs only a small fraction of the material was used compared to the IP (~5%). After this correction, the signal from the IP was divided by the signal from the input for each biological replicate, which yields the percent of input. The average of the biological replicates is shown in the figures. The variation was calculated by using the distance from the mean when duplicates were used or by using the standard deviation when more than two replicates were used.

## Supporting information

Optimized ChIP protocol

Methods per figure plus supplemental tables

## Acknowledgements

We thank the members of the Holstege and Kemmeren groups for their support and discussions. We thank Tineke Lenstra for advice regarding the protocol. We are grateful to Jeff DeMartino for critical reading of the manuscript. This work was supported by the Netherlands Organisation for Scientific Research (NWO) grant 86411010 and by the European Research Council (ERC) grant 671174 DynaMech.

## References

Acevedo LG, Leonardo Iniguez A, Holster HL, Zhang X, Green R, Farnham PJ (2007) Genome-scale ChIP-chip analysis using 10,000 human cells. BioTechniques 43: 791–797

Adli M, Zhu J, Bernstein BE (2010) Genome-wide chromatin maps derived from limited numbers of hematopoietic progenitors. Nat Methods 7: 615–618

Arrigoni L, Al-Hasani H, Ramírez F, Panzeri I, Ryan DP, Santacruz D, Kress N, Pospisilik JA, Bönisch U, Manke T (2018) RELACS nuclei barcoding enables high-throughput ChIP-seq. Commun Biol 1: 214

Arrigoni L, Richter AS, Betancourt E, Bruder K, Diehl S, Manke T, Bönisch U (2016) Standardizing chromatin research: a simple and universal method for ChIP-seq. Nucleic Acids Res 44: e67

Baranello L, Kouzine F, Sanford S, Levens D (2015) ChIP bias as a function of cross-linking time. Chromosome Res 24: 175–181

Brind’Amour J, Liu S, Hudson M, Chen C, Karimi MM, Lorincz MC (2015) An ultra-low-input native ChIP-seq protocol for genome-wide profiling of rare cell populations. Nat Commun 6: 6033

Buchman AR, Kornberg RD (1990) A yeast ARS-binding protein activates transcription synergistically in combination with other weak activating factors. Mol Cell Biol 10: 887–897

Bulyk ML, Huang X, Choo Y, Church GM (2001) Exploring the DNA-binding specificities of zinc fingers with DNA microarrays. Proc Natl Acad Sci U S A 98: 7158–7163

Cai M, Davis RW (1990) Yeast centromere binding protein CBF1, of the helix-loop-helix protein family, is required for chromosome stability and methionine prototrophy. Cell 61: 437–446

Chasman DI, Lue NF, Buchman AR, LaPointe JW, Lorch Y, Kornberg RD (1990) A yeast protein that influences the chromatin structure of UASG and functions as a powerful auxiliary gene activator. Genes Dev 4: 503–514

Cremazy FGE, Manders EMM, Bastiaens PIH, Kramer G, Hager GL, van Munster EB, Verschure PJ, Gadella TJ, van Driel R (2005) Imaging in situ protein-DNA interactions in the cell nucleus using FRET-FLIM. Exp Cell Res 309: 390–396

Dahl JA, Collas P (2007) Q2ChIP, a Quick and Quantitative Chromatin Immunoprecipitation Assay, Unravels Epigenetic Dynamics of Developmentally Regulated Genes in Human Carcinoma Cells. Stem Cells 25: 1037–1046

Dedon PC, Soults JA, David Allis C, Gorovsky MA (1991) A simplified formaldehyde fixation and immunoprecipitation technique for studying protein-DNA interactions. Anal Biochem 197: 83–90

Dey B, Thukral S, Krishnan S, Chakrobarty M, Gupta S, Manghani C, Rani V (2012) DNA-protein interactions: methods for detection and analysis. Mol Cell Biochem 365: 279–299

Ellington AD, Szostak JW (1990) In vitro selection of RNA molecules that bind specific ligands. Nature 346: 818

Field J, Nikawa J, Broek D, MacDonald B, Rodgers L, Wilson IA, Lerner RA, Wigler M (1988) Purification of a RAS-responsive adenylyl cyclase complex from Saccharomyces cerevisiae by use of an epitope addition method. Mol Cell Biol 8: 2159–2165

Galas DJ, Schmitz A (1978) DNAse footprinting: a simple method for the detection of protein-DNA binding specificity. Nucleic Acids Res 5: 3157–3170

Ganapathi M, Palumbo MJ, Ansari SA, He Q, Tsui K, Nislow C, Morse RH (2011) Extensive role of the general regulatory factors, Abf1 and Rap1, in determining genome-wide chromatin structure in budding yeast. Nucleic Acids Res 39: 2032–2044

Garner MM, Revzin A (1981) A gel electrophoresis method for quantifying the binding of proteins to specific DNA regions: application to components of the Escherichia coli lactose operon regulatory system. Nucleic Acids Res 9: 3047–3060

Ghaemmaghami S, Huh W-K, Bower K, Howson RW, Belle A, Dephoure N, O’Shea EK, Weissman JS (2003) Global analysis of protein expression in yeast. Nature 425: 737–741

Gilmour DS, Lis JT (1984) Detecting protein-DNA interactions in vivo: distribution of RNA polymerase on specific bacterial genes. Proc Natl Acad Sci U S A 81: 4275–4279

Goren A, Ozsolak F, Shoresh N, Ku M, Adli M, Hart C, Gymrek M, Zuk O, Regev A, Milos PM, Bernstein BE (2010) Chromatin profiling by directly sequencing small quantities of immunoprecipitated DNA. Nat Methods 7: 47–49

Gutin J, Sadeh R, Bodenheimer N, Joseph-Strauss D, Klein-Brill A, Alajem A, Ram O, Friedman N (2018) Fine-Resolution Mapping of TF Binding and Chromatin Interactions. Cell Rep 22: 2797–2807

Haber JE (2012) Mating-Type Genes and MAT Switching in Saccharomyces cerevisiae. Genetics 191: 33–64

Hartley PD, Madhani HD (2009) Mechanisms that specify promoter nucleosome location and identity. Cell 137: 445–458

Haruki H, Nishikawa J, Laemmli UK (2008) The anchor-away technique: rapid, conditional establishment of yeast mutant phenotypes. Mol Cell 31: 925–932

He Q, Johnston J, Zeitlinger J (2015) ChIP-nexus enables improved detection of in vivo transcription factor binding footprints. Nat Biotechnol 33: 395–401

Hebbes TR, Thorne AW, Crane-Robinson C (1988) A direct link between core histone acetylation and transcriptionally active chromatin. EMBO J 7: 1395–1402

Hoffman EA, Frey BL, Smith LM, Auble DT (2015) Formaldehyde Crosslinking: A Tool for the Study of Chromatin Complexes. J Biol Chem 290: 26404–26411

Hopp TP, Prickett KS, Price VL, Libby RT, March CJ, Cerretti DP, Urdal DL, Conlon PJ (1988) A Short Polypeptide Marker Sequence Useful for Recombinant Protein Identification and Purification. Bio/Technology 6: 1204

Irlbacher H, Franke J, Manke T, Vingron M, Ehrenhofer-Murray AE (2005) Control of replication initiation and heterochromatin formation in Saccharomyces cerevisiae by a regulator of meiotic gene expression. Genes Dev 19: 1811–1822

Jackson V (1978) Studies on histone organization in the nucleosome using formaldehyde as a reversible cross-linking agent. Cell 15: 945–954

de Jonge WJ, O’Duibhir E, Lijnzaad P, van Leenen D, Groot Koerkamp MJ, Kemmeren P, Holstege FC (2017) Molecular mechanisms that distinguish TFIID housekeeping from regulatable SAGA promoters. EMBO J 36: 274–290

Ju QD, Morrow BE, Warner JR (1990) REB1, a yeast DNA-binding protein with many targets, is essential for growth and bears some resemblance to the oncogene myb. Mol Cell Biol 10: 5226–5234

Kasinathan S, Orsi GA, Zentner GE, Ahmad K, Henikoff S (2014) High-resolution mapping of transcription factor binding sites on native chromatin. Nat Methods 11: 203–209

Kawashima Y, Kodera Y, Singh A, Matsumoto M, Matsumoto H (2014) Efficient extraction of proteins from formalin-fixed paraffin-embedded tissues requires higher concentration of tris(hydroxymethyl)aminomethane. Clin Proteomics 11: 4

Kemmeren P, Sameith K, van de Pasch LAL, Benschop JJ, Lenstra TL, Margaritis T, O’Duibhir E, Apweiler E, van Wageningen S, Ko CW, van Heesch S, Kashani MM, Ampatziadis-Michailidis G, Brok MO, Brabers NACH, Miles AJ, Bouwmeester D, van Hooff SR, van Bakel H, Sluiters E, et al(2014) Large-scale genetic perturbations reveal regulatory networks and an abundance of gene-specific repressors. Cell 157: 740–752

Kent NA, Eibert SM, Mellor J (2004) Cbf1p is required for chromatin remodeling at promoter-proximal CACGTG motifs in yeast. J Biol Chem 279: 27116–27123

Kuo MH, Allis CD (1999) In vivo cross-linking and immunoprecipitation for studying dynamic Protein:DNA associations in a chromatin environment. Methods San Diego Calif 19: 425–433

Kuo MH, Grayhack E (1994) A library of yeast genomic MCM1 binding sites contains genes involved in cell cycle control, cell wall and membrane structure, and metabolism. Mol Cell Biol 14: 348–359

Lara-Astiaso D, Weiner A, Lorenzo-Vivas E, Zaretsky I, Jaitin DA, David E, Keren-Shaul H, Mildner A, Winter D, Jung S, Friedman N, Amit I (2014) Immunogenetics. Chromatin state dynamics during blood formation. Science 345: 943–949

Lushnikov AY, Potaman VN, Oussatcheva EA, Sinden RR, Lyubchenko YL (2006) DNA strand arrangement within the SfiI-DNA complex: atomic force microscopy analysis. Biochemistry 45: 152–158

MacIsaac KD, Wang T, Gordon DB, Gifford DK, Stormo GD, Fraenkel E (2006) An improved map of conserved regulatory sites for Saccharomyces cerevisiae. BMC Bioinformatics 7: 113

Maerkl SJ, Quake SR (2007) A systems approach to measuring the binding energy landscapes of transcription factors. Science 315: 233–237

McIsaac RS, Petti AA, Bussemaker HJ, Botstein D (2012) Perturbation-based analysis and modeling of combinatorial regulation in the yeast sulfur assimilation pathway. Mol Biol Cell 23: 2993–3007

Messenguy F, Dubois E (2003) Role of MADS box proteins and their cofactors in combinatorial control of gene expression and cell development. Gene 316: 1–21

Miyake T, Reese J, Loch CM, Auble DT, Li R (2004) Genome-wide analysis of ARS (autonomously replicating sequence) binding factor 1 (Abf1p)-mediated transcriptional regulation in Saccharomyces cerevisiae. J Biol Chem 279: 34865–34872

Nick H, Gilbert W (1985) Detection in vivo of protein-DNA interactions within the lac operon of Escherichia coli. Nature 313: 795–798

O’Neill LP, Turner BM (2003) Immunoprecipitation of native chromatin: NChIP. Methods San Diego Calif 31: 76–82

O’Neill LP, VerMilyea MD, Turner BM (2006) Epigenetic characterization of the early embryo with a chromatin immunoprecipitation protocol applicable to small cell populations. Nat Genet 38: 835–841

Park D, Lee Y, Bhupindersingh G, Iyer VR (2013) Widespread misinterpretable ChIP-seq bias in yeast. PloS One 8: e83506

Passmore S, Maine GT, Elble R, Christ C, Tye BK (1988) Saccharomyces cerevisiae protein involved in plasmid maintenance is necessary for mating of MAT alpha cells. J Mol Biol 204: 593–606

Pchelintsev NA, Adams PD, Nelson DM (2016) Critical Parameters for Efficient Sonication and Improved Chromatin Immunoprecipitation of High Molecular Weight Proteins. PloS One 11: e0148023

Poorey K, Viswanathan R, Carver MN, Karpova TS, Cirimotich SM, McNally JG, Bekiranov S, Auble DT (2013) Measuring Chromatin Interaction Dynamics on the Second Time Scale at Single-Copy Genes. Science 342: 369–372

Rando OJ (2010) Genome-wide mapping of nucleosomes in yeast. Methods Enzymol 470: 105–118

Reed SH, Akiyama M, Stillman B, Friedberg EC (1999) Yeast autonomously replicating sequence binding factor is involved in nucleotide excision repair. Genes Dev 13: 3052–3058

Rhee HS, Pugh BF (2011) Comprehensive genome-wide protein-DNA interactions detected at single-nucleotide resolution. Cell 147: 1408–1419

Rhode PR, Elsasser S, Campbell JL (1992) Role of multifunctional autonomously replicating sequence binding factor 1 in the initiation of DNA replication and transcriptional control in Saccharomyces cerevisiae. Mol Cell Biol 12: 1064–1077

Rhode PR, Sweder KS, Oegema KF, Campbell JL (1989) The gene encoding ARS-binding factor I is essential for the viability of yeast. Genes Dev 3: 1926–1939

Schmid M, Durussel T, Laemmli UK (2004) ChIC and ChEC; genomic mapping of chromatin proteins. Mol Cell 16: 147–157

Schmidl C, Rendeiro AF, Sheffield NC, Bock C (2015) ChIPmentation: fast, robust, low-input ChIP-seq for histones and transcription factors. Nat Methods 12: 963–965

Schneider CA, Rasband WS, Eliceiri KW (2012) NIH Image to ImageJ: 25 years of image analysis. Nat Methods 9: 671–675

Siebenlist U, Gilbert W (1980) Contacts between Escherichia coli RNA polymerase and an early promoter of phage T7. Proc Natl Acad Sci U S A 77: 122–126

Siebenlist U, Simpson RB, Gilbert W (1980) E. coli RNA polymerase interacts homologously with two different promoters. Cell 20: 269–281

Skene PJ, Henikoff S (2015) A simple method for generating high-resolution maps of genome-wide protein binding. eLife 4: e09225

Skene PJ, Henikoff S (2017) An efficient targeted nuclease strategy for high-resolution mapping of DNA binding sites. eLife 6: e21856

Smith GP (1985) Filamentous fusion phage: novel expression vectors that display cloned antigens on the virion surface. Science 228: 1315–1317

Solomon MJ, Larsen PL, Varshavsky A (1988) Mapping proteinDNA interactions in vivo with formaldehyde: Evidence that histone H4 is retained on a highly transcribed gene. Cell 53: 937–947

Southern JA, Young DF, Heaney F, Baumgärtner WK, Randall RE (1991) Identification of an epitope on the P and V proteins of simian virus 5 that distinguishes between two isolates with different biological characteristics. J Gen Virol 72: 1551–1557

van Steensel B, Henikoff S (2000) Identification of in vivo DNA targets of chromatin proteins using tethered dam methyltransferase. Nat Biotechnol 18: 424–428

Sutherland BW, Toews J, Kast J (2008) Utility of formaldehyde cross-linking and mass spectrometry in the study of protein-protein interactions. J Mass Spectrom JMS 43: 699–715

Teytelman L, Thurtle DM, Rine J, van Oudenaarden A (2013) Highly expressed loci are vulnerable to misleading ChIP localization of multiple unrelated proteins. Proc Natl Acad Sci U S A 110: 18602–18607

Tuerk C, Gold L (1990) Systematic evolution of ligands by exponential enrichment: RNA ligands to bacteriophage T4 DNA polymerase. Science 249: 505–510

Wallerman O, Nord H, Bysani M, Borghini L, Wadelius C (2015) lobChIP: from cells to sequencing ready ChIP libraries in a single day. Epigenetics Chromatin 8: 25

Wang MM, Reed RR (1993) Molecular cloning of the olfactory neuronal transcription factor Olf-1 by genetic selection in yeast. Nature 364: 121–126

Xie J, Pierce M, Gailus-Durner V, Wagner M, Winter E, Vershon AK (1999) Sum1 and Hst1 repress middle sporulation-specific gene expression during mitosis in Saccharomyces cerevisiae. EMBO J 18: 6448–6454

Yin H, Wang MD, Svoboda K, Landick R, Block SM, Gelles J (1995) Transcription against an applied force. Science 270: 1653–1657

Zaidi H, Hoffman EA, Shetty SJ, Bekiranov S, Auble DT (2017) Second-generation method for analysis of chromatin binding with formaldehyde–cross-linking kinetics. J Biol Chem 292: 19338–19355

Zaret KS, Mango SE (2016) Pioneer transcription factors, chromatin dynamics, and cell fate control. Curr Opin Genet Dev 37: 76–81

Zentner GE, Kasinathan S, Xin B, Rohs R, Henikoff S (2015) ChEC-seq kinetics discriminates transcription factor binding sites by DNA sequence and shape in vivo. Nat Commun 6: 8733

