## Supplementary material for "An extensively optimized chromatin immunoprecipitation protocol for quantitatively comparable and robust results": Optimized ChIP protocol

### Solutions needed for the ChIP protocol

| | |
|---|---|
| Formaldehyde | 37% formaldehyde containing 10-15% methanol (Sigma-Aldrich #252549)<br>Never use > 3 months old, preferably less than 1 month old. |
| Tris | 4.5 M Tris pH 8.0 |
| TBS | 150 mM NaCl<br>10 mM Tris pH 7.5 |
| FA lysis buffer | 50 mM HEPES-KOH pH 7.5<br>150 mM NaCl<br>1 mM EDTA pH 8.0<br>1% Triton X-100<br>0.1% Na-deoxycholate<br>0.1% SDS |
| Pepstatin A | 1 mg/ml leupeptin in 100% methanol (1.51 mM). Store at -20°C. |
| Leupeptin | 1 mg/ml pepstatin in MQ (2.10 mM). Store at -20°C. |
| PMSF | 200 mM Phenylmethanesulfonyl fluoride (PMSF) in isopropanol.<br>Heat to 37°C to dissolve and store at-20°C. |
| RNAse A/T1 | RNAse A 2mg/ml & RNase T1 5000 U/ml mix (Thermo Scientific #EN0551) |
| Proteinase K | 10 μg/μl in TE (10 mM Tris pH 8, 1 mM EDTA pH 8) |
| PBS | 137 mM NaCl<br>2.7 mM KCl<br>10 mM $Na_2HPO_4$<br>1.47 mM $KH_2PO_4$<br>1 mM CaCl<br>0.5 mM MgCl<br>pH adjusted to 7.3 using HCl |
| PBS-T | PBS + 0.02% Tween-20 |
| TE/SDS | 10 mM Tris pH 8<br>1 mM EDTA pH 8<br>1% SDS |
| BSA in TBS-T | Dissolved to 10 mg/ml in TBS-T (150 mM NaCl, 10 mM Tris pH 7.5, 0.05% Tween-20) |
| 5X sample buffer (optional) | 5% SDS<br>200 mM Tris pH 6.8<br>25% glycerol<br>1.43 M β-mercaptoethanol<br>0.032% bromophenol blue) |

| | |
|---|---|
| ChIP wash buffer 1 (optional) | 50 mM HEPES-KOH pH 7.5<br>500 mM NaCl<br>1 mM EDTA pH 8.0<br>1% Triton X-100<br>0.1% Na-deoxycholate<br>0.1% SDS<br>(FA lysis with 500 mM NaCl)<br>When adding protease inhibitors (fresh on that day) per 1 ml:<br>30 µl Aprotinin, 1 µl Pepstatin, 1 µl Leupeptin and 10 µl PMSF |
| ChIP wash buffer 2 (optional) | 10 mM Tris pH 8.0<br>0.25 M LiCl<br>0.5% Nonidet P-40<br>0.5% Na-deoxycholate<br>1 mM EDTA pH 8.0<br>When adding protease inhibitors (fresh on that day) per 1 ml:<br>30 µl Aprotinin, 1 µl Pepstatin, 1 µl Leupeptin and 10 µl PMSF |

**1. Growth and *in vivo* cross-linking – Day 1**

1. *Day -3.* Streak the strains of interest on appropriate selection plates and incubate for 3 days @ 30°C.
2. *Day 0.* In the morning: for each strain/condition pick several colonies from a fresh plate and inoculate each colony in 1.5 ml SC-medium in a 24-well plate. Grow the cultures with shaking (230 rpm) at 30°C. At the end of the day combine (1.5 ml) with 13.5 ml warm SC medium for an o/n culture of 15 ml in a 100 ml flask.
    - All incubations are performed at 30°C and 230 rpm.
    - If more starting culture is required it is possible to combine several (3) cultures and make an overnight culture of 20 ml by adding 3x 1.5 ml to 15.5 ml of SC.
3. *Day 1.* Measure (1:50) and dilute cultures in 110 ml pre warmed SC-medium to an OD of 0.20 in a 500 ml Erlenmeyer, mix well and measure OD.
    - There is an extra 10 ml for sampling OD. Remove this, to make sure that there is exactly 100 ml left, before the addition of formaldehyde.
4. Grow yeast until OD 0.8 (2 doublings), this is equivalent to approximately $2*10^7$ cells/ml.
5. Add 5.7 ml of 37% formaldehyde to a final concentration of 2%. Incubate outside the stove on a heated (30°C) stir plate under agitation/stirring (using a stir bar) for 5 min.
    - It's useful to make aliquots (in 50 ml tubes) of formaldehyde and Tris in advance. To add, just pour the formaldehyde in the culture. Be careful to pour it straight into the culture without touching the walls of the Erlenmeyer.
    - Never use formaldehyde that is > 3 months old, preferable < 1 month old.
6. Add Tris 4.5 M to a final concentration of 1.5 M by adding 52.9 ml to stop the cross-linking reaction, and incubate for 1 min using the same agitation as the cross-linking.
7. Split the cultures over 3x 50 ml tubes and spin at 3220g (4000 rpm) for 3 min at 4°C (Eppendorf 5810 R).
8. Wash pellet by resuspending the first pellet in 10 ml ice-cold TBS pH 7.5, combine the three pellets and spin down for 3 min at 3220g (4000 rpm) at 4°C (Eppendorf 5810 R).
9. Remove supernatant, resuspend pellet in 1 ml ice-cold MQ, and transfer to a 2 ml safe-lock tube.
10. Spin down at 3381g (6000 rpm) for 20 sec in a centrifuge (Eppendorf 5424).
11. Remove supernatant, freeze pellet in liquid nitrogen and store at –80°C.

**2. Short bead beating procedure – Day 2**

This protocol is for 100 ml of mid-log cells, OD = 0.8. Preferably process 6 samples at the same time to increase speed and prevent proteolytic degradation of the samples.

- Keep the samples on ice all the time and pre-chill all tubes on ice! Work as fast as possible to prevent degradation.
- The FA lysis buffer is kept on ice and four different protease inhibitors are added, ~1.9 ml FA lysis buffer per sample is needed. Add the following protease inhibitors:

| **Name** | **Amount to add per ml** | **Note** |
|---|---|---|
| **Aprotinin** | 30 µl | Stock at 4°C 33.3x |
| **Leupeptin** | 1 µl | Stock in MQ -20°C, 1000x |
| **Pepstatin A** | 1 µl | Stock in Methanol -20°C, 1000x |
| **PMSF** | 10 µl | Stock in isopropanol -20°C, 100x |

**IMPORTANT**: PMSF is a highly toxic neurotoxin. Be very careful when preparing the stock, and always add this in a fume hood.

12. Add 500 µl of zirconium beads (0.5 µm) to 2.0 ml screw-cap tubes. Measure the right amount of beads by using a 0.5 ml Eppendorf tube.
    - Pre-cool both the centrifuges (Eppendorf 5810 R and 5424 R).
13. Add the protease inhibitors to the FA lysis buffer right before you add the buffer to the cells. For 6 pellets add to 14.5 ml of FA lysis buffer: 450 µl aprotinin, 15 µl pepstatin, 15 µl leupeptin first and then 150 µl PMSF.
    - Be careful with the PMSF, this is a dangerous neurotoxin: add in the fume hood. To add, take the stock out of -20°C and warm the tube by rubbing it between your hands. PMSF crystallizes when it is cold so you need to re-solubilize it. Alternate between a gentle vortex and rubbing until re-solubilized. When you add the PMSF, **mix immediately** to prevent precipitation. Add PMSF last as it will lose activity in an aqueous environment rapidly.
14. Resuspend the frozen pellet carefully in ~900 µl the FA lysis buffer with protease inhibitors and transfer to the 2.0 ml screw-cap tube with beads very slowly, to prevent any air getting trapped in the beads.
    - Optional: take a 5 µl sample for western here (pre-bead beat).
    - If there is any air trapped in the beads: close tube and turn it upside down to release the air that is trapped between the beads.
15. Add more FA lysis buffer containing protease inhibitors to fill the tube completely, keeping as little air as possible before bead beating.
16. Disrupt cells with a genie disrupter at 4°C by bead beating 7 times 3 min. Put samples on ice for 1 min between each run.
    - During this time pre-chill the sonicator and prepare/label tubes!
17. Prepare 15 ml tubes containing a 1 ml pipet tip (cut the end of the 1 ml tips to avoid blockage).
18. Recover each extract by burning a hole in the bottom of the screw-cap tube with a hot 23G needle and quickly placing it in the corning tube on top of the 1 ml pipet tip and burn a second hole in the top of the tube to facilitate release. Centrifuge this combination at 201g (1000 rpm) for 1 minute at 4°C (Eppendorf 5810 R).
19. Transfer the complete lysates to a 2.0 ml Eppendorf tube and spin down at 1503g (4000 rpm) for 2 min at 4°C (Eppendorf 5424 R).
    - This step will remove the majority of the cell debris that could interfere in the subsequent sonication.
    - With long cross-linking times it is possible to see the chromatin as a vague white layer of the supernatant.
20. Transfer all supernatant to a new 2.0 ml Eppendorf tube (take as much as possible (~1400 µl))

- With long cross-linking times, a part of the chromatin may co-precipitate with the pellet and form a transparent white layer above the cells. Make sure to transfer this as well.

21. Transfer 2x 300 µl to 1.5 ml bioruptor pico sonication tubes (300 µl per tube, this is about half of the total volume) and shear for 10 (0 min or 5 min cross-linking) or 8 (10 min or 20 min cross-linking) cycles 15'' on, 30'' off in a bioruptor pico that is connected to a water cooler, which is set to 4°C.
    - Before sonication, add again fresh protease inhibitors. Make a mix of 63 µl Aprotinin, 2.1 µl Leupeptin and 2.1 µl Pepstatin. Mix and add 4.8 µl to all 300 µl samples in sonication tubes first and then add 3 µl of PMSF (in fumehood), mix immediately by very gentle vortexing, making sure the extract does not touch the lid!
22. Spin down sample for 20 min at max speed (21130g, 15000 rpm, 5424 R Eppendorf) at 4°C.
23. Combine the two supernatants per sample in a new 2.0 ml Eppendorf tube.
24. Take 20 µl for QC and take 1 µl for Bradford (dilute 1:41 by adding it to 40 µl of MQ). Optional: also take 10 µl for a Western, add 2.5 µl of 5x sample buffer and boil for 30 min @ 95°C. (For the Western load 10 µl on the gel).
25. Snap freeze the sample in liquid nitrogen.
26. Continue with reverse cross-linking for QC.

**3. Reversing the cross-link for QC – Day 2 & Day 3**

27. Put together:
    - 20 µl chromatin extract.
    - TE/SDS 1% to 95 µl.
28. Reverse cross-link by incubating overnight at 65°C in a thermoshaker (800 rpm).
29. The next morning (*Day 3*): Add 5 µl RNase A/T1 Mix (Thermo Scientific #EN0551) and incubate for 30 min at 37°C.
30. Add 40 µl of proteinase K (10 µg/µl) and incubate for 2 hours at 37°C.
31. Clean up with the Qiagen PCR purification kit (all steps performed at room temperature):
    - Add 5 volumes of PB buffer (700 µl for 140 µl sample).
    - Add sample to spin column.
    - Spin for 1 min at max speed (14000 rpm in Eppendorf 5417R).
    - Discard flow-through.
    - Wash 3 times with 500 µl of PE and spin down at max speed.
    - Transfer column to new 2.0 ml tube without lid and spin down at max speed for 1 min (to remove last bit of PE buffer).
    - Elute by adding 40 µl of EB, incubate for 1 min, and spin down at max speed in 1.5 ml Eppendorf tube.
32. Check DNA fragment size by loading 1 µl of purified DNA on Bioanalyzer with a High-Sensitivity DNA Chip (or equivalent).
    - The shearing is acceptable if the bioanalyzer peak is between 200-300bp. Depending on the application, longer fragment may also be acceptable. If there is still a large amount of longer fragments detected, the chromatin can be resheared for a few additional cycles. When this is done, fresh protease inhibitors should be added.

**4. Immunoprecipitation using magnetic beads – Day 3**

First bind the antibody to chromatin:

33. Thaw the chromatin on ice and take 20 µapart for input control
    - During this step make a mix of (per 6 samples): 210 µl Aprotinin, 7 µl Leupeptin and 7 µl Pepstatin.
34. Transfer 450 µl of the chromatin to a 1.5 ml Eppendorf tube.
35. Add 16 µl of the protease inhibitor mix and subsequently add 5 µl PMSF (in fumehood). Mix immediately by vortexing gently.
36. Add 1 µl of antibody to the chromatin extract
    - Depending on the antibody and the abundance of the target more antibody may be needed.
37. Incubate with rotation 2hrs @ 4 °C.

During the incubation of the chromatin with the antibody, the magnetic beads can be prepared:

38. Resuspend the magnetic beads (Dynabeads) in the vial (vortex > 30 sec).
39. Transfer 25 µl (0.75 mg) beads per ChIP to a 1.5 ml eppendorf tube.
40. Place the tube on the magnet (DynaMag-2) to separate the beads from the solution, and remove the supernatant.
41. Wash once with 500 µl PBS-T (PBS/0.02% Tween-20) by adding the PBS-T and doing a gentle vortex.
42. Place the tube on the magnet to separate the beads from the solution, and remove the supernatant.
43. Add 400 µl PBS+ 25 µl BSA (10mg/ml in TBST).
44. Incubate with rotation for ~2hrs @ 4 °C,
45. Place the tube on the magnet to separate the beads from the solution, and remove the supernatant (about 5-10 minutes before the incubation of the chromatin + antibody is ready).
46. Wash once with 500 µl PBS-T.
47. Place the tube on the magnet to separate the beads from the solution, and remove the supernatant.

Bind chromatin bound antibody to the beads:

48. Add 5 µl PMSF to the sample containing the antigen (CE 450ul + V5 antibody), add this to the beads and gently vortex to resuspend the beads. Incubate with rotation for 20 min at room temperature to allow the antibody to bind to the beads.
    - Depending on the antibody this incubation may need to be longer.
49. Do a quick spin and place the tube on the magnet. Transfer the supernatant to a clean tube for further analysis, if desired.
50. Wash the bead complex 2 times using 200 µl PBS for each wash. Separate on the magnet between each wash, remove supernatant and resuspend by gentle vortexing.
    - If the protein of interest is a particular strong binder, more stringent washes may help remove non-crosslinked proteins. Use wash buffer 1 and/or wash buffer 2 (2-3 washes each).
51. Wash the beads 1 time using 200 µl PBS-T. Resuspend by gentle vortexing, separate on the magnet after wash and remove supernatant.
52. Resuspend the beads in 100 µl PBS-T and transfer the bead suspension to a clean LoBind Eppendorf tube. This is recommended to avoid co-elution of proteins bound to the tube wall.
53. Proceed to Elution and reverse cross-linking.

**5. Elution and reverse cross-linking – Day 3 & Day 4**

54. Place the tube (from step 52 in "Immunoprecipitation using magnetic beads") on the magnet and remove the supernatant.
55. Add 98 µl TE/SDS.
56. Resuspend the beads-Ab-Ag complex by gentle vortexing.
57. Reverse the cross-links overnight using shaking (800 rpm) at 65$^{o}$C.
58. For input samples use 20 µl extract and 75 µl TE/SDS.
59. The next morning (*Day 4*): add 2 µl RNAse A/T1 mix (thermo #EN0551) for IP samples and 5 µl for Input samples and incubate 30 min @ 37 °C.
60. Add 40 µl of proteinase K (10 mg/ml in TE), and incubate for 2 hours at 37 °C.
61. Place the tube on the magnet and transfer the supernatant containing eluted DNA to a clean LoBind-tube.
62. Clean up with the Qiagen PCR purification up kit.
    - Add 5 volumes of PB buffer (700 µl for 140 µl sample).
    - Add sample to spin column.
    - Spin for 1 min at max speed.
    - Discard flow-through.
    - Wash 3 times with 500 µl of PE and spin down at max speed.
    - Transfer column to an empty 2.0 ml tube without lid and spin down at max speed for 1 min to remove last bit of PE buffer.
    - Elute by adding 40 µl of EB, incubate for >1 min, and spin down at max speed in new Lobind 1.5 ml tube.
        - The DNA can also be eluted in 30 µl of EB for more concentrated DNA.
63. Quantify the binding using qPCRs and/or proceed to make sequencing libraries.

**General notes for the protocol**

1. Making 4.5M Tris pH 8.0 is challenging, because this concentration is nearly at the solution limit of Tris. We recommend making 2 liters at the same time. For 100 ml cultures cross-linked with 2% formaldehyde, almost 55 ml of 4.5M Tris is used to quench. This means that for 18 samples nearly 1 liter of 4.5M Tris is needed. When dissolving Tris, a substantial amount of HCL must be added to get the Tris to dissolve at all, take this into account when adding the MQ. For 1L use 545.13 g Tris and ~211.48 ml of 12.1 M HCl. We recommend filter sterilizing the Tris.
2. During the protocol, a Qiagen PCR cleanup kit is used to clean up the IPs, inputs and QCs. The recovery of this kit is about 70%. We tried several other cleanups (Zymoclear ChIP kit, Ampure beads, phenol chloroform isolation and Qiagen MinElute DNA kit), but the Qiagen PCR cleanup was the best in both recovery and reproducibility.
3. Use an appropriate number of replicates. Since there are many steps in the protocol, the chances of variation arising anywhere in the protocol are substantial. We recommend using always at least three biological replicates, if possible.
4. Protein degradation during the procedure can have detrimental effects on the ChIP signal. It is therefore important that fresh protease inhibitors are added at the steps indicated in the protocol.

**Growth and cross-linking**

5. Always use fresh formaldehyde, ideally less than 1 month old (Rando, 2010). We have used formaldehyde solutions up to 3 months old. We recommend purchasing small bottles of formaldehyde and finishing them within a few (2-3) weeks after the first use, with minimal opening/closing of the bottle. Formaldehyde can polymerize when exposed to oxygen, which will lower the crosslinking efficiency.
6. To make sure that there is no artefactual binding of the protein of interest during the procedure, take along a non-cross-linking control. If there is no binding observed in this control, this indicates that there is no artificial binding of the protein of interest during the procedure. Treat this culture exactly the same as the cross-linked samples, but omit the formaldehyde addition step.
7. Be quick. Even though quenching with Tris is efficient, it is best to keep the time that samples spend on ice to a minimum. Ideally, only a few samples are ready at the same time, such that the samples can easily be processed together. If, for example, a time course experiment is performed with replicates, having 1 hour between the time courses gives enough time to finish harvesting the first, before the second has to be processed. We recommend harvesting the cultures with two people at the same time to speed up the process.

**Cell Lysis**

8. Although the bead beat protocol is more cumbersome, we prefer to use this over the zymolyase protocol, because of the extensive protein degradation during zymolyase protocol. However, the zymolyase protocol was not extensively tested with different proteins and perhaps other proteins are less susceptible to this degradation. A likely cause of the extensive degradation are the proteases present in zymolyase preparations. If lysis with zymolyase is preferred, it would be best to add protease inhibitors, and to wash extensively in Buffer Z/Sorbitol to remove the majority of proteases. However, care has to be taken that the spheroplasts are not lysed during the washing steps.
9. Step 19 in the bead beat protocol is an important step. During this step, the samples are centrifuged to pellet the unbroken cells and cell debris. The supernatant contains the chromatin. With longer cross-linking times, there is an increased chance of chromatin co-precipitating with the pellet. If this happens, it should be visible as a vague white layer on top of the pellet. Make sure to transfer this chromatin pellet together with the supernatant, otherwise the majority of the (cross-linked) DNA will be lost!

- What will count as "long cross-linking times" depends on the concentration of formaldehyde used. With 1% formaldehyde we noticed the co-precipitation of the chromatin when cross-linking for 20 or 30 minutes, but not with cross-linking for 10 minutes.

10. During the full bead beat protocol, there is an additional centrifugation step before the sonication, to separate the chromatin (CE) from the rest of the cell lysate. We omitted this step in the short bead beat protocol, because in our experience it was often very difficult to resuspend the pellet. In addition, the chromatin is harder to pellet with short (5 min) cross-linking times, which may lead to loss of some of the chromatin during this step. However, when the CE and the WCL are not separated, this means that unbound TF is mixed with the chromatin during the IP, which could potentially lead to *in vitro* binding of the DNA.
11. The sonication is a tricky part of the protocol, which has to be optimized for each device. In this protocol a bioruptor pico is used. Sonication can break down the protein of interest, but this mainly happens when proteins are really big (> 80-100 kDa). This protocol uses 15 seconds on / 30 seconds off to reduce the extent of breakdown.
    - To assess the extent of degradation during this step, samples taken before and after sonication can be monitored using Western blotting.
    - If the degradation is severe, enzymatic fragmentation using for example MNase may be used instead of sonication.

**QC**

12. For all samples a quality check should be done to assess the extent of the fragmentation. It is best to run all QC samples on a bioanalyzer High Sensitivity DNA ChIP (or equivalent) to check if the fragmentation was sufficient and reproducible. It is important that the average size is comparable between samples, because it will have an effect on the quantitation.

**IP**

13. After the incubation with the chromatin and antibody, the magnetic beads are washed 2x with PBS and 2x with PBS-T. This was sufficient for most proteins tested. However, if the protein of interest is a strong binder, more stringent washes may be needed. Examples of more stringent wash buffers are wash buffer 1 (FA lysis with 500 mM NaCl) or wash buffer 2 (10 mM Tris pH 8.0, 0.25 mM LiCl, 1 mM EDTA pH 8.0, 0.5% Nonidet P-40 and 0.5% Na-deoxycholate). Taking along a non-cross-linking control should show whether the more stringent washes are sufficient or not and whether a protein can bind to DNA without crosslinking.

**qPCR**

14. When doing qPCR, always take a (5x 10-fold dilution) standard curve along for each primer pair on each plate. It is best if this standard is created from fragmented, cleaned, genomic DNA and that the same material is used for all standard curves. This way, different plates can be compared with each other.
15. Always include a background control with each qPCR: a promoter that does not have binding of the TF of interest.
    - When a background control is included, the ratio over background can be calculated. However, if the background is very low, which is usually the case when using magnetic beads, the background can be hard to quantify using qPCR. If there is a lot of variation in the background signal, the enrichment over background will also have a lot of variation.
