## Supplementary material for "An extensively optimized chromatin immunoprecipitation protocol for quantitatively comparable and robust results": Methods per figure plus supplemental tables

### Figure 2

*Figure 2A*

Cbf1-aa Stb4-V5 cells used for Figure 2A-2C were prepared in a big batch, by growing several 250 ml cultures. Cultures were cross-linked with 1% formaldehyde by adding 6.8 ml of 37% formaldehyde and incubating them for 5 minutes. Subsequently, 12.8 ml 2.5 M glycine was added to a final concentration of 125 mM and incubated for 5 minutes. Cells were harvested as described in the general methods section.

Six cell pellets of Cbf1-aa strains were lysed using the zymolyase protocol, by treating them with zymolyase solution (10 mg/ml in buffer Z) for 10 minutes. After two careful washes with buffer Z, each cell pellet was resuspended in 540 μl FA lysis buffer. All samples were pooled, mixed and split over 12x 1.5 ml sonication tubes, by putting 300 μl in each tube. Half of the samples were sonicated 30 seconds on / 30 seconds off for 3 cycles and the other half for 4 cycles in a bioruptor pico. All samples that were sheared for the same number of cycles were pooled and 250 μl of chromatin was used for each IP.

Eight samples (4 that were sonicated for 3 cycles and 4 that were sonicated for 4 cycles) were used for the IP with magnetic beads while 4 samples (2 that were sonicated for 3 cycles and 2 that were sonicated for 4 cycles) were used for the IP using agarose beads. The IPs with magnetic and agarose beads were performed as described in the general methods section with some modifications. 50 μl beads and 2 μl anti-V5 antibody were used for the IPs with magnetic beads, while 20 μl of anti-V5-agarose beads were used. The agarose beads were prepared by washing 3 times in FA lysis buffer prior to use, while the magnetic beads were washed with 500 μl PBS-T. After incubation with the chromatin, the agarose beads were washed using FA lysis buffer, wash buffer 1 and wash buffer 2 as described in the general methods section. Several replicates of the magnetic IPs were performed with different washes. Two IPs with material that was sonicated for 3 cycles were washed with the standard washes of 2x PBS and 2x PBS-T. Two IPs with material that was sonicated for 4 cycles were washed by doing 3 washes using PBS. The other four IPs (two that were sonicated for 3 cycles and two that were sonicated for 4 cycles) were washed first 2x with PBS and then 2x using wash buffer 2. The DNA was recovered by eluting overnight in TE/SDS at 65°C. Although the washes were different between the samples, the average of all the agarose beads IPs (four replicates) and the average of all the magnetic beads IPs (eight replicates) are shown in Figure 2A.

*Figure 2B and 2C*

The results shown in Figure 2B and 2C were performed in the same experiment on the same day. The results were split over 2 figures for clarity. The same samples are shown in Figure 2B: TE/SDS and Figure 2C: 20 min beads inc.

Cbf1-aa Stb4-V5 cells used for Figure 2A-2C were prepared in a big batch, by growing several 250 ml cultures. The cultures were cross-linked with 1% formaldehyde by adding 6.8 ml of 37% formaldehyde and incubating them for 5 minutes. Subsequently, 12.8 ml 2.5 M glycine was added to a final concentration of 125 mM and incubated for 5 minutes. The cells were harvested as described in the general methods section.

Five pellets were lysed using the zymolyase protocol, by incubating them in zymolyase solution (10 mg/ml in buffer Z) for 10 minutes. After two careful washes with buffer Z, each cell pellet was resuspended in 550 μl FA lysis buffer. All samples were pooled, mixed and split over 10x 1.5 ml sonication tubes, by putting 300 µl in each tube. The samples were sonicated for 3 cycles, 30 seconds on / 30 seconds off. All samples were pooled and 250 µl of chromatin was used for each IP.

The IPs with magnetic beads were performed as described in the general methods section, with some modifications. 50 µl magnetic beads and 2 µl anti-V5 antibody were used for the IPs. Prior to the incubation, the beads were washed with 500 µl PBS-T. For the IPs shown in Figure 2B and 2C, the antibody was incubated with the chromatin overnight at 4$^{o}$C, except for the samples "pre inc Ab + beads", where the beads were incubated with the antibody overnight at 4$^{o}$C in 200 µl PBS-T. For Figure 2B, the chromatin + antibody was bound to the beads by incubating them with the beads at RT for 20 minutes. For Figure 2C, this incubation was done for either 20 minutes or 60 minutes. In addition, the antibody conjugated beads were incubated with the chromatin at RT for 20 minutes as well. The beads were washed with 2x PBS and 2x PBS-T. The DNA was eluted either by incubating the beads overnight at 65$^{o}$C in 98 µl TE/SDS or by incubating the beads in 20 µl 50 mM glycine pH 2.8 for 2 minutes. The glycine eluted beads were separated from the supernatant and the beads were resuspended in 98 µl TE/SDS and a second elution was done overnight at 65$^{o}$C. The 20 µl supernatant was mixed with 75 µl TE/SDS and incubated overnight at 65$^{o}$C as well. All IPs were performed in duplicate.

*Figure 2D*
Cbf1-aa cells used in Figure 2D were prepared in a big batch, by growing several 250 ml cultures. The cultures were cross-linked with 1% formaldehyde by adding 6.8 ml of 37% formaldehyde to the cultures and incubating them for 5 minutes. Subsequently, 12.8 ml 2.5 M glycine was added to a final concentration of 125 mM and incubated for 5 minutes. The cells were harvested as described in the general methods section.

Six Cbf1-aa pellets were lysed using the zymolyase protocol, by incubating them in zymolyase solution (10 mg/ml in buffer Z) for 10 minutes. After two careful washes with buffer Z, each cell pellet was resuspended in 550 µl FA lysis buffer. All samples were pooled, mixed and split over 12x 1.5 ml sonication tubes, by putting 300 µl in each tube. The samples were sonicated for 3 cycles, 30 seconds on / 30 seconds off. 3 tubes were pooled and 200 µl was used for the IPs of the overnight samples. The next day, the rest of the tubes were pooled and 200 µl of this pool was used for each IP where the chromatin and antibody were incubated for 1-4 hours.

The IPs with magnetic beads were performed as described in the general methods section with some modifications. 25 µl magnetic beads and 1 µl anti-V5 antibody were used for the IPs. The chromatin was incubated with the anti-V5 antibody for 1 hour, 2 hours, 4 hours or overnight. The incubations were staggered such that all IPs were ready at the same time, and all washes were performed in parallel. The standard wash of 2x PBS and 2x PBS-T was used for all samples. The DNA was recovered by eluting overnight in TE/SDS at 65$^{o}$C. The IPs were performed in triplicate.

*Figure 2E*
Cbf1-aa cells used in Figure 2E were grown as 100 ml cultures. The cells were cross-linked by adding 2.7 ml of 37% formaldehyde to a final concentration of 1% and incubating them for 5 minutes. Subsequently, 5.1 ml of 2.5M glycine was added to a final concentration of 125 mM and the cells were incubated for 5 minutes. The cells were harvested as described in the general methods section.

The Cbf1-aa pellets were lysed using the zymolyase protocol, by incubating them in zymolyase solution (10 mg/ml in buffer Z) for 10 minutes. After two careful washes with buffer Z, each cell pellet was resuspended in 550 µl FA lysis buffer. All samples were pooled, mixed and split over 12x 1.5 ml sonication tubes, by putting 300 µl in each tube. The samples were sonicated for 3 cycles, 30 seconds on / 30 seconds off and the sheared chromatin was pooled per cell pellet and 200 μl of chromatin was used for each IP.

The IPs with magnetic beads were performed as described in the general methods section with some modifications. 25 μl magnetic beads and 1 μl anti-V5 antibody were used for the IPs. The chromatin was incubated with the anti-V5 antibody for 2 hours. During the incubation of the chromatin with the antibody, the beads were prepared. 25 μl beads per IP was washed in individual 1.5 ml Eppendorf tubes using 500 μl PBS-T. For the samples without BSA, right before the addition of the chromatin and antibody, the beads were washed once in 500 μl PBS-T. For the sample with BSA, the beads were resuspended in 200 μl PBS, 12.5 μl BSA (10 mg/ml in TBS-T) was added and the beads were incubated at 4$^{o}$C while the chromatin was incubating with the antibody. Just before this incubation was finished, the beads were washed again with 500 μl PBS-T. The standard wash of 2x PBS and 2x PBS-T was used for all samples. The DNA was recovered by eluting overnight in TE/SDS at 65$^{o}$C. The IPs were performed in triplicate.

**Figure 3**

*Figure 3A and 3B*
The results shown in Figure 3A and 3B were performed in the same experiment on the same day. The results were split over 2 figures for clarity. The same samples are shown for 5 min 125 mM glycine in both Figure 3A and 3B.

Cbf1-aa Cha4-V5 cells used for Figure 3A and 3B were grown as 100 ml cultures. The cells were cross-linked by adding 2.7 ml of 37% formaldehyde to a final concentration of 1% and incubating them for 1 minute. Subsequently, 5.1 ml of 2.5M glycine was added to a final concentration of 125 mM and the cells were incubated for 1, 5 or 10 minutes. In addition, some samples were incubated with 10.7 ml 2.5M glycine, at final concentration of 250 mM instead of 125 mM, and incubated for 5 minutes. The cells were harvested as described in the general methods section.

The Cbf1-aa Cha4-V5 pellets were lysed using the zymolyase protocol, by incubating them in zymolyase solution (10 mg/ml in buffer Z) for 10 minutes. After two careful washes with buffer Z, each cell pellet was resuspended in 550 μl FA lysis buffer. The samples were split over 2x 1.5 ml sonication tubes, by putting 300 μl in each tube. All samples were sonicated for 3 cycles, 30 seconds on / 30 seconds off and the sheared chromatin was pooled per cell pellet. 200 μl of chromatin was used for each IP.

The IPs with magnetic beads were performed as described in the general methods section. 25 μl magnetic beads and 1 μl anti-V5 antibody were used for the IPs. The chromatin was incubated with the anti-V5 antibody for 2 hours. During the incubation of the chromatin with the antibody, the beads were prepared. 25 μl beads per IP was washed in individual 1.5 ml Eppendorf tubes using 500 μl PBS-T. The beads were resuspended in 200 μl PBS, 12.5 μl BSA (10 mg/ml in TBS-T) was added and the beads were incubated at 4$^{o}$C while the chromatin was incubating with the antibody. Just before this incubation was finished, the beads were washed again with 500 μl PBS-T. The standard wash of 2x PBS and 2x PBS-T was used for all samples. The DNA was recovered by eluting overnight in TE/SDS at 65$^{o}$C. The IPs were performed in triplicate.

*Figure 3C and 3D*
Cbf1-aa cells used for Figure 3A and 3B were grown as 100 ml cultures. The cells were cross-linked by adding 2.7 ml of 37% formaldehyde to a final concentration of 1% and incubating them for 1 minute (1 min non-quenched, glycine quenched and Tris quenched samples) or 6 minutes (6 min non-quenched sample). Samples that were not quenched were immediately transferred to 50 ml tubes and

centrifuged as described in the general methods sections. To the glycine quenched samples, 5.1 ml of 2.5M glycine was added to a final concentration of 125 mM and the cells were incubated for 5 minutes. The Tris quenched samples were incubated with 20.5 ml 4.5M Tris pH 8.0 at a final concentration of 750 mM for 1, 5 or 10 minutes. All samples were harvested as described in the general methods section.

The Cbf1-aa pellets were lysed using the zymolyase protocol, by incubating them in zymolyase solution (10 mg/ml in buffer Z) for 10 minutes. After two careful washes with buffer Z, each cell pellet was resuspended in 550 µl FA lysis buffer. All samples were split over 2x 1.5 ml sonication tubes, by putting 300 µl in each tube. The samples were sonicated for 3 cycles, 30 seconds on / 30 seconds off and the sheared chromatin was pooled per cell. 200 µl of chromatin was used for each IP.

The IPs with magnetic beads were performed as described in the general methods section. 25 µl magnetic beads and 1 µl anti-V5 antibody were used for the IPs. The chromatin was incubated with the anti-V5 antibody for 2 hours. During the incubation of the chromatin with the antibody, the beads were prepared. 25 µl beads per IP was washed in individual 1.5 ml Eppendorf tubes using 500 µl PBS-T. The beads were resuspended in 200 µl PBS, 12.5 µl BSA (10 mg/ml in TBS-T) was added and the beads were incubated at 4°C while the chromatin was incubating with the antibody. Just before this incubation was finished, the beads were washed again with 500 µl PBS-T. The standard wash of 2x PBS and 2x PBS-T was used for all samples. The DNA was recovered by eluting overnight in TE/SDS at 65°C. The IPs were performed in triplicate.

**Figure 4**

*Figure 4A*
Cbf1-aa cells used in Figure 4A were grown in the same big batch as those used for Figure 2D, by growing several 250 ml cultures. The cultures were cross-linked with 1% formaldehyde by adding 6.8 ml of 37% formaldehyde and incubating them for 5 minutes. Subsequently, 12.8 ml 2.5 M glycine was added to a final concentration of 125 mM and incubated for 5 minutes. The cells were harvested as described in the general methods section.

The Cbf1-aa pellets were lysed using the zymolyase protocol, by incubating them in zymolyase solution (10 mg/ml in buffer Z) for 10 or 25 minutes. After two careful washes with buffer Z, each cell pellet was resuspended in 550 µl FA lysis buffer. The samples were split over 2x 1.5 ml sonication tubes, by putting 300 µl in each tube. All samples were sonicated for 3 cycles, 30 seconds on / 30 seconds off and the sheared chromatin was pooled per cell pellet. 200 µl of chromatin was used for each IP.

The IPs with magnetic beads were performed as described in the general methods section. 25 µl magnetic beads and 1 µl anti-V5 antibody were used for the IPs. The chromatin was incubated with the anti-V5 antibody for 2 hours. The beads were washed once in 500 µl PBS-T, the chromatin and antibody was added to the beads and incubated at RT for 20 minutes. The beads were washed with the standard wash of 2x PBS and 2x PBS-T and DNA was recovered by eluting overnight in TE/SDS at 65°C. The IPs were performed in triplicate.

*Figure 4B and 4C*
Cbf1-aa cells used in Figure 4B and 4C were grown in the same big batch as those used for Figure 2D and 4A, by growing several 250 ml cultures. The cultures were cross-linked with 1% formaldehyde by adding 6.8 ml of 37% formaldehyde and incubating them for 5 minutes. Subsequently, 12.8 ml 2.5 M glycine was added to a final concentration of 125 mM and incubated for 5 minutes. The cells were harvested as described in the general methods section.

Four Cbf1-aa pellets, which are equivalent to 100 ml cell cultures OD 0.8, plus some left over Cbf1-aa material was resuspended in 720 μl buffer Z and divided over 6 tubes. A 20 μl aliquot was taken as a pre-lysis control for Western blotting. The cells were lysed using the zymolyase protocol, by incubating them in zymolyase solution (10 mg/ml in buffer Z) for 10 minutes. Two samples were incubated without protease inhibitors and 4 samples were incubated with protease inhibitors (one tablet of EDTA-free cOmplete protease inhibitor cocktail (Roche #11873580001) per 10 ml of zymolyase solution). The samples that were lysed without protease inhibitors and two of the samples that were lysed with protease inhibitors were washed twice with buffer Z without protease inhibitors. The other two samples that were lysed in the presence of protease inhibitors were washed with buffer Z containing one tablet cOmplete protease inhibitor cocktail (Roche #11836145001) per 10 ml buffer Z. After the washes, each cell pellet was resuspended in 550 μl FA lysis buffer and a 20 μl aliquot was taken for Western blotting. The samples were split over 2x 1.5 ml sonication tubes, by putting 300 μl in each tube, and were sonicated for 3 cycles, 30 seconds on / 30 seconds off. The sheared chromatin was pooled per cell pellet and for each sample a 20 μl aliquot was taken for Western blotting both before and after the centrifugation step. Because there were multiple aliquots taken for Western blotting, only 150 μl of chromatin was left to use in each IP.

The IPs with magnetic beads were performed as described in the general methods section. 25 μl magnetic beads and 1 μl anti-V5 antibody were used for the IPs. The chromatin was incubated with the anti-V5 antibody for 2 hours. 25 μl beads per IP was washed in individual 1.5 ml Eppendorf tubes using 500 μl PBS-T. The beads were resuspended in 200 μl PBS, 12.5 μl BSA (10 mg/ml in TBS-T) was added and the beads were incubated at 4°C while the chromatin was incubating with the antibody. Just before this incubation was finished, the beads were washed again with 500 μl PBS-T. The chromatin and antibody were added to the beads, and incubated at RT for 20 minutes. The beads were washed with the standard wash of 2x PBS and 2x PBS-T and DNA was recovered by eluting overnight in TE/SDS at 65°C. The IPs were performed in duplicate. The results shown in Figure 4B are the average of two IPs (without inhibitors) or of four IPs (with inhibitors). The average signal of the samples with inhibitors during the zymolyase treatment is shown, regardless of whether or not they were washed with buffer Z containing protease inhibitors.

Samples for Western blotting were prepared by mixing 20 μl of the sample with 20 μl of 5X sample buffer (5% SDS, 200 mM Tris pH 6.8, 25% glycerol, 1.43 M β-mercaptoethanol, 0.032% bromophenol blue). The pre-lysis samples were prepared as described in the general methods section, by lysing the cells using NaOH. All but the control samples were heated to 95°C for 30 minutes to reverse cross-links. The samples (10 μl) were separated on a 10% polyacrylamide gel. As a control, a non-cross-linked crude lysate of a WT-aa and a Cbf1-aa strain were taken along, which were heated to 95°C for 5 minutes prior to loading. A chromatin extract of a Cbf1-aa Cha4-V5 strain prepared with the full bead beating protocol was also included. After the proteins had migrated through the gel, the proteins were transferred to a nitrocellulose membrane. The membrane was stained with an anti-V5 antibody (Life Technologies #R96025) and binding was visualized using a goat-anti-mouse-HRP conjugated antibody (Bio-Rad #1706516) and ECL solution, as described in the general methods section.

**Figure 5**

*Figure 5A*

Cbf1-aa Cha4-V5 cells used in Figure 5A were grown as 100 ml cultures. The cells were cross-linked by adding 2.7 ml of 37% formaldehyde to a final concentration of 1% and incubating them for 5 minutes. Subsequently, 20.5 ml of 4.5M Tris pH 8.0 was added to a final concentration of 750 mM and the cells were incubated for 1 minute. The cells were harvested as described in the general methods section

Cells were lysed either using zymolyase or the full bead beating protocol. When using zymolyase to lyse the cells, the Cbf1-aa Cha4-V5 cell pellets were incubated in zymolyase solution (10 mg/ml in buffer Z) for 10 minutes. After two careful washes with buffer Z, each cell pellet was resuspended in 550 μl FA lysis buffer. The samples were split over 2x 1.5 ml sonication tubes, by putting 300 μl in each tube. All samples were sonicated for 3 cycles, 30 seconds on / 30 seconds off. Sheared chromatin was pooled per cell pellet. The samples that did not have appropriate fragment lengths were sheared for an additional cycle. 200 μl of chromatin was used for each IP. The cells that were lysed with the bead beating protocol were bead beated for 7x 3 minutes in a genie disruptor. The cell debris was pelleted and the lysate centrifuged at 18407g for 15 minutes to pellet the chromatin. The chromatin was then washed with FA lysis buffer and resuspended in 600 μl FA lysis buffer. The chromatin was subsequently fragmented by splitting each sample over 2x 1.5 ml sonication tubes and sonicating 30 seconds on / 30 seconds off. The sheared chromatin was pooled per cell pellet. 200 μl chromatin was used for each IP.

IPs with magnetic beads were performed as described in the general methods section. 25 μl magnetic beads and 1 μl anti-V5 antibody were used for the IPs. The chromatin was incubated with the anti-V5 antibody for 2 hours. 25 μl beads per IP was washed in individual 1.5 ml Eppendorf tubes using 500 μl PBS-T. The beads were resuspended in 200 μl PBS, 12.5 μl BSA (10 mg/ml in TBS-T) was added and the beads were incubated at 4$^{o}$C while the chromatin was incubating with the antibody. Just before this incubation was finished, the beads were washed again with 500 μl PBS-T. The chromatin and antibody were added to the beads and incubated at RT for 20 minutes. The beads were washed with the standard wash of 2x PBS and 2x PBS-T and DNA was recovered by eluting overnight in TE/SDS at 65$^{o}$C. The IPs were performed in triplicate.

*Figure 5B-5D*
Cbf1-aa Cha4-V5 cells used in Figure 5B-5D were grown in a big batch, by growing several 250 ml cultures. The cultures were cross-linked with 1% formaldehyde by adding 7.0 ml of 37% formaldehyde and incubating them for 5 minutes. Subsequently, 32.1 ml 4.5 M Tris was added to a final concentration of 500 mM and incubated for 1 minute. The cells were harvested as described in the general methods section.

The Cbf1-aa Cha4-V5 cells were lysed either using the zymolyase protocol, the full bead beating protocol or the short bead beating protocol. 4 Cbf1-aa Cha4-V5 cell pellets were resuspended in 1 ml of Buffer Z and a 20 μl aliquot was taken as a pre-lysis control for Western blotting. The cells were then lysed using the zymolyase protocol, by incubating them in zymolyase solution (10 mg/ml zymolyase in buffer Z containing one tablet of EDTA-free cOmplete protease inhibitor cocktail (Roche #11873580001) per 10 ml) for 10 minutes. The samples were subsequently washed with buffer Z containing one tablet Roche cOmplete protease inhibitor cocktail (Roche #11873580001) per 10 ml buffer Z. After the washes, each cell pellet was resuspended in 550 μl FA lysis buffer and a 20 μl aliquot was taken for Western blotting (pre-sonication). The samples were split over 2x 1.5 ml sonication tubes, by putting 300 μl in each tube and were sonicated for 3 cycles, 30 seconds on / 30 seconds off. All sheared chromatin was pooled per cell pellet and for each sample a 20 μl aliquot was taken for Western blotting.

The other samples were lysed using either the full or the short bead beating protocol. 8 Cbf1-aa Cha4-V5 pellets were resuspended in 1 ml of FA lysis buffer and a 20 μl aliquot was taken for Western blotting. The tubes were filled with FA lysis buffer and the cells were subsequently bead beated for 7x 3 minutes in a genie disruptor. The cell debris was pelleted and for the 4 samples that were processed with the full bead beat protocol, the lysate was centrifuged at 18407g for 15 minutes to pellet the chromatin, which contains the CE. The rest of the lysate (WCL) was kept on ice. The CE was then washed once with FA lysis buffer and resuspended in 600 μl FA lysis buffer. A 20 μl sample was taken for Western blotting. The CE was subsequently fragmented by splitting each sample over 2x 1.5 ml sonication tubes and sonicating for 3 cycles, 30 seconds on / 30 seconds off. The sheared chromatin was pooled per cell pellet. Before sonicating the WCL and the short bead beat chromatin, a 20 μl sample was taken for Western Blotting, and then the samples were sheared in 15 ml sonication tubes containing 300 μl sonication beads (Diagenode), also for 3 cycles 30 seconds on / 30 seconds off. A 20 μl sample was taken for Western blotting from 2 samples of all 3 protocols after the sonication. 200 μl of the CE was used for IPs, while 450 μl of the WCL and the short bead beat chromatin was used.

The IPs with magnetic beads were performed as described in the general methods section. 25 μl magnetic beads and 1 μl anti-V5 antibody were used for the IPs. The chromatin was incubated with the anti-V5 antibody for 2 hours. 25 μl beads per IP was washed in individual 1.5 ml Eppendorf tubes using 500 μl PBS-T. The beads were resuspended in 200 μl PBS, 12.5 μl BSA (10 mg/ml in TBS-T) was added and the beads were incubated at 4°C while the chromatin was incubating with the antibody. Just before this incubation was finished, the beads were washed again with 500 μl PBS-T. The chromatin and antibody were added to the beads and incubated at RT for 20 minutes. The beads were washed with the standard wash of 2x PBS and 2x PBS-T and DNA was recovered by eluting overnight in TE/SDS at 65°C. The IPs were performed either in duplicate (WCL) or quadruplicate (CE and short bead beat protocol).

To calculate the relative amount of input DNA, the starting quantities (SQs) of the input samples from Figure 5B were taken and the average SQ of the inputs of the CE was set to 100%. The values of all input samples were scaled accordingly.

The samples for Western blotting were prepared by mixing 20 μl of the sample with 20 μl of 5X sample buffer (5% SDS, 200 mM Tris pH 6.8, 25% glycerol, 1.43 M β-mercaptoethanol, 0.032% bromophenol blue). The pre-lysis samples were prepared as described in the general methods section, by lysing the cells using NaOH. All samples were heated to 95°C for 30 minutes to reverse cross-links. The samples (10 μl) were separated on a 10% polyacrylamide gel. As a control, a non-cross-linked crude lysate of a Cbf1-aa strain, heated to 95°C for 5 minutes prior to loading, was taken along on both gels. After the proteins had migrated through the gel, the proteins were transferred to a nitrocellulose membrane. The membrane was stained with an anti-V5 antibody (Life Technologies #R96025) and binding was visualized using a goat-anti-mouse-HRP conjugated antibody (Bio-Rad #1706516) and ECL solution, as described in the general methods section.

**Figure 6**

*Figure 6A*

Cbf1-aa Cha4-V5 cells used in Figure 6A were grown as 100 ml cultures. At OD = 0.52, 1 hour before the cells reached OD = 0.8, 367.4 μl 2 mM rapamycin (60 min depleted samples, final concentration 7.5 μM) or 367.4 μl or DMSO (non-depleted samples) was added. The cells that were cross-linked (depleted and non-depleted) were incubated with 2.7 ml of 37% formaldehyde to a final concentration of 1% for 5 minutes. Subsequently, 20.5 ml of 4.5M Tris pH 8.0 was added to a final concentration of 750 mM and the cells were incubated for 1 minute. The non-cross-linked samples were either incubated for 6 minutes with 20.5 ml of 4.5M Tris pH 8.0 at a final concentration of 750 mM or first incubated with 20.5 ml of 4.5M Tris pH 8.0 for 1 minute and then 2.7 ml of 37% formaldehyde was added to a final concentration of 1% for 5 minutes. The cells were harvested as described in the general methods section

The Cbf1-aa cells were bead beated for 7x 3 minutes in a genie disruptor. The cell debris was pelleted and the supernatant was used for sonication. The chromatin was fragmented by splitting each sample over 2x 1.5 ml sonication tubes and sonicating 30 seconds on / 30 seconds off. The sheared chromatin was pooled per cell pellet. 450 µl of chromatin was used for each IP.

The IPs with magnetic beads were performed as described in the general methods section. 25 µl magnetic beads and 1 µl anti-V5 antibody were used for the IPs. The chromatin was incubated with the anti-V5 antibody for 2 hours. 25 µl beads per IP was washed in individual 1.5 ml Eppendorf tubes using 500 µl PBS-T. The beads were resuspended in 200 µl PBS, 12.5 µl BSA (10 mg/ml in TBS-T) was added and the beads were incubated at 4$^{o}$C while the chromatin was incubating with the antibody. Just before this incubation was finished, the beads were washed again with 500 µl PBS-T. The chromatin and antibody were added to the beads and incubated at RT for 20 minutes. The beads were washed with the standard wash of 2x PBS and 2x PBS-T and DNA was recovered by eluting overnight in TE/SDS at 65$^{o}$C. The IPs were performed in triplicate, and the average of the samples where only Tris was added and the samples where Tris was added before the formaldehyde was used as the signal of the non-cross-linked control (6 replicates in total).

*Figure 6B*
Cbf1-aa cells were grown in 100 ml cultures. 60 minutes (at OD = 0.52) and 15 minutes (at OD = 0.72) before the cells reached OD = 0.8, 367.4 µl 2 mM rapamycin was added to the different cultures (final concentration 7.5 µM). As a non-depleted control, a culture was taken along where 367.4 µl DMSO was added at OD = 0.52. When the cultures were ready, a 1 ml aliquot was taken which was fixed using 100% methanol as is described in the general methods section. 1.5 µl was mixed with 1.5 µl 1% agarose on a slide to image the cells.

*Figure 6C and 6D*
Abf1-aa (Figure 6C) and Reb1-aa (Figure 6D) cells were grown as 100 ml cultures. The cells were cross-linked by adding 2.7 ml of 37% formaldehyde to a final concentration of 1% and incubating them for 5 minutes. Subsequently, 20.5 ml of 4.5M Tris pH 8.0 was added to a final concentration of 750 mM and the cells were incubated for 1 minute. There was no formaldehyde added to the non-cross-linked samples, but these samples were still incubated with Tris. The cells were harvested as described in the general methods section.

The Abf1-aa and Reb1-aa cells were bead beated for 7x 3 minutes in a genie disruptor. The cell debris was pelleted and the supernatant was used for sonication. The chromatin was fragmented by splitting each sample over 2x 1.5 ml sonication tubes and sonicating for 4 cycles 30 seconds on / 30 seconds off. The sheared chromatin was pooled per cell pellet. 450 µl of chromatin was used for each IP.

The IPs with magnetic beads were performed as described in the general methods section. 25 µl magnetic beads and 1 µl anti-V5 antibody were used for the IPs. The chromatin was incubated with the anti-V5 antibody for 2 hours. 25 µl beads per IP was washed in individual 1.5 ml Eppendorf tubes using 500 µl PBS-T. The beads were resuspended in 400 µl PBS, 25 µl BSA (10 mg/ml in TBS-T) was added and the beads were incubated at 4$^{o}$C while the chromatin was incubating with the antibody. Just before this incubation was finished, the beads were washed again with 500 µl PBS-T. The chromatin and antibody were added to the beads and incubated at RT for 20 minutes. The Abf1-aa samples were washed with the standard wash of 2x PBS and 2x PBS-T, while the Reb1-aa were washed twice with wash buffer 1 (FA lysis buffer containing 500 mM NaCl) before the standard washes. DNA was recovered by eluting overnight in TE/SDS at 65$^{o}$C. The IPs were performed in triplicate

*Figure 6E and 6F*
The Abf1-aa (Figure 6E) and Reb1-aa (Figure 6F) cells were grown first as 200 ml cultures. 750 µl 2 mM rapamycin was added to a final concentration of 7.5 µM at OD=0.56 to deplete the proteins from

the nucleus. To the non-depleted samples 750 µl DMSO was added. 100 ml per culture was cross-linked by adding 2.7 ml of 37% formaldehyde to a final concentration of 1% and incubating them for 5 minutes. Subsequently, 20.5 ml of 4.5M Tris pH 8.0 was added to a final concentration of 750 mM and the cells were incubated for 1 minute. The cells were harvested as described in the general methods section.

The Abf1-aa and Reb1-aa cells were bead beated for 7x 3 minutes in a genie disruptor. The cell debris was pelleted and the supernatant was used for sonication. The chromatin was fragmented by splitting each sample over 2x 1.5 ml sonication tubes and sonicating for 4 cycles 30 seconds on / 30 seconds off. The sheared chromatin was pooled per cell pellet. 450 µl of chromatin was used for each IP.

The IPs with magnetic beads were performed as described in the general methods section. 25 µl magnetic beads and 1 µl anti-V5 antibody were used for the IPs. The chromatin was incubated with the anti-V5 antibody for 2 hours. 25 µl beads per IP was washed in individual 1.5 ml Eppendorf tubes using 500 µl PBS-T. The beads were resuspended in 400 µl PBS, 25 µl BSA (10 mg/ml in TBS-T) was added and the beads were incubated at 4$^{o}$C while the chromatin was incubating with the antibody. Just before this incubation was finished, the beads were washed again with 500 µl PBS-T. The chromatin and antibody were added to the beads and incubated at RT for 20 minutes. The beads were washed with the standard wash of 2x PBS and 2x PBS-T and DNA was recovered by eluting overnight in TE/SDS at 65$^{o}$C. The IPs were performed in triplicate

**Figure 7**

The results shown in Figure 7B, 7C and 7D were performed in the same experiments on the same days. The results were split over 2 figures for clarity. The same samples are shown in Figure 7B: Standard wash and Figure 7C and 7D: RT 20 min.

Sum1-aa cells used in Figure 7 were prepared in a big batch, by growing several 250 ml cultures. The cultures were cross-linked with 1% formaldehyde by adding 7.0 ml of 37% formaldehyde and incubating them for 5 minutes. Subsequently, 32.1 ml 4.5 M Tris was added to a final concentration of 500 mM and incubated for 1 minute. The cells were harvested as described in the general methods section.

The Sum1-aa cells were bead beated for 7x 3 minutes in a genie disruptor. The cell debris was pelleted and the supernatant was used for sonication. The chromatin was fragmented by splitting each sample over 2x 1.5 ml sonication tubes and sonicating for 4 cycles 30 seconds on / 30 seconds off. The sheared chromatin was pooled per cell pellet. 450 µl of chromatin was used for each IP.

On each day that IPs were performed, several chromatin extracts were pooled and split again to use for the IPs (the results shown in Figure 7A were obtained on a different day than the results shown in Figure 7B-7D). 25 µl (Figure 7A) or 50 µl (Figure 7B-7D) magnetic beads and 1 µl anti-V5 antibody were used for the IPs. The chromatin was incubated with the anti-V5 antibody for 2 hours. 25 µl beads per IP was washed in individual 1.5 ml Eppendorf tubes using 500 µl PBS-T. The beads were resuspended in 400 µl PBS, 25 µl BSA (10 mg/ml in TBS-T) was added and the beads were incubated at 4$^{o}$C while the chromatin was incubating with the antibody. Just before this incubation was finished, the beads were washed again with 500 µl PBS-T. The chromatin and antibody were added to the beads and incubated at RT for 20 minutes or at 4$^{o}$C for 20 or 60 minutes (Figure 7D and 7D). The beads were washed with the standard wash of 2x PBS and 2x PBS-T, 1x with PBS and 1x with PBS-T (Figure 7B low wash), with 2x FA lysis buffer, 2x wash buffer 1 (FA lysis buffer with 500 mM NaCl), 2x wash buffer 2 (10 mM Tris pH 8.0, 0.25 mM LiCl, 1 mM EDTA pH 8.0, 0.5% Nonidet P-40 and 0.5% Na-deoxycholate) and 1x PBS-T or with 2x PBS, 2x wash buffer 1 and 3x PBS-T . DNA was recovered by eluting overnight in TE/SDS at 65°C. For the samples that were eluted a second time (Figure 7A), the next morning the beads were separated from the supernatant and the beads were again incubated with 98 μl of TE/SDS for 2 hours at 65°C before proteinase K treatment. The IPs were performed in duplicate (Figure 7A, 7B: High wash 1, and 7C: 4°C 20 min and 4°C 60 min and 7D: 4°C 20 min and 4°C 60 min), triplicate (Figure 7B: standard wash and 7C: RT 20 min) or quadruplicate (Figure 7B: Low wash and High wash2).

**Figure 8**

*Figure 8A and 8B*

Abf1-aa strains used in Figure 8A and 8B were grown in a big batch, by growing several 250 ml cultures. The cultures were cross-linked with 1% formaldehyde by adding 7.0 ml of 37% formaldehyde and incubating them for 5 minutes. Subsequently, 51.4 ml 4.5 M Tris was added to a final concentration of 750 mM and incubated for 1 minute. The cells were harvested as described in the general methods section.

Abf1-aa cells were bead beated for 7x 3 minutes in a genie disruptor in the presence of protease inhibitors, either by addition of a protease inhibitor tablet per 25 ml FA lysis buffer (Roche #11836145001) or by addition of 30 μl aprotinin (Sigma-Aldrich: #A6279), 1 μl leupeptin (Sigma-Aldrich #L2884, 1 mg/ml in MQ), 1 μl pepstatin (Sigma-Aldrich #P4265: 1 mg/ml in 100% Methanol) and 15 μl PMSF (Sigma-Aldrich: #P7626, 200 mM in isopropanol) per milliliter of FA lysis buffer. A 20 μl sample was taken before lysis for Western blotting (pre-lysis). The cell debris was pelleted and the supernatant was used for sonication. A 40 μl aliquot was taken for Western blotting (post-lysis). The chromatin was fragmented by splitting each sample over 2x 1.5 ml sonication tubes and sonicating for 4 cycles 30 seconds on / 30 seconds off. For the samples with the separate protease inhibitors, 10 μl was added right before the sonication, because PMSF loses activity in aqueous solutions. The sheared chromatin was pooled per cell pellet. A 40 μl aliquot was taken for Western blotting (post-lysis) and 450 μl of chromatin was used for each IP.

The IPs with magnetic beads were performed as described in the general methods section. 50 μl magnetic beads and 1 μl anti-V5 antibody were used for the IPs. The chromatin was incubated with the anti-V5 antibody for 2 hours. To the samples that were prepared with the separate protease inhibitors, again protease inhibitors were added: 15 μl aprotinin, 0.5 μl pepstatin A (1 mg/ml in 100% Methanol), 0.5 μl leupeptin (1 mg/ml in MQ) and 5 μl PMSF (200 mM in isopropanol). 50 μl beads per IP was washed in individual 1.5 ml Eppendorf tubes using 500 μl PBS-T. The beads were resuspended in 400 μl PBS, 25 μl BSA (10 mg/ml in TBS-T) was added and the beads were incubated at 4°C while the chromatin was incubating with the antibody. Just before this incubation was finished, the beads were washed again with 500 μl PBS-T. The chromatin and antibody were added to the beads and incubated at RT for 20 minutes. To the samples that were prepared with the separate protease inhibitors, again 5 μl PMSF was added. The beads were washed with the standard wash of 2x PBS and 2x PBS-T and DNA was recovered by eluting overnight in TE/SDS at 65°C. The IPs were performed in duplicate.

Samples for Western blotting were prepared by mixing 40 μl of the sample with 10 μl of 5X sample buffer (5% SDS, 200 mM Tris pH 6.8, 25% glycerol, 1.43 M β-mercaptoethanol, 0.032% bromophenol blue). Pre-lysis samples were prepared as described in the general methods section, by lysing the cells using NaOH. All samples were heated to 95°C for 30 minutes to reverse cross-links. The samples (10 μl) were separated on a 10% stain-free acrylamide gel (Bio-Rad #1610182). As a control, a non-cross-linked crude lysate of a WT-aa was taken along, which was heated to 95°C for 5 minutes prior to loading. After the proteins had migrated through the gel, the proteins were transferred to a nitrocellulose membrane. The membrane was stained with an anti-V5 antibody (Life Technologies

#R96025) and binding was visualized using a goat-anti-mouse-HRP conjugated antibody (Bio-Rad #1706516) and ECL solution, as described in the general methods section.

*Figure 8C*

Cbf1-aa, Mcm1-aa, Reb1-aa and Sum1-aa strains were grown on a different day than the Abf1-aa strains used in Figure 8A, as 100 ml cultures. The cells were cross-linked by adding 2.8 ml of 37% formaldehyde to a final concentration of 1% and incubating them for 5 minutes. Subsequently, 20.6 ml of 4.5M Tris pH 8.0 was added to a final concentration of 750 mM and the cells were incubated for 1 minute. Cells were harvested as described in the general methods section

The cells were bead beated for 7x 3 minutes in a genie disruptor in the presence of protease inhibitors by addition of 30 µl aprotinin (Sigma-Aldrich: #A6279), 1 µl leupeptin (Sigma-Aldrich #L2884, 1 mg/ml in MQ), 1 µl pepstatin (Sigma-Aldrich #P4265: 1 mg/ml in 100% Methanol) and 15 µl PMSF (Sigma-Aldrich: #P7626, 200 mM in isopropanol) per milliliter of FA lysis buffer. A 20 µl sample was taken before lysis for Western blotting (pre-lysis). The cell debris was pelleted and the supernatant was used for sonication. A 40 µl aliquot was taken for Western blotting (post-lysis). The chromatin was fragmented by splitting each sample over 2x 1.5 ml sonication tubes and sonicating for 4 cycles 30 seconds on / 30 seconds off. 5 µl PMSF was again added right before the sonication, because PMSF loses activity in aqueous solutions. The sheared chromatin was pooled per cell pellet. A 40 µl aliquot was taken for Western blotting (post-lysis) and 450 µl of chromatin was used for each IP.

The samples for Western blotting were prepared by mixing 40 µl of the sample with 10 µl of 5X sample buffer (5% SDS, 200 mM Tris pH 6.8, 25% glycerol, 1.43 M β-mercaptoethanol, 0.032% bromophenol blue). The pre-lysis samples were prepared as described in the general methods section, by lysing the cells using NaOH. All samples were heated to 95°C for 30 minutes to reverse cross-links. The samples (10 µl) were separated on a 10% stain-free acrylamide gel (Bio-Rad #1610182). As a control, a non-cross-linked crude lysate of a WT-aa was taken along, which was heated to 95°C for 5 minutes prior to loading. After the proteins had migrated through the gel, the proteins were transferred to a nitrocellulose membrane. The membrane was stained with an anti-V5 antibody (Life Technologies #R96025) and binding was visualized using a goat-anti-mouse-HRP conjugated antibody (Bio-Rad #1706516) and ECL solution, as described in the general methods section.

**Figure 9**

Abf1-aa and Reb1-aa strains were grown as 100 ml cultures. The cells were cross-linked for 5 minutes using a final concentration of 1%, 2% or 3% formaldehyde, by adding 2.8 ml, 5.7 ml or 8.8 ml 37% formaldehyde, respectively. The samples were subsequently quenched with a final concentration of 750 mM (1% formaldehyde), 1.5M (2% and 3% formaldehyde) or 2.0M (3% formaldehyde) by adding 20.6 ml (1% formaldhyde, 750 mM Tris), 52.9 (2% formaldehyde, 1.5 mM Tris), 54.4 (3% formaldehyde, 1.5 mM Tris) or 87.1 ml (3% formaldehyde, 2.0 mM Tris) 4.5M Tris pH 8.0 and incubating the samples for 1 minute. The cells were harvested as described in the general methods section

The cells were bead beated for 7x 3 minutes in a genie disruptor in the presence of protease inhibitors by addition of 30 µl aprotinin (Sigma-Aldrich: #A6279), 1 µl leupeptin (Sigma-Aldrich #L2884, 1 mg/ml in MQ), 1 µl pepstatin (Sigma-Aldrich #P4265: 1 mg/ml in 100% Methanol) and 15 µl PMSF (Sigma-Aldrich: #P7626, 200 mM in isopropanol) per milliliter of FA lysis buffer. A 5 µl sample was taken before lysis for Western blotting (pre-lysis). The cell debris was pelleted and the supernatant was used for sonication. A 10 µl aliquot was taken for Western blotting (post-lysis). The chromatin was fragmented by splitting each sample over 2x 1.5 ml sonication tubes and sonicating for 10 cycles 30 seconds on / 15 seconds off. 5 µl of PMSF was again added right before the sonication, because PMSF loses activity in aqueous solutions. The sheared chromatin was pooled per cell pellet. A 10 µl aliquot was taken for Western blotting (post-lysis) and 450 µl of chromatin was used for each IP.

The IPs with magnetic beads were performed as described in the general methods section. 25 µl magnetic beads and 1 µl anti-V5 antibody were used for the IPs. The chromatin was incubated with the anti-V5 antibody for 2 hours. Before the incubation, again protease inhibitors were added: 0.5 µl pepstatin, 0.5 µl leupeptin, 15 µl aprotinin and 5 µl PMSF. 25 µl beads per IP was washed in individual 1.5 ml Eppendorf tubes using 500 µl PBS-T. The beads were resuspended in 400 µl PBS, 25 µl BSA (10 mg/ml in TBS-T) was added and the beads were incubated at 4°C while the chromatin was incubating with the antibody. Just before this incubation was finished, the beads were washed again with 500 µl PBS-T. The chromatin and antibody were added to the beads and incubated at RT for 20 minutes. Right before the incubation of the chromatin and antibody with the beads, again 5 µl PMSF was added. The beads were washed with the standard wash of 2x PBS and 2x PBS-T and DNA was recovered by eluting overnight in TE/SDS at 65°C. The IPs were performed without replicates.

The samples for Western blotting were prepared by mixing 10 µl of the sample with 2.5 µl of 5X sample buffer (5% SDS, 200 mM Tris pH 6.8, 25% glycerol, 1.43 M β-mercaptoethanol, 0.032% bromophenol blue). The pre-lysis samples were prepared as described in the general methods section, by lysing the cells using NaOH. All samples were heated to 95°C for 30 minutes to reverse cross-links. The samples (10 µl) were separated on a 10% stain-free acrylamide gel (Bio-Rad #1610182). As a control, a non-cross-linked crude lysate of a WT-aa was taken along, which was heated to 95°C for 5 minutes prior to loading. After the proteins had migrated through the gel, the proteins were transferred to a nitrocellulose membrane. The membrane was stained with an anti-V5 antibody (Life Technologies #R96025) and binding was visualized using a goat-anti-mouse-HRP conjugated antibody (Bio-Rad #1706516) and ECL solution, as described in the general methods section.

**Table S1 List of genotypes of strains used in this study**

| Strain | Genotype |
|---|---|
| WT-aa | tor1-1; Δfpr1; RPL13-2xFKBP12-NATMX6; MET15; his3-1; leu2; lys2; ura3; MATalpha |
| Cbf1-aa | tor1-1; Δfpr1; RPL13-2xFKBP12-NATMX6; MET15; his3-1; leu2; lys2; ura3; CBF1-FRB-yEGFP-3V5-hphMX6; MATalpha |
| Cbf1-aa Cha4-V5 | tor1-1; Δfpr1; RPL13-2xFKBP12-NATMX6; MET15; his3-1; leu2; lys2; ura3; CHA4-3V5-bleMX6; CBF1-FRB-yEGFP-3V5-hphMX6 MATalpha |
| Abf1-aa | tor1-1; Δfpr1; RPL13-2xFKBP12-NATMX6; MET15; his3-1; leu2; lys2; ura3; ABF1-FRB-yEGFP-3V5-hphMX6; MATalpha |
| Reb1-aa | tor1-1; Δfpr1; RPL13-2xFKBP12-NATMX6; MET15; his3-1; leu2; lys2; ura3; REB1-FRB-yEGFP-3V5-hphMX6; MATalpha |
| Mcm1-aa | tor1-1; Δfpr1; RPL13-2xFKBP12-NATMX6; MET15; his3-1; leu2; lys2; ura3; MCM1-FRB-yEGFP-3V5-hphMX6; MATalpha |
| Sum1-aa | tor1-1; Δfpr1; RPL13-2xFKBP12-NATMX6; MET15; his3-1; leu2; lys2; ura3; SUM1-FRB-yEGFP-3V5-hphMX6; MATalpha |

**Table S2 list of primers used in this study**

| Name | Sequence (5′ -> 3′ ) | Target | Figure |
|---|---|---|---|
| ACT1_nucl-F | ATATGTTTAGAGGTTGCTGCTTTG | Background | 2A-2D |
| ACT1_nucl-R | AACCGGCTTTACACATACCA | | |
| BNA2_qP_F | TTTCTCTATGGGCTGACG | Sum1 | 7 |
| BNA2_qP_R | ATGCTAAAGATACATGGACATC | | |
| CFD1-F | CGGGATCTTTGGTTCCTATC | Background | 2E |
| CFD1-R | AGCCTTCCGATTTCTTTCC | | |
| CHA1-F | GGGCGGCTCCTGTTAAG | Cha4 | 3, 4B, 5 |
| CHA1-R | TCCTCCTCATATTGTCCCTTT | | |
| CPA2_TF_F | CACAATCGTTACGACATGGAG | Background | 2E, 3, 4B, 5A |
| CPA2_TF_R | GACTCTTATTGATGAGATGGCAATA | | |
| EMC6_qP_F | TACGGTCACGCCAATTC | Reb1 | 9B |
| EMC6_qP_R | GCTTGAGCAATCCAACATAAG | | |
| FCF1_qP_F | ACTATCGGTTCCTACTGGAAGA | Abf1 | 8, 9A |
| FCF1_qP_R | GGCATCATTCAAGAATAGTAGCAAG | | |
| GLN1_qP_F | TACCCGCATACGGTTCT | Reb1 | 9B |
| GLN1_qP_R | GGAGCGCAGTCATCAAT | | |
| QCR10-F | TAACGCTGTCGCACTTTGA | Cbf1 | 2A-2D, 3-6 |
| QCR10-R | GAAACAACGGGTTGAACCATATT | | |
| NHX1_qP_F | CACAAACGTGATAGCAAGGAAC | Abf1 | 8, 9A |
| NHX1_qP_R | TCTTGGGTGCCCTTTATCTTAG | | |
| TFC1_pr_qP_F | TCTTTAAGCTCTGCTGTGTT | Background | 6-9 |
| TFC1_pr_qP_R | AGCGAAGAAGCGAAGAAG | | |
| TNA1_qP_F | ACTCTCCAAGCTATAAGCATAC | Sum1 | 7 |
| TNA1_qP_R | TTTCAGTCGCTGTCTCAC | | |
| TUB1_pr_F | TACAGATCTTGGGTGGCGAGAACT | Background | 2A-2D, 4A, 5B, 6-8 |
| TUB1_pr_R | AAACGCCTCGAGCCAAGGGAAA | | |
| YOS1-F | GCTCGTAATGTCTCGAAATTTGTC | Cbf1 | 2-6 |
| YOS1-R | CTTATTGAGAAGGCTCCCAGTC | | |

**Table 3 list of buffers used in this study**

| Solution name | Components |
|---|---|
| Formaldehyde | 37% formaldehyde containing 10-15% methanol (Sigma-Aldrich #252549) |
| Glycine | 2.5 M Glycine |
| Tris | 4.5 M Tris pH 8.0 |
| Buffer Z | 1 M sorbitol<br>50 mM Tris pH 7.5 |
| Zymolyase solution | Buffer Z<br>10 mM β-mercaptoethanol<br>10 mg/ml zymolyase (zymolyase 20 T MP biomedical #08320921) |
| TBS | 150 mM NaCl<br>10 mM Tris pH 7.5 |
| FA lysis buffer | 50 mM HEPES-KOH pH 7.5<br>150 mM NaCl<br>1 mM EDTA pH 8.0<br>1% Triton X-100<br>0.1% Na-deoxycholate<br>0.1% SDS |
| Aprotinin | 5 - 10 TIU /ml (Sigma-Aldrich: #A6279), |
| Pepstatin A | 1 mg/ml leupeptin in 100% methanol (1.51 mM) (Sigma-Aldrich #L2884) |
| Leupeptin | 1 mg/ml pepstatin A in MQ (2.10 mM) (Sigma-Aldrich #P4265) |
| PMSF | 200 mM Phenylmethanesulfonyl fluoride (PMSF) in isopropanol (Sigma #P7626) |
| RNAse A/T1 | RNAse A 2mg/ml & RNase T1 5000 U/ml mix (Thermo Scientific #EN0551) |
| Proteinase K | 10 μg/μl in TE (10 mM Tris pH 8, 1 mM EDTA pH 8) |
| PBS | 137 mM NaCl<br>2.7 mM KCl<br>10 mM $Na_2HPO_4$<br>1.47 mM $KH_2PO_4$<br>1 mM CaCl<br>0.5 mM MgCl<br>pH adjusted to 7.3 using HCl |
| PBS-T | PBS + 0.02% Tween-20 |
| Wash buffer 1 | FA lysis buffer<br>500 mM NaCl (final concentration) |
| Wash buffer 2 | 10 mM Tris pH 8.0<br>0.25 mM LiCl<br>1 mM EDTA pH 8.0<br>0.5% Nonidet P-40<br>0.5% Na-deoxycholate |
| TE/SDS | 10 mM Tris pH 8.0<br>1 mM EDTA pH 8.0<br>1% SDS |
| BSA | 10 mg/ml in TBS-T (150 mM NaCl, 10 mM Tris pH 7.5, 0.05% Tween-20) |
| 2X sample buffer | 2% SDS<br>80 mM Tris pH 6.8<br>10 % glycerol<br>572 mM β-mercaptoethanol<br>0.016% bromophenol blue |
| 5X sample buffer | 5% SDS |

| | |
|---|---|
| | 200 mM Tris pH 6.8<br>25% glycerol<br>1.43 M β-mercaptoethanol<br>0.032% bromophenol blue |
